# scDesign2: a transparent simulator that generates high-fidelity single-cell gene expression count data with gene correlations captured

**DOI:** 10.1101/2020.11.17.387795

**Authors:** Tianyi Sun, Dongyuan Song, Wei Vivian Li, Jingyi Jessica Li

## Abstract

In the burgeoning field of single-cell transcriptomics, a pressing challenge is to benchmark various experimental protocols and numerous computational methods in an unbiased manner. Although dozens of simulators have been developed for single-cell RNA-seq (scRNA-seq) data, they lack the capacity to simultaneously achieve all the three goals: preserving genes, capturing gene correlations, and generating any number of cells with varying sequencing depths. To fill in this gap, here we propose scDesign2, a transparent simulator that achieves all the three goals and generates high-fidelity synthetic data for multiple scRNA-seq protocols and other single-cell gene expression count-based technologies. Compared with existing simulators, scDesign2 is advantageous in its transparent use of probabilistic models and is unique in its ability to capture gene correlations via copula. We verify that scDesign2 generates more realistic synthetic data for four scRNA-seq protocols (10x Genomics, CEL-Seq2, Fluidigm C1, and Smart-Seq2) and two single-cell spatial transcriptomics protocols (MERFISH and pciSeq) than existing simulators do. Under two typical computational tasks, cell clustering and rare cell type detection, we demonstrate that scDesign2 provides informative guidance on deciding the optimal sequencing depth and cell number in single-cell RNA-seq experimental design, and that scDesign2 can effectively benchmark computational methods under varying sequencing depths and cell numbers. With these advantages, scDesign2 is a powerful tool for single-cell researchers to design experiments, develop computational methods, and choose appropriate methods for specific data analysis needs.

## Introduction

The recent development of the single-cell RNA-seq (scRNA-seq) technologies has revolutionized transcriptomic studies by revealing the genome-wide gene expression levels within individual cells [1, 2]. In contrast to bulk RNA sequencing, scRNA-seq technology captures cell-specific transcriptome landscapes, which can reveal crucial information about cell-to-cell heterogeneity across different tissues, organs, and systems and enable the discovery of novel cell types and new transient cell states [3–8]. Already, scRNA-seq technologies have led to breakthroughs in understanding biological processes such as stem cell differentiation and embryogenesis [9, 10], neurological disorders [11, 12], and tumorigenesis [13, 14].

Since the first scRNA-seq study was published in 2009 [15], many experimental protocols have been developed [16–18]. Broadly speaking, the existing protocols fall into two categories: tag-based and full-length [19]. Tag-based protocols (e.g., 10x Genomics [20], CEL-Seq2 [21], Drop-seq [22], and Seq-Well [23]) only capture and sequence one end of RNA transcripts, while full-length protocols (e.g., Smart-Seq2 [24], Fluidigm C1 [25], and MATQ-seq [26]) sequence fragments from full-length RNA transcripts [18, 27]. Typically, compared to full-length protocols (given the sequencing depth), tag-based protocols sequence more cells but with fewer transcripts captured per cell [28]. In addition to this cell-number vs. per-cell-depth trade-off, tag-based protocols use unique molecular identifiers (UMIs) to remove polymerase chain reaction (PCR) amplification biases [29], while full-length protocols do not have this advantage and can only output reads without UMIs. Therefore, these protocols have different advantages in throughput (number of cells and number of genes captured) and accuracy (number of non-biological zeros and PCR biases) [18, 30, 31]. Moreover, when designing experiments, researchers often face the practical issue of having a limited budget. In this case, they need guidance to choose either sequencing more cells with fewer reads (or UMIs) in each cell, or sequencing fewer cells with more reads (or UMIs) in each cell [32–34].

In addition to selecting experimental protocols before conducting scRNA-seq experiments, a common challenge after collecting scRNA-seq data is to choose among the many available data analysis methods in an unbiased manner. For example, many algorithms have been developed for missing gene expression imputation [35, 36], dimensionality reduction [37–39], cell clustering [40–43], rare cell type detection [44–46], differentially expressed gene identification [47–49], and trajectory inference [50–54]. Even though several benchmark and comparative studies have been carried out for common analysis tasks [55–59], most of them have only evaluated a subset of available computational methods using data from limited experimental protocols. Hence, they cannot meet the diverse needs of ongoing and future analyses of scRNA-seq data. In short, single-cell researchers lack a systematic and flexible approach to select appropriate computational methods for their specific data analysis needs.

One solution to the above two issues is to use *in silico* synthetic datasets, which carry ground truths (cell types, cell trajectories, differentially expressed genes, etc.) and do not induce extra experimental costs. Below we summarize six properties that an ideal simulator should achieve.

1. The simulator can be trained by real data so that it is adaptive to various experimental protocols and biological conditions.
2. The simulator can preserve genes so that its synthetic cells contain expression levels of real genes. The simulator should retain every gene’s distribution of expression levels in its synthetic data without deleting genes in real data. This property is essential for benchmarking differential gene expression analysis.
3. The simulator can capture gene correlations so that its synthetic data maintain a similar gene correlation structure to that in real data. This property relies on the last property and is essential for benchmarking multi-gene analyses such as cell dimensionality reduction (e.g., principal component analysis (PCA), t-distributed stochastic neighbor embedding (t-SNE) [60, 61], and uniform manifold approximation and projection (UMAP) [62, 63]), cell clustering, rare cell type detection, and cell trajectory inference.
4. The simulator can generate synthetic data with both varying cell number and sequencing depth, under the same biological condition of training data. This property is essential for guiding experimental design and benchmarking robustness of computational methods.
5. The simulator is transparent so that its model parameters can be easily understood and adjusted. For example, key statistical properties, such as every gene’ expression mean, variance, zero proportions, and every gene pair’s expression correlation, can be easily accessed from the model. This property is essential for model diagnostics and customized simulation. Specifically, with a transparent model, whenever the synthetic data do not resemble the real data, computational researchers can easily access how well the model fits to each gene’s marginal distribution and what genes’ correlations are well captured or missed. Moreover, a transparent model offers users an opportunity to generate data from their specified parameter values, e.g., gene expression means.
6. The simulator is computationally and sample efficient so that its training does not require expensive hardware, take excessive computational time, or rely on an enormous number of real cells to achieve good training. This property is essential for the simulator to be userfriendly and adaptive to full-length protocols that generate hundreds to thousands of cells, e.g., Fluidigm C1 and Smart-Seq2.

Although many simulators have been developed for scRNA-seq data and various methodological advances have been made [34, 38, 64–72], to the best of our knowledge, none of them achieves all the six properties. We summarize 14 representative simulators in Table 1. Except scGAN [68], these simulators all use probabilistic models or differential equations that are transparent and easy to fit, thus satisfying properties 5 and 6. However, scDesign [34], three simulators in the splatter package (splat simple, splat, and kersplat) [65], and SymSim [66] do not preserve genes, failing properties 2 and 3; ZINB-WaVE^1^ [38, 65] and SPARSim [67] cannot vary cell number or sequencing depth, failing property 4; SERGIO [72] requires a user-specified gene regulatory network as input and does not estimate gene correlations from real data^2^, thus not achieving property 3. Although scGAN preserves genes and uses a deep neural network to capture gene correlations, it cannot vary sequencing depth (not satisfying property 4), and its black-box nature, requirement for GPU, and long computational time make it fail properties 5 and 6. Hence, a simulator that achieves all the six properties is in demand.

**Table 1:**
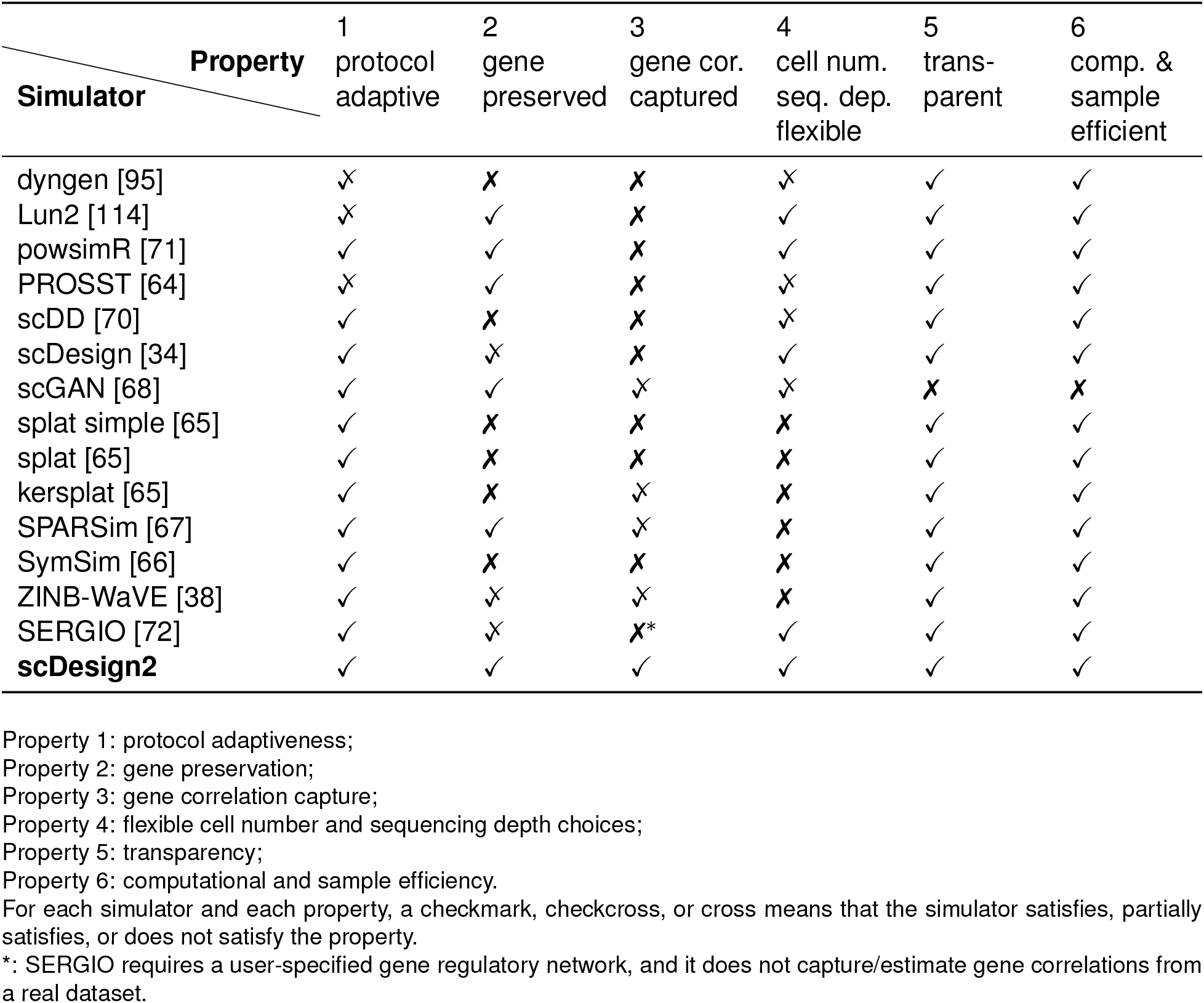
Summary of 14 simulators (including our proposed scDesign2) in six properties.

Here we propose scDesign2 as the first simulator that achieves all the six properties and generates realistic synthetic data for multiple single-cell gene expression count-based technologies. Inheriting its name from our previous simulator scDesign, scDesign2 has achieved a significant methodological advance and become the first transparent simulator that reliably captures gene correlations. This advance is enabled by probabilistic modeling of not only marginal distributions of individual genes but also the joint distribution of thousands of genes. Thanks to its achievement of the six properties, scDesign2 will serve as a powerful tool for guiding experimental design and benchmarking computational methods in the single-cell transcriptomics field.

## Results

### An overview of scDesign2

The statistical framework of scDesign2 consists of two steps: (1) model-fitting and (2) synthetic data generation (Fig. 1). In the model-fitting step, scDesign2 fits a multivariate generative model to a real scRNA-seq dataset. If the dataset contains more than one cell type (defined by marker genes or cell clustering; see Methods), then scDesign2 divides the dataset into subsets, one per cell type, and fits a cell-type-specific model to each subset. In the data-generation step, scDesign2 generates synthetic scRNA-seq data from the fitted model for each cell type.

**Figure 1:**
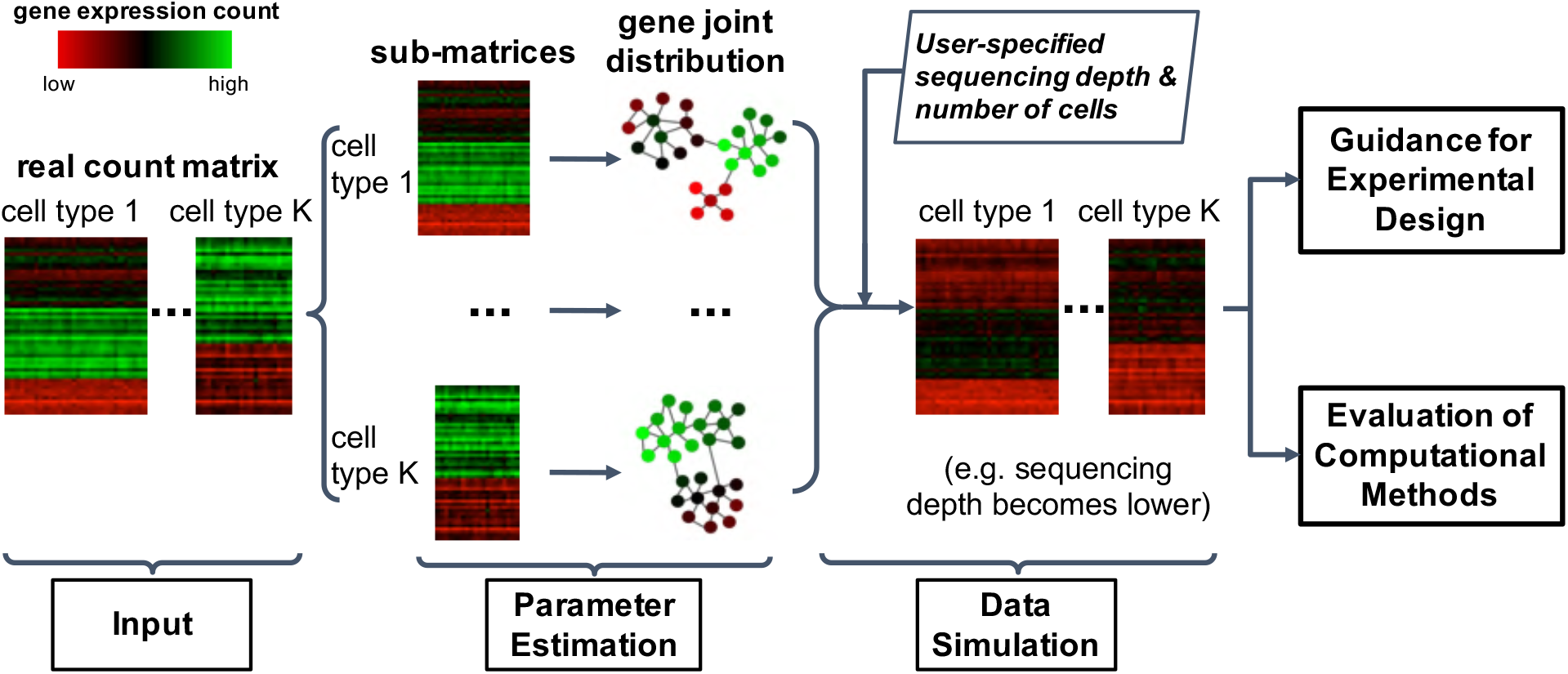
An overview of scDesign2. The input for scDesign2 is a gene-by-cell count matrix with cells labelled as cell types or clusters. For cells in each type or cluster, scDesign2 uses the copula framework to fit a joint distribution of gene expression counts. Then given user-specified sequencing depth and number of cells, scDesign2 generates synthetic data for each cell type or cluster. The synthetic data can be used to guide experimental design and evaluate computational methods.

The model-fitting step is composed of the following two sub-steps. First, scDesign2 fits a univariate count distribution to each gene’s counts in cells of the same type. Four count distributions are considered: Poisson, zero-inflated Poisson (ZIP), negative binomial (NB), and zero-inflated negative binomial (ZINB), with the former three as special cases of the ZINB. All the four distributions have been widely used to model a gene’s read or UMI counts in a homogeneous group of cells [38, 73–75]. From these four distributions, scDesign2 chooses one distribution for every gene in every cell type in a data-driven way. Second, scDesign2 captures the correlations of thousands of genes (all the moderately to highly expressed genes) by fitting a Gaussian copula in each cell type. We choose the Gaussian copula for its easiness to fit and good transparency, and we find it capturing gene correlations well (Fig. 2 and Supplementary Figs. S1–S5).

**Figure 2:**
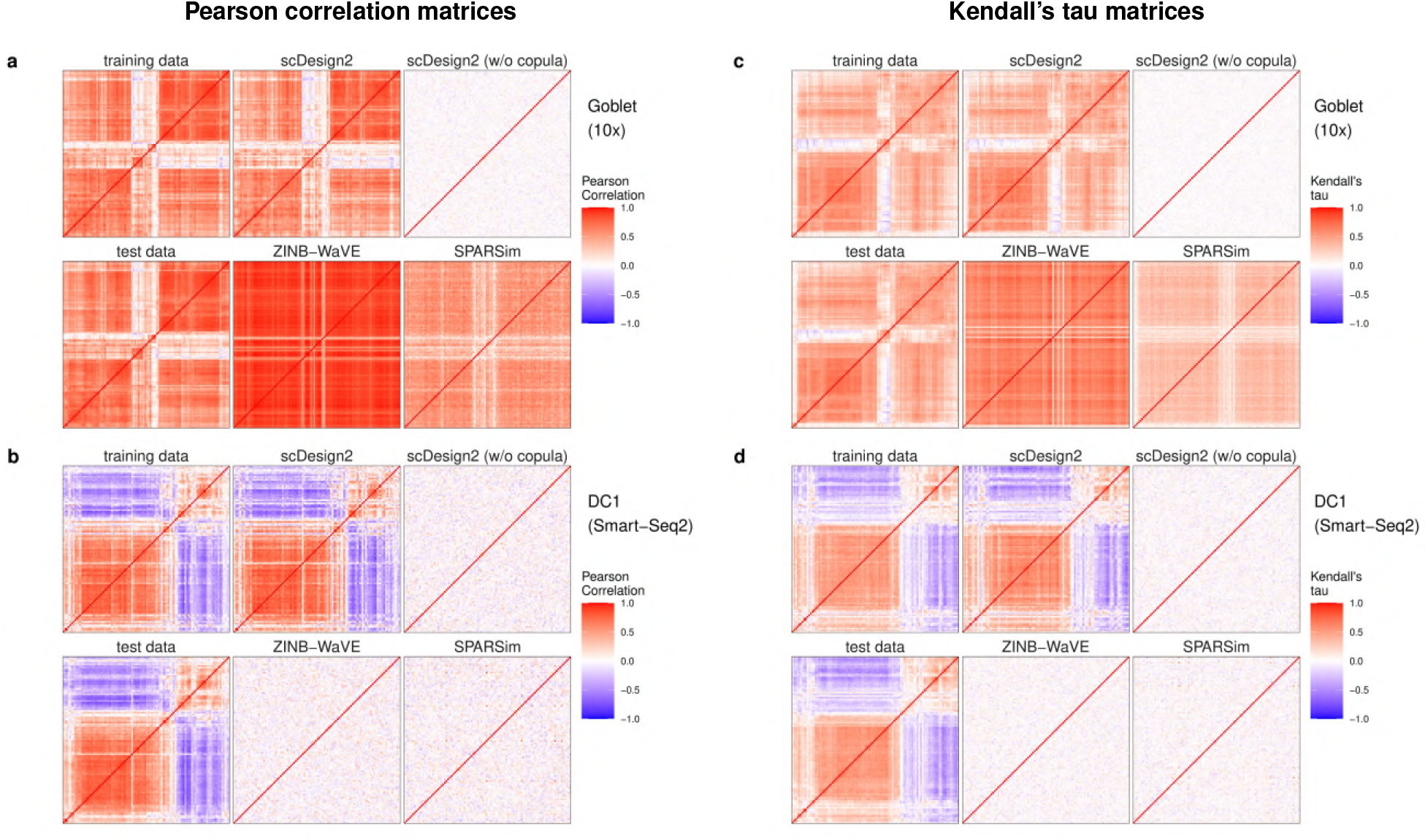
Heatmaps of gene correlation matrices estimated from real data and synthetic data generated by scDesign2, its variant without copula, ZINB-WaVE, and SPARSim. (a)-(b) Pearson correlation matrices; (c)-(d) Kendall’s tau matrices. In (a) and (c), training and test data contain goblet cells measured by 10x Genomics [76]; In (b) and (d), training and test data contain cells of dendrocytes subtype 1 (DC1) measured by Smart-Seq2 [79]. For each cell type, the Pearson correlation matrices and Kendall’s tau matrices are shown for the 100 genes with the highest mean expression values in the test data; the rows and columns (i.e., genes) of all the matrices are ordered by the complete-linkage hierarchical clustering of genes (using Pearson correlation as the similarity in (a)-(b) and Kendall’s tau in (c)-(d)) in the test data. We find that the correlation matrices estimated from the sythetic data generated by scDesign2 most resemble those of training and test data.

As the first simulator that explicitly captures gene correlations, scDesign2 leverages a unique advantage of the copula framework: the separate modeling of each gene’s marginal distribution and the correlation structure of thousands of genes together. This separation and its resulting flexibility are critical for scDesign2 to model single-cell gene expression count data generated by various experimental protocols. Thanks to this flexibility, scDesign2 can choose a count distribution from Poisson, ZIP, NB, and ZINB to fit each gene’s expression counts and reveal biological insights of that gene’s expression pattern.

### Synthetic data generated by scDesign2 most resemble real scRNA-seq data in benchmarking against existing simulators

We benchmark scDesign2 against eight existing simulators—ZINB-WaVE, SPARSim, scGAN, scDesign, three variants of splat in the splatter package (splat simple, splat, and kersplat), and SymSim. We also compare scDesign2 with its own variant that only uses gene marginal distributions and no copula (w/o copula). Among these ten simulators, only scDesign2, its w/o copula variant, ZINB-WaVE, SPARSim, and scGAN preserve genes. We apply these ten simulators to four scRNA-seq datasets (in which cells are labelled with curated cell types) generated by different experimental protocols (10x Genomics [76], CEL-Seq2 [77], Fluidigm C1 [78], and Smart-Seq2 [79]). For each dataset, we randomly split its cells into two halves, with one half (“training data”) to be used for training every simulator on each cell type individually and the other half (“test data”) to serve as the benchmark standard to be compared with the synthetic data generated by each simulator.

We use three sets of benchmark analyses to compare synthetic data with the corresponding test data. Here is an overview. First, we select three cell types from each dataset (measured by each experimental protocol), obtaining a total of 12 cell-type–protocol combinations. For each combination, we evaluate eight key statistics: four gene-wise (expression mean, variance, coefficient of variation (cv), and zero proportion); two cell-wise (zero proportion and library size); two gene-pair-wise (Pearson correlation and Kendall’s tau). (Note that we include Kendall’s tau instead of Spearman rank correlation as a rank-based correlation statistic because Kendall’s tau can account for ties.) For each statistic, we compare its empirical distribution—across genes (for gene-wise statistics), across cells (for cell-wise statistics), or across gene-pairs (for gene-pair-wise statistics)—in the test data with that in the synthetic data generated by each simulator. For the four gene-wise and two gene-pair-wise statistics, we also directly compare their values in the test data with those in the synthetic data generated by scDesign2, ZINB-WaVE, SPARSim, and scGAN— the four simulators that preserve genes. We cannot do this for the other simulators, because the values of these gene-related statistics are not comparable if the genes are not preserved. The results are summarized in Fig. 3 and Supplementary Figs. S6–S28. Second, for each of the 12 cell-type–protocol combinations, we compare the gene correlation matrix estimated from the test data with that from the synthetic data generated by each simulator that preserves genes. We exclude the simulators that do not preserve genes because the gene expression matrices estimated from their synthetic data do not align with those from real data (i.e., the genes of the synthetic data matrix cannot be matched one-to-one to the genes of the training data matrix). The results are summarized in Fig. 2 and Supplementary Figs. S1–S5. Third, for each of the four protocols, we use 2D visualization—t-SNE and PCA—to compare cells of multiple types in the test data and the synthetic data generated by each simulator that preserves genes. Again, we exclude the simulators that do not preserve genes because their synthetic cells cannot be combined with real cells for joint visualization (dimesionality reduction requires all cells to have the same original dimensions, i.e., genes). The results are summarized in Fig. 4 and Supplementary Figs. S29–S31.

**Figure 3:**
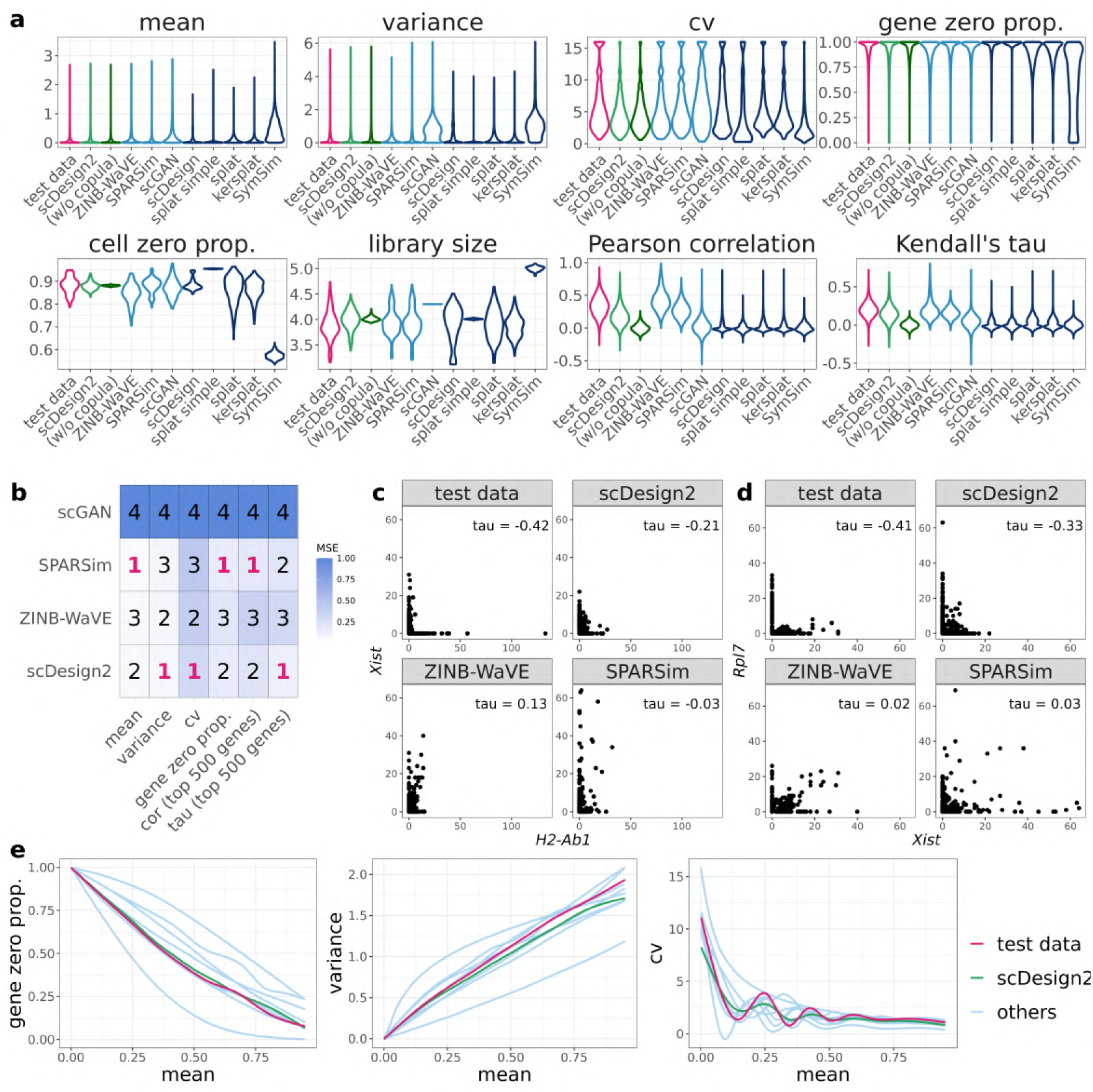
Benchmarking scDesign2 against its variant without copula and eight existing scRNA-seq simulators for generating goblet cells measured by 10x Genomics. (a) Distributions of eight summary statistics (gene-wise expression mean, variance, coefficient of variation (cv), and zero proportion; cell-wise zero proportion and library size; gene-pair-wise Pearson correlation and Kendall’s tau) are plotted based on the real data (test data unused for training simulators) and the synthetic data generated by scDesign2, scDesign2 without copula (w/o copula), ZINB-WaVE, SPARSim, scGAN, scDesign, three variants of the splatter package (splat simple, splat, and kersplat), and SymSim. (b) Ranking (with 1 being the best-performing method) of scDesign2, ZINB-WaVE, SPARSim, and scGAN, the only four methods that preserve genes, in terms of the mean-squared error (MSE) of each of six summary statistics (four gene-wise and two gene-pair-wise) between the statistic values in the real data and the synthetic data generated by each simulator. Note that the color scale shows the normalized MSE: for each statistic (column in the table), the normalized MSEs are the MSEs divided by the largest MSE of that statistic. scDesign is ranked the top for three out of the six statistics. For the two gene-pair-wise statistics, we focus on the top 500 highly expressed genes, because as analyzed in the text, they are more meaningful, both biologically and statistically, than the correlations of the lowly expressed genes. (c)-(d) Scatterplots of two example gene pairs—*Xist* vs. *H2-Ab1* and *Rpl7* vs. *Xist*—based on the real data and the synthetic data generated by scDesign2, ZINB-WaVE, and SPARSim. The Kendall’s tau values in the synthetic data generated by scDesign2 resemble most the values in the test data. (e) Smoothed relationships between three pairs of gene-wise statistics (zero proportion vs. mean, variance vs. mean, and cv vs. mean) across all genes (curves plotted by the R function `geom_smooth()`) in the real data and the synthetic data generated by scDesign2 and the eight existing simulators (others). Note that ZINB-WaVE and SymSim filter out certain genes when simulating new data; Pearson correlation and Kendall’s tau are only calculated between the genes whose zero proportions are less than 50%; gene-wise mean and variance and cell-wise library size are transformed to the log_10_(l + *x*) scale (where *x* represents a statistic’s value).

**Figure 4:**
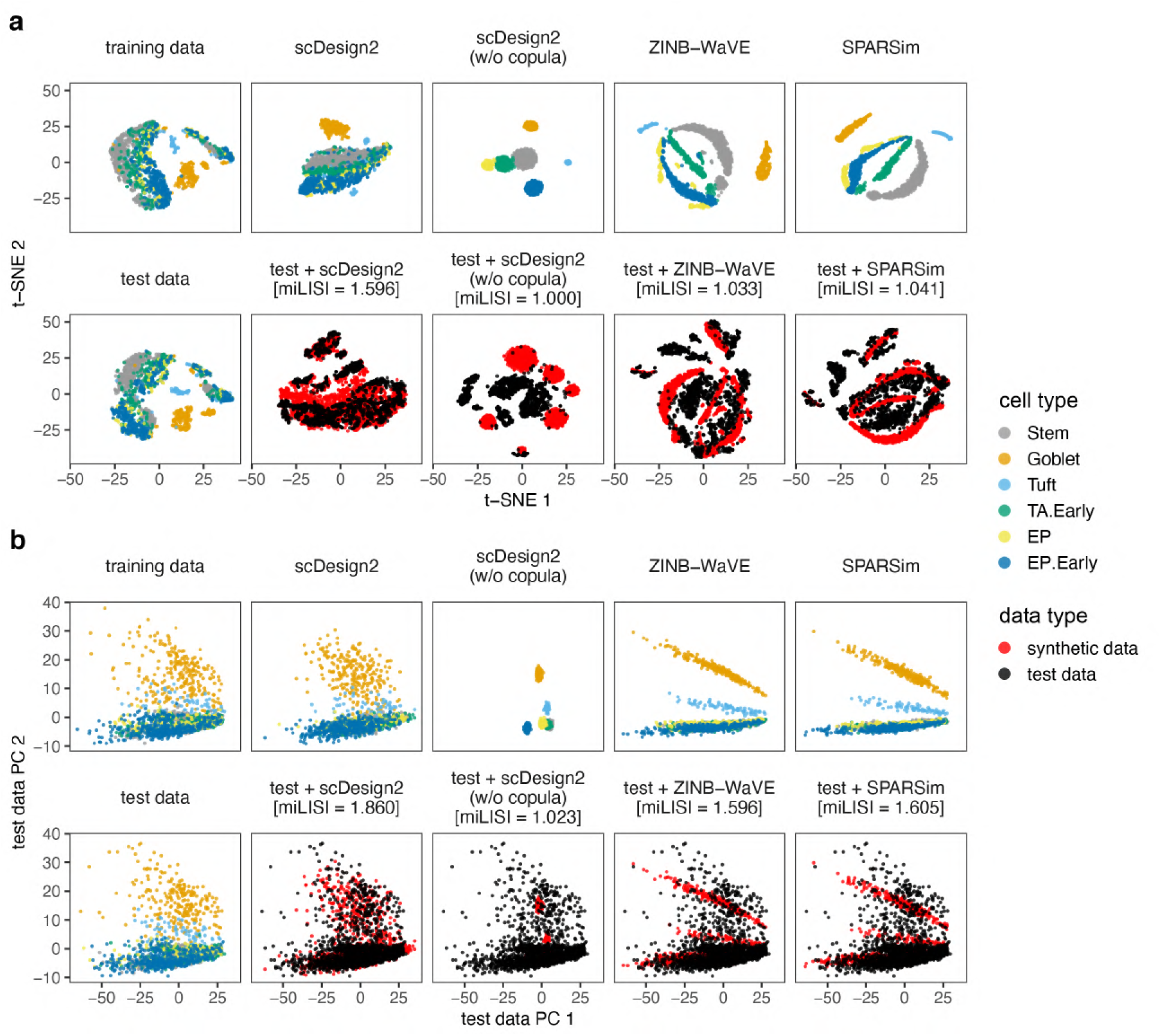
Comparison of 10x Genomics data and synthetic data generated by scDesign2, its variant without copula, ZINB-WaVE, and SPARSim in 2D visualization. (a) t-SNE plots and (b) principle component (PC) plots of training data, test data, synthetic data generated by each simulator, and combinations of test data and each synthetic dataset. Gene expression counts are transformed as log(l + count) before dimensionality reduction. miLISI is short for median integration local inverse Simpson’s Index, a higher value of which indicates that the simulated data mix better with the test data in the 2D visualization plot. By visually inspecting the patterns in these plots as well as comparing the miLISI values, we find that the synthetic data generated by scDesign2 most resemble the test data.

Overall, we find that the synthetic data generated by scDesign2 most resemble the test data for all four protocols. In our first set of analyses, we categorize the eight existing simulators into two types: simulators that preserve genes (ZINB-WaVE, SPARSim, and scGAN) and others. First, by comparing the distributions of eight key statistics between test data and synthetic data, we find that the simulators capable of preserving genes have overall better performance than other simulators, across cell types and protocols (Fig. 3a and Supplementary Figs. S6a–S16a).

Second, we further benchmark the gene-preserving simulators by directly comparing their synthetic data and test data in terms of the gene-wise and gene-pair-wise statistics’ values. Note that we cannot compare these statistics’ values for simulators that do not preserve genes because the “genes” in those simulators’ synthetic data cannot be matched to any genes in real data. In detail, we calculate the mean-squared errors (MSEs) of the four gene-wise statistics and the two gene-pair-wise statistics between test data and synthetic data generated by scDesign2, ZINB-WaVE, SPARSim, and scGAN. Fig. 3b shows that scGAN, a deep-learning-based method, consistently has the worst MSEs for all the six statistics. Due to its long computational time^3^, difficult implementation, and unsatisfactory performance, we exclude it from the following comparisons.

Out of 48 comparisons of gene-wise statistics (4 statistics times 12 cell-type–protocol combinations), scDesign2 achieves the best MSEs in 37 comparisons and demonstrates a clear advantage over ZINB-WaVE and SPARSim (Fig. 3b and Supplementary Figs. S6b–S16b). Out of 24 comparisons of gene-pair-wise statistics (2 correlation statistics times 12 cell-type–protocol combinations) based on the 500 most highly expressed genes (in terms of their mean expression levels across cells) in each cell-type–protocol combination, scDesign2 achieves the best MSEs in 15 comparisons (Fig. 3b and Supplementary Figs. S6b–S16b). We highlight the highly expressed genes because their Pearson correlations and Kendall’s tau values are more biologically meaningful; in all cell-type–protocol combinations, the top 500 highly expressed genes, ranked by either mean expression levels or non-zero proportions across cells, explain at least 50% of reads or UMIs (Supplementary Figs. S17c–S28c), confirming that these genes play dominant roles in transcriptional programs in cells. In addition, we include the comparison results based on more genes in Supplementary Figs. S17d&e–S28d&e, which show that, as more lowly expressed genes are included, the MSEs of all these simulators decrease and become less distinguishable (because lowly expressed gene pairs have correlations close to zero in test data and all synthetic data), making the comparison less meaningful.

Third, we examine correlations of individual gene pairs and observe that scDesign2 can preserve strong negative gene correlations missed by ZINB-WaVE and SPARSim, which wrongly capture these correlations as weak or even positive (Fig. 3c-d and Supplementary Figs. S6c-d– S16c-d). This observation is further confirmed by our second set of analyses below. Furthermore, we compare the relationships of three pairs of gene-wise statistics (zero proportion vs. mean, variance vs. mean, and cv vs. mean) between test data and synthetic data generated by each simulator, and we find that scDesign2 better captures the relationships than existing simulators do across cell types and experimental protocols (Fig. 3e and Supplementary Figs. S6e–S16e).

In our second set of analyses, we compare gene correlation matrices in terms of both Pearson correlation and Kendall’s tau between test data and synthetic data generated by scDesign2, ZINB-WaVE, and SPARSim. Heatmap visualization shows that scDesign2 captures gene correlations most accurately and consistently across cell types and experimental protocols (Fig. 2 and Supplementary Figs. S1–S5). Notably, for highly expressed genes in Smart-Seq2 data, ZINB-WaVE and SPARSim miss almost all the gene correlations, while scDesign2 well preserves positive and negative gene correlations in its synthetic data (Fig. 2b & d and Fig. S5b & d).

In our third set of analyses, we use 2D visualization to compare cells in test data and those in synthetic data generated by scDesign2, ZINB-WaVE, and SPARSim. Both t-SNE and PCA 2D plots show that cells in synthetic data generated by scDesign2 most resemble cells in test data (Fig. 4 and Supplementary Figs. S29–S31). In particular, by overlaying real and synthetic cells in the same 2D plot, we find synthetic cells generated by scDesign2 least distinguishable from real cells. On the contrary, synthetic cells generated by ZINB-WaVE and SPARSim exhibit spurious patterns unseen in real cells.

To quantify the similarity between synthetic cells and real test cells, we use the median integration local inverse Simpson’s index (miLISI) [80], whose value is between 1 and 2, with a larger value indicating a greater similarity. Specifically, we compute an integration local inverse Simpson’s index (iLISI) to represent the effective number of cell labels (with 1 meaning synthetic or real cells only, and 2 meaning equal numbers of synthetic and real cells) in the local neighborhood of each (synthetic or real) cell; the closer iLISI is to 2, the more equal presence synthetic and real cells have in the local neighborhood. Taking the median of the iLISIs of all cells, we obtain the miLISI, which quantifies the overall mixing of synthetic cells with real cells. Using the R package LISI [80], we calculate the miLISI value for each of the overlaying 2D plots containing real and synthetic cells (Fig. 4 and Supplementary Figs. S29–S31), and we find that scDesign2 consistently leads to the highest miLISI value, with greater advantages in 2D tSNE plots than 2D PCA plots. Since 2D t-SNE projection preserves cell clusters better than 2D PCA does and is more widely used for visualizing single-cell gene expression data, our results suggest that the synthetic data by scDesign2 best capture the cluster structure in real cells. Together, the miLISI values confirm the superb performance and the realistic nature of scDesign2.

These three sets of analyses also verify the advantage of using copula in scDesign2. Compared with scDesign2, its w/o copula variant, as expected, cannot capture gene correlations at all (Fig. 3a, Fig. 2, Supplementary Figs. S6a–S16a, and Supplementary Figs. S1–S5). As a result, the synthetic data generated by the w/o copula variant do not resemble the corresponding real data in 2D visualization (Fig. 4 and Supplementary Figs. S29–S31).

In addition to its realistic nature, scDesign2 also has two more unique advantages over ZINB-WaVE and SPARSim. Unlike the other two simulators, scDesign2 only considers genes as features and models their joint distribution, and it regards cells as observations instead of features. This formulation is aligned with the statistical thinking that genes are fixed quantities but cells are randomly sampled from a population of cells. Thanks to this principled formulation, scDesign2 can generate synthetic cells of any number, in contrast to ZINB-WaVE and SPARSim that can only generate the same number of synthetic cells as real cells. It is also worth noting that, although scDesign2 does not explicitly model the distribution of cell library sizes, it recovers that distribution rather faithfully (see the cell library size distributions in Fig. 3a and Supplementary Figs. S6a– S16a). This is achieved by modeling joint gene distributions and accounting for gene correlations through the use of copula. Compared to scGAN, the training of scDesign2 is fast and does not rely on a large number of input real cells for good training quality.

### Refinement of scDesign2 training: calibration of cell types by ROGUE scores

For a dataset containing multiple cell types, scDesign2 needs to fit a model to each cell type before generating synthetic data. To ensure the quality of its synthetic data, scDesign2 must have one of its count models (ZINB model and its three simplified variants) fit well to each gene’s real expression levels in each cell type; otherwise, the synthetic data may not well mimic real data due to the poorness of model fitting. We observe this issue in the 10x Genomics dataset (Fig. 4a), where some cell types such as transit-amplifying early (TA.Early) cells and goblet cells are composed of discrete sub-clusters in 2D tSNE illustration. As a result, some genes’ expression levels within one of such cell types cannot be fit well by scDesign2’s count models, leading to a discrepancy between synthetic data and real data (synthetic TA.Early and goblet cells do not appear to have cell sub-clusters in 2D tSNE illustration).

To address this issue, we calibrate each cell type using the ROGUE score [81], which measures the homogeneity of that cell type, before training scDesign2. Concretely, we first partition the cell type into sub-clusters using the Louvain clustering algorithm [82] in the Seurat R package [41]. Employing varying resolution parameters in the Louvain algorithm, we partition the cell type into a varying number of sub-clusters. Second, we calculate the ROGUE score of every sub-cluster, and then we compute the average ROUGE score across sub-clusters for each number of sub-clusters, ranging from 1 to 6. Third, we examine how the average ROGUE score increases as the number of sub-clusters increases (Fig. 5a), together with 2D t-SNE visualization (Fig. 5b), to determine an appropriate number of sub-clusters, which is usually the “elbow point” where the average ROGUE score saturates.

**Figure 5:**
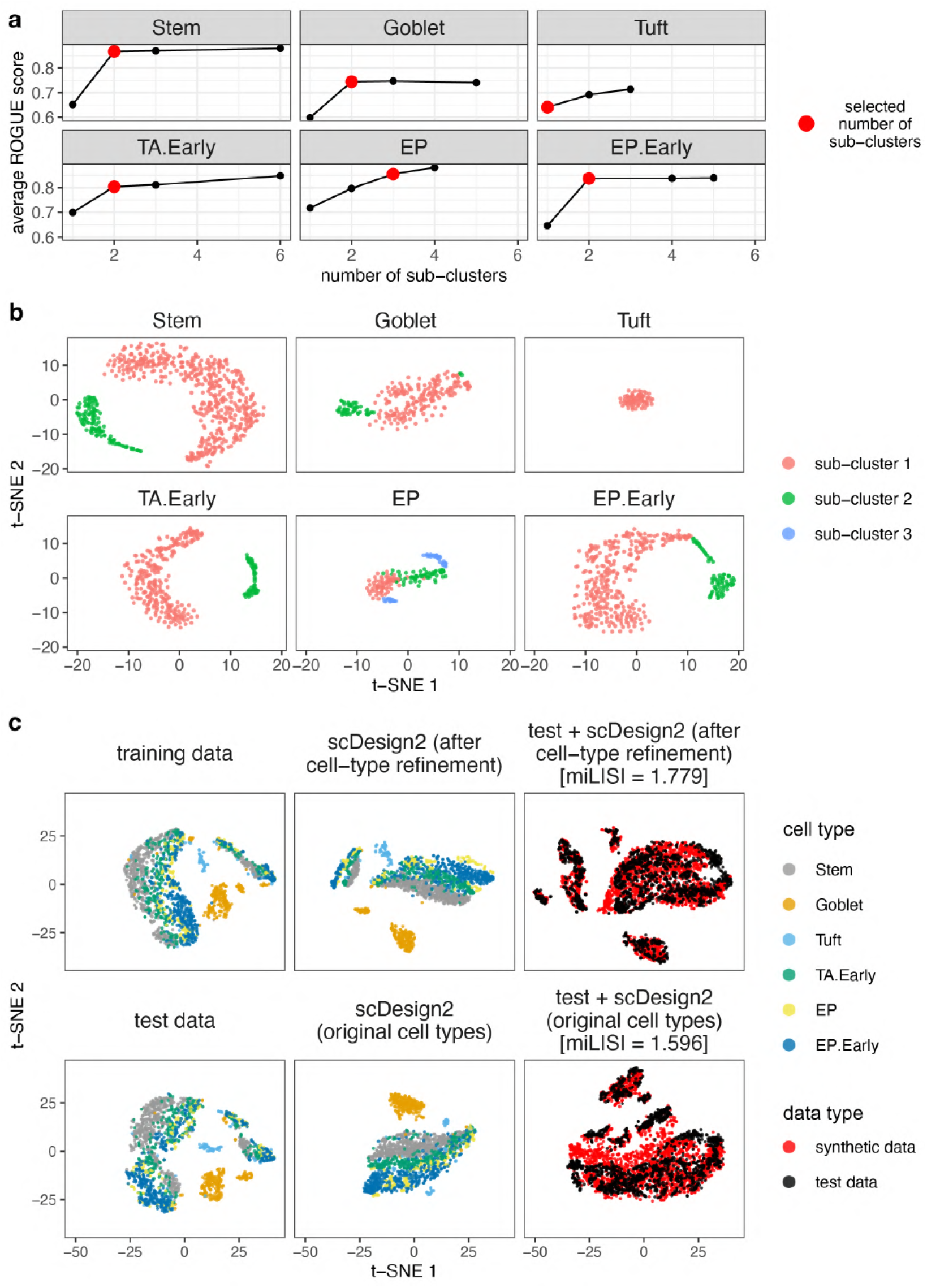
Application of ROGUE scores combined with dimensionality reduction plots to refine cell types before training scDesign2. This refinement approach is demonstrated on the 10x Genomics dataset. (a) In each cell type, the relationship between the average ROGUE score across sub-clusters and the number of sub-clusters. Before a ROGUE score is calculated for each sub-cluster, the Louvain clustering algorithm is applied to each cell type with a varying resolution parameter so that a varying number of sub-clusters is obtained. Based on how the average ROGUE score saturates, a number of sub-clusters is selected and marked in red for each cell type. (b) The t-SNE plots of each cell type with the sub-clusters, whose number is marked in (a), labelled with distinct colors. (c) The t-SNE plots of training data (top left), test data (bottom left), synthetic data of scDesign2 trained with the refined sub-clusters (middle left) or the original cell types (middle bottom), and combination of test data with each set of synthetic data. Gene expression counts are transformed as log(l + count) before dimensionality reduction. We find that, after the cell-type refinement, the simulated data of scDesign2 resemble the real data better, as indicated by the higher miLISI value.

Applying this strategy to refining the six cell types in the 10x Genomics dataset, we observe that, after being trained with the refined cell types, scDesign2 generates more realistic synthetic data (Fig. 5c; the miLISI value increases from 1.596 to 1.779).

### Application 1: scDesign2 generates realistic synthetic data for other single-cell expression count-based technologies

Beyond scRNA-seq data, we demonstrate that scDesign2 can also generate realistic synthetic data for other single-cell count-based technologies that do not necessarily use next-generation sequencing, as long as individual genes’ count distributions can be well approximated by Poisson, ZIP, NB or ZINB. For instance, single-cell spatial transcriptomics technologies, usually based on fluorescence in situ hybridization (FISH), are known to yield Poisson or NB distributed counts [83, 84]. The versatility of scDesign2 is endowed by its data-driven way of selecting marginal distributions for individual genes, regardless of each distribution being Poisson or NB, zero-inflated or not.

We demonstrate the accuracy of scDesign2 based on two single-cell spatial transcriptome datasets: one dataset of cells in the mouse hypothalamic preoptic region measured by multiplexed error robust fluorescence in situ hybridization (MERFISH) [85] and another dataset of cells in the mouse hippocampal area CA1 measured by probabilistic cell typing by in situ sequencing (pciSeq) [86], a newly developed spatial transcriptome profiling technology. Both datasets contain labeled cell types. Due to the lack of simulators specifically designed for single-cell spatial transcriptome data, we still benchmark scDesign2 against its w/o copula variant, as well as ZINB-WaVE and SPARSim, the two simulators that preserve genes. Note that for all the simulators considered, they only generate gene counts, not spatial coordinates, for synthetic cells. Similar to our previous analysis, for each cell type in each dataset, we randomly split the cells into two halves, with one half (“training data”) to be used for training every simulator and the other half (“test data”) to serve as the benchmark standard to be compared with the synthetic data generated by each simulator. Fig. 6 and Supplementary Fig. S32 demonstrate the 2D visualization of each real dataset, the corresponding synthetic data generated by each simulator, as well as the combination of test data and each synthetic dataset. For both technologies and in both t-SNE and PCA visualization, scDesign2 outperforms SPARSim and ZINB-WaVE by generating synthetic data that most resemble the real data. In particular, scDesign2 consistently achieves the highest miLISI values in the 2D visualization plots of combined data, indicating that the synthetic cells generated by scDesign2 are least distinguishable from real cells. These results confirm the versatility and robustness of scDesign2.

**Figure 6:**
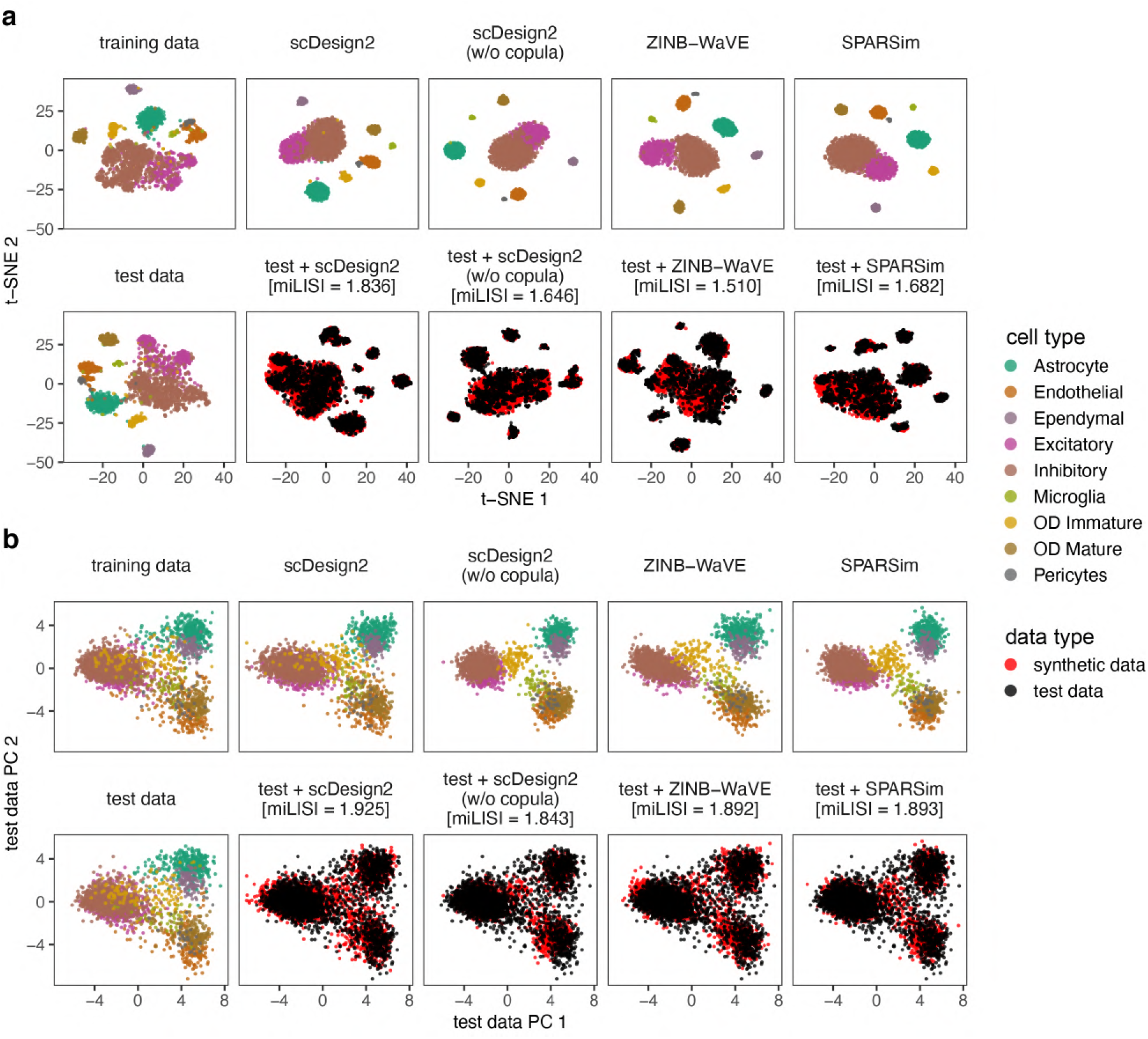
Comparison of MERFISH data and synthetic data generated by scDesign2, its variant without copula, ZINB-WaVE, and SPARSim in 2D visualization. (a) t-SNE plots and (b) principle component (PC) plots of training data, test data, synthetic data generated by each simulator, and combinations of test data and each synthetic dataset. Gene expression counts are transformed as log(l + count) before dimensionality reduction. miLISI is short for median integration local inverse Simpson’s Index, a higher value of which indicates that the simulated data mix better with the test data in the 2D visualization plot. By visually inspecting the patterns in these plots as well as comparing the miLISI values, we find that the synthetic data generated by scDesign2 most resemble the test data.

### Application 2: scDesign2 guides experimental design and computational method benchmarking in cell clustering

Cell clustering is a ubiquitous computational task in single-cell research. Here we demonstrate how scDesign2 can guide experimental design (i.e., deciding the optimal cell number and sequencing depth) and benchmark computational methods for the cell clustering task.

After training scDesign2 on each of the four scRNA-seq datasets generated by different experimental protocols (10x Genomics [76], CEL-Seq2 [77], and Fluidigm C1 [78], Smart-Seq2 [79]), we apply the trained scDesign2 to generate synthetic data under three experimental design scenarios: (1) varying sequencing depths, where the total number of reads (or UMIs) varies but the cell number is fixed; (2) varying cell numbers, where the number of sequenced cells varies but the sequencing depth is fixed; (3) fixing the per-cell average sequencing depth, where the both the number of sequenced cells and the total sequencing depth vary, but the average number of reads (or UMIs) in each cell is fixed. For each protocol, scDesign2 generates a synthetic dataset per sequencing depth and cell number.

To guide the choices of sequencing depth and cell number based on clustering accuracy, we apply two popular scRNA-seq cell clustering methods—Seurat (the kNN-Jaccard-Louvain algorithm) [41, 82] and SC3 [40]—to the synthetic datasets and use the adjusted mutual information (AMI) [87] and the adjusted Rand index (ARI) [88] as two clustering accuracy measures. Note that SC3 can be specified to output the same number of cell clusters as the annotated cell types, while Seurat cannot due to the nature of the Louvain algorithm it uses [82]. The results are summarized in Figs. 7–9 and Supplementary Figs. S33–S41.

**Figure 7:**
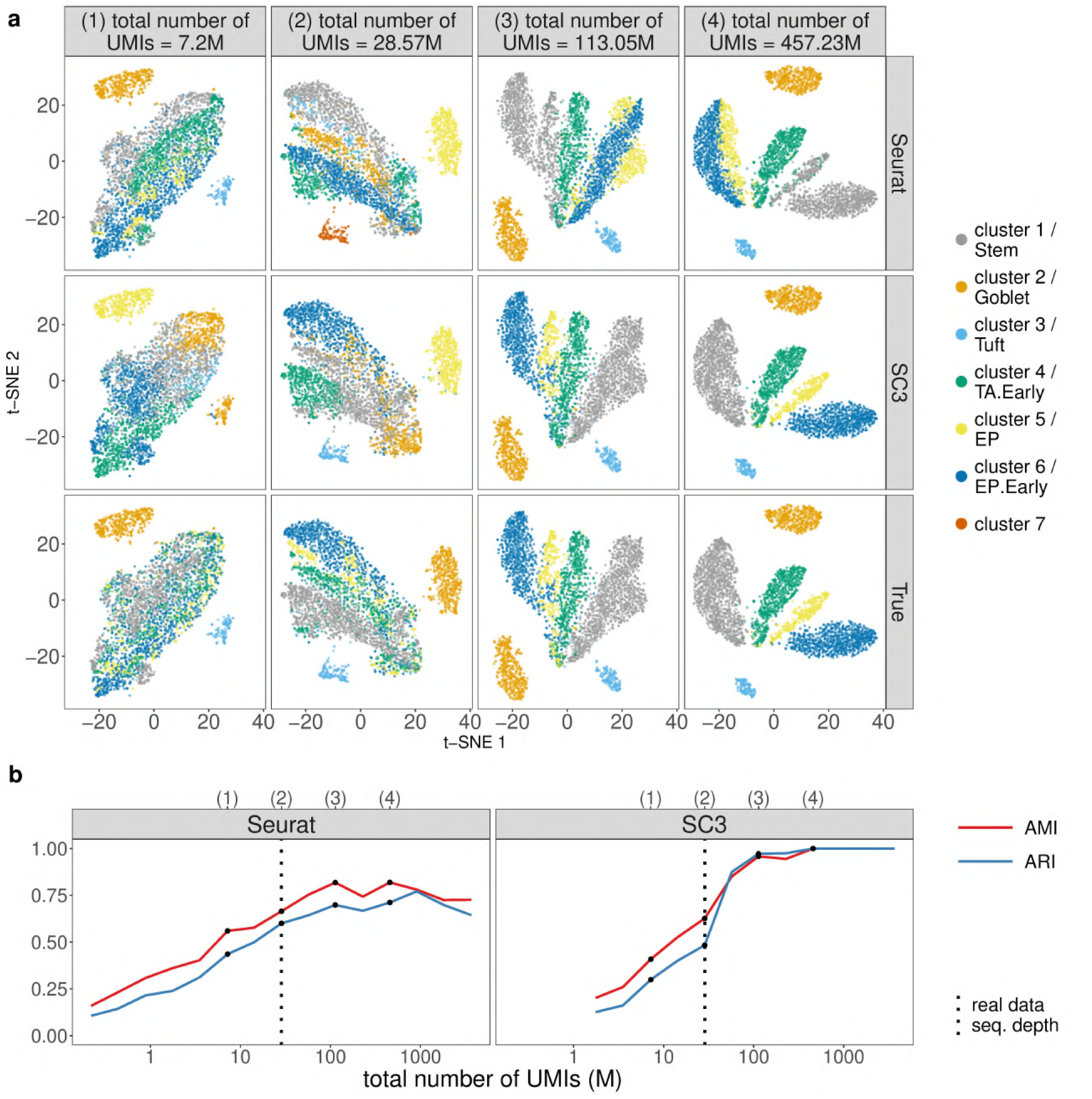
scDesign2 guides the choice of sequencing depth in cell clustering. scDesign2 generates synthetic 10x Genomics data with fifteen sequencing depths. Two cell clustering methods—Seurat and SC3—are applied to each synthetic dataset to partition cells into cell clusters. (a) t-SNE visualization of four synthetic datasets, where cells are labelled by Seurat clusters (top), SC3 clusters (middle), and annotated cell types (bottom). (b) Two clustering accuracy measures (AMI and ARI) vs. sequencing depth; left: Seurat; right: SC3. In (b), the results of the four sequencing depths in (a) are marked as dots and in the top, and the sequencing depth of the real dataset [76] is marked as vertical dashed lines.

For the first, varying-sequencing-depth scenario, we expect the clustering accuracy to improve as the sequencing depth increases, because a larger sequencing depth would better reveal every cell’s transcriptome profile and thus lead to better clustering. Moreover, we also expect there to be a saturation effect: the clustering accuracy no longer improves much after the sequencing depth increases to a point, which we regard as the optimal sequencing depth that balances clustering accuracy and budget. The results confirm our expectation. For the two UMI-based protocols 10x Genomics and CEL-Seq2, we observe the improvement and the saturation effect in clustering accuracy, based on both Seurat and SC3, as the sequencing depth increases. In detail, the saturation for 10x Genomics data occurs at 113.05 million UMIs for 3793 cells, while the real dataset has only 28.57 million UMIs (Fig. 7); the saturation for CEL-Seq2 data occurs at 42.72 million UMIs for 2279 cells, while the real dataset contains 172.14 million UMIs (Fig. S33). For the two non-UMI-based protocols Fluidigm C1 and Smart-Seq2, we observe the saturation effect even at the lowest sequencing depth we consider, likely due to the fact that these two protocols provide a deeper profiling of every cell than the UMI-based protocols do. In detail, the saturation for Fluidigm C1 data occurs at 26.74 (based on Seurat) or 110.52 (based on SC3) million reads for 317 cells, while the real dataset contains 869.24 million reads (Fig. S36); the saturation for Smart-Seq2 data occurs at 33.68 million reads for 1078 cells, based on both Seurat and SC3, while the real dataset contains 1074.97 million reads (Fig. S39). The t-SNE visualization supports the observed trends of clustering accuracy. In each t-SNE plot that corresponds to one sequencing depth and one set of cell clusters/types (by Seurat, SC3, or annotated cell types), synthetic cells are labelled by their cell clusters/types; contrasting a tSNE plot of cell clusters with that showing cell types illustrates clustering accuracy (Fig. 7a and Supplementary Figs. S33a, S36a, and S39a). In conclusion, for clustering purpose, we would recommend increasing the 10x Genomics sequencing depth to 113.05 million UMIs, if budget allows, and using SC3 for clustering; for CEL-Seq2, Fluidigm C1, and Smart-Seq2, we would recommend decreasing the sequencing depths to 42.72 million UMIs, 110.52 million reads, and 33.68 million reads, respectively, to save budget and using either Seurat or SC3 for clustering.

For the second, varying-cell-number scenario, we expect the clustering accuracy to first increase and then decrease as the cell number increases. The reason is that good clustering requires both a reasonable number of cells of each type and a clear-enough gene expression profile (where enough genes are captured) of every cell, thus posing a tradeoff—given the sequencing depth, the larger the cell number, the less clear each cell’s profile would be. Hence, as the cell number increases from low, while every cell’s profile is still clear, clustering accuracy increases; however, as the cell number reaches a point where every cell type has more than enough cells, further increasing the cell number would obscure every cell’s profile and deteriorate clustering accuracy. For the two UMI-based protocols 10x Genomics and CEL-Seq2, our expectation is confirmed: we observe an overall trend of clustering accuracy that first increases and then decreases (Fig. 8b and Supplementary Fig. S34b). In detail, for 10x Genomics data, both Seurat and SC3 have their accuracy maximized at 948 cells. This optimality is supported by the t-SNE visualization, which shows that the Seurat and SC3 cell clusters best agree with the annotated cell types at this optimal cell number (Fig. 8a). Hence, the real data cell number 3,793 is not optimal for distinguishing the annotated cell types by Seurat or SC3. For CEL-Seq2 data, Seurat and SC3 have optimal accuracy at 2,279 and 570 cells, respectively, also supported by the t-SNE visualization (Supplementary Fig. S34a). This suggests that the real data cell number 2,279 can lead to optimal cell clustering by Seurat. In contrast, for the two non-UMI-based protocols Fluidigm C1 and Smart-Seq2, we only observe a first-increasing-and-then-saturated trend of clustering accuracy as the cell number increases, without seeing the trend decreasing (except for SC3 on Smart-Seq2 data) (Supplementary Figs. S37b and S40b). A likely reason is that these two protocols can provide a clear profile of every cell up to a large cell number around 10,000 given their deep sequencing depths in real data (869.24 million reads in the Fluidigm C1 data and 1074.95 million reads in the Smart-Seq2 data). For both Seurat and SC3, the cell numbers at which their performance saturates are close to the cell numbers in real data: 317 cells in the Fluidigm C1 data and 1078 cells in the Smart-Seq2 data. In conclusion, we use scDesign2 to find that the cell numbers are close to being optimal in the CEL-Seq2, Fluidigm C1, and Smart-Seq2 datasets. For 10x Genomics, we would recommend decreasing the cell number to 948 cells (while keeping the sequencing depth at 28.58 million UMIs) to optimize the clustering accuracy by either Seurat or SC3.

**Figure 8:**
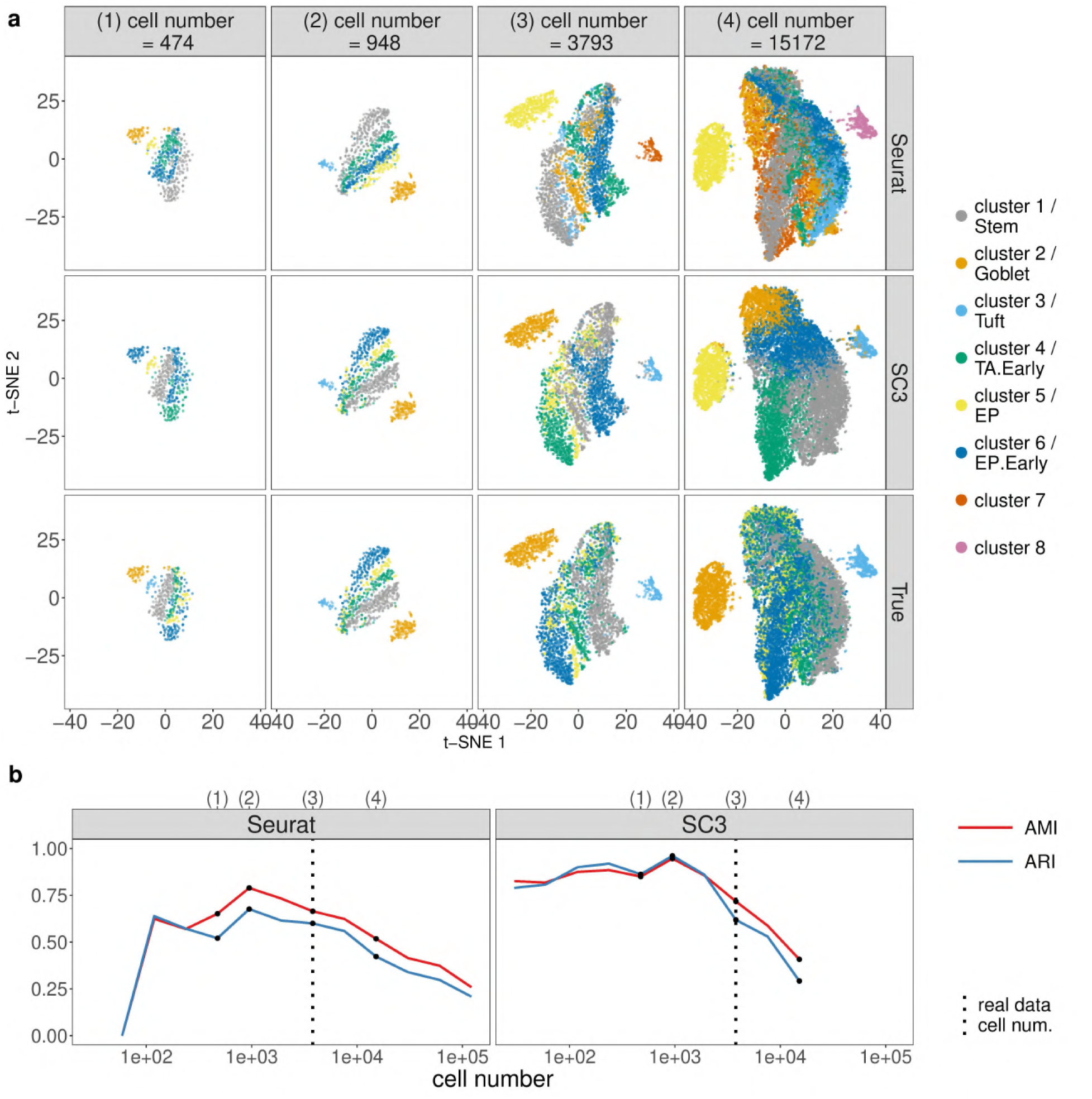
scDesign2 guides the choice of cell number in cell clustering, in the case where the total sequencing depth is kept as fixed. scDesign2 generates synthetic 10x Genomics data with twelve cell numbers. Two cell clustering methods—Seurat and SC3—are applied to each synthetic dataset to partition cells into cell clusters. (a) t-SNE visualization of four synthetic datasets, where cells are labelled by Seurat clusters (top), SC3 clusters (middle), and annotated cell types (bottom). (b) Two clustering accuracy measures (AMI and ARI) vs. sequencing depth; left: Seurat; right: SC3. In (b), the results of the four cell numbers in (a) are marked as dots and in the top, and the cell number of the real dataset [76] is marked as vertical dashed lines.

For the third, fixing-average-sequencing-depth scenario, we expect the clustering accuracy to improve as the cell number (and also the total sequencing depth) increases, because more cells will make the identification of cell types easier. Moreover, we expect there to be a saturation effect: the clustering accuracy no longer improves much after the cell number increases to a point, which we regard as the optimal cell number that balances clustering accuracy and budget. The results confirm our expectation. In all four protocols, we observe the expected trend of clustering accuracy for both Seurat and SC3, as well as the saturation effect, which is more obvious for Seurat. In detail, the saturation for 10x Genomics data occurs at 948 cells (based on Seurat), or 3793 cells (based on SC3), while the real dataset has 3,793 cells (Fig. 9); the saturation for CEL-Seq2 data occurs at 1,140 cells, while the real dataset has 2,279 cells (Fig. S35); the saturation for Fluidigm C1 data occurs at 317 cells (based on Seurat), which is the same cell number as the real dataset, while the optimal clustering accuracy occurs at 1,268 cells based on SC3 (Fig. S38); the saturation for Smart-Seq2 data occurs at 4,312 cells, while the real dataset has 1,078 cells (Fig. S41). In conclusion, when the average read (or UMI) count per cell is kept as fixed, for clustering purpose, we recommend keeping the cell number as in the original design for 10x Genomics and using SC3 for clustering; for CEL-Seq2, we recommend decreasing the cell number to 1,140 to save budget and using Seurat for clustering; for Fluidigm C1, if budget allows, we recommend increasing the cell number to 1,268 and using SC3 for clustering; for Smart-Seq2, if budget allows, we recommend increasing the cell number to 4,312 and using either Seurat or SC3 for clustering.

**Figure 9:**
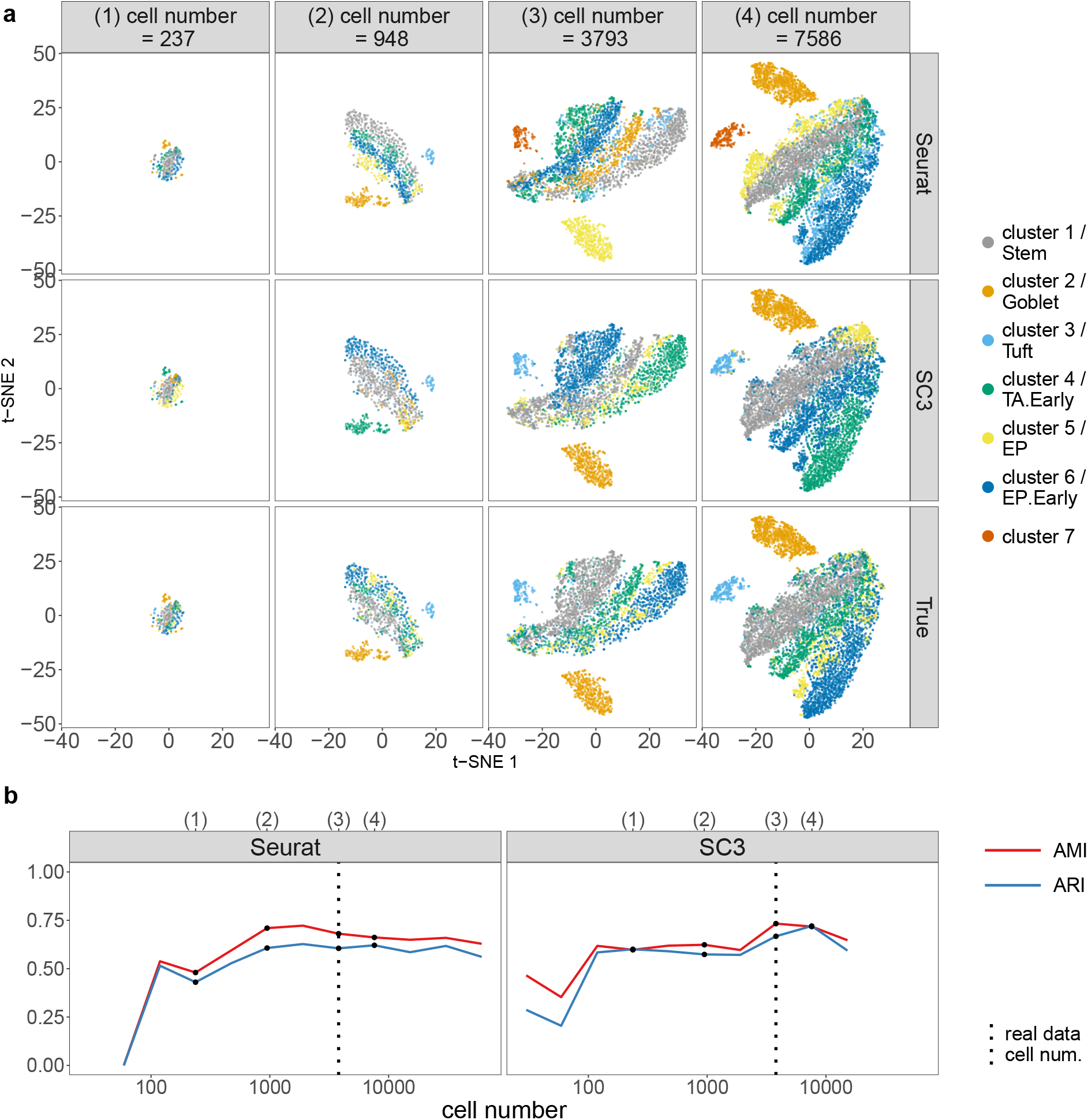
scDesign2 guides the choice of cell number in cell clustering, in the case where the average sequencing depth is kept as fixed. scDesign2 generates synthetic 10x Genomics data with eleven cell numbers. Two cell clustering methods—Seurat and SC3—are applied to each synthetic dataset to partition cells into cell clusters. (a) t-SNE visualization of four synthetic datasets, where cells are labelled by Seurat clusters (top), SC3 clusters (middle), and annotated cell types (bottom). (b) Two clustering accuracy measures (AMI and ARI) vs. sequencing depth; left: Seurat; right: SC3. In (b), the results of the four cell numbers in (a) are marked as dots and in the top, and the cell number of the real dataset [76] is marked as vertical dashed lines.

Beyond experimental design, scDesign2 also provides a comprehensive comparison of Seurat and SC3 across sequencing depths and cell numbers. Overall, both methods demonstrate superb accuracy in a wide range of sequencing depths and cell numbers for every protocol. At close-to-optimal sequencing depths and cell numbers for each method, SC3 has better accuracy than Seurat. However, Seurat and SC3 has different robustness: Seurat is a more robust method for 10x genomics data when the sequencing depth is too low or the cell number is too large (Figs. 7b–9b), while SC3 is more robust when the cell number is small for CEL-Seq2 (Supplementary Figs. S34b–S35b), Fluidigm C1 (Supplementary Figs. S37b–S38b), and Smart-Seq2 (Supplementary Figs. S40b–S41b). This finding is consistent with the fact that SC3 is an ensemble method that is more robust against a small number of cells but cannot be easily scaled up when the cell number is too large.

### Application 3: scDesign2 guides experimental design and computational method benchmarking in rare cell type detection

Rare cell type detection is another important application of scRNA-seq, whose high-throughput profiling of cells opens an unprecedented opportunity to identify unknown cell types that are often rare but critical. Here we demonstrate how scDesign2 can guide experimental design (i.e., deciding the optimal cell number and sequencing depth) and benchmark computational methods for the rare cell type detection task.

From the 10x Genomics dataset of mouse intestine epithelial tissue [76], we select six cell types—stem cells (Stem), goblet cells (Goblet), tuft cells (Tuft), early transit amplifying cells (TA.Early), enterocyte progenitors (EP), and early enterocyte progenitors (EP.Early), among which Tuft is the rare cell type [89] and has a proportion less than 5% among the six cell types. After training scDesign2 on this dataset, we use scDesign2 to generate synthetic data under three experimental design scenarios: (1) varying sequencing depths, where the total number of UMIs varies but the cell number is fixed; (2) varying cell numbers, where the number of sequenced cells varies but the sequencing depth is fixed; (3) fixing the per-cell average sequencing depth, where the both the number of sequenced cells and the total sequencing depth vary, but the average number of reads (or UMIs) in each cell is fixed. For every sequencing depth and cell number, scDesign2 generates a synthetic dataset.

To guide the choices of sequencing depth and cell number based on rare-cell-type detection accuracy, we apply two popular methods—FiRE [46] and GiniClust2 [45]—to the synthetic datasets and evaluate four accuracy measures: precision (the percentage of truly rare cells among the detected rare cells), recall (the percentage of detected rare cells among the truly rare cells), F1-score (the harmonic mean of the precision and recall), and AUPRC (the area under the precision-recall curve). Since GiniClust2 does not allow adjustment of its detection threshold, we cannot calculate its AUPRC. However, as most users of FiRE would stick with its default threshold, the AUPRC measure is not as informative as the other three measures from a user’s perspective.

For the first, varying-sequencing-depth scenario, we expect that the detection accuracy would improve as the sequencing depth increases and there would be a saturation effect, similar to our expectation for cell clustering. The detection accuracy of FiRE and GiniClust2 roughly confirm our expectation. Across twelve sequencing depths ranging from 1.76 to 3612.4 million UMIs (with the cell number fixed as 3793, the number of cells in real data), we observe an overall trend of increasing detection accuracy with few exceptions (Fig. 10). For FiRE, its accuracy exhibits saturation after the sequencing depth reaches 457.23 million UMIs (Fig. 10a & c), while for GiniClust2 the saturation occurs earlier at a sequencing depth of 113.05 million UMIs (Fig. 10b & d). The t-SNE visualization supports the observed trends of precision and recall. In each t-SNE plot that corresponds to one sequencing depth and one detection method (FiRE or GiniClust2), synthetic cells are labelled as one of four types: true positive (TP; the rare cells correctly detected), false positive (FP; the unrare cells falsely detected), false negative (FN; the rare cells falsely undetected), and true negative (TN; the unrare cells correctly undetected). The numbers of TP, FP, FN, and TN cells determine the precision and recall: a large precision requires many TP cells and few FP cells; a large recall requires many TP cells and few FN cells. Notably, the abnormal accuracy of GiniClust2 at 457.23 million UMIs (Fig. 10d) is explained by the t-SNE visualization (Fig. 10b), which shows that GiniClust2 misidentifies the second largest cell cluster as the rare cell type, leads to many FP and FN cells, and results in close to zero precision and recall. Combining the FiRE and GiniClust2 results, we conclude that the real data sequencing depth at 28.57 million UMIs for 3793 cells is not optimal for detecting the rare cell type Tuft (Fig. 10c–d). If budget allows, we would recommend increasing the sequencing depth to 113.06 million UMIs and use GiniClust2 to detect tuft cells.

**Figure 10:**
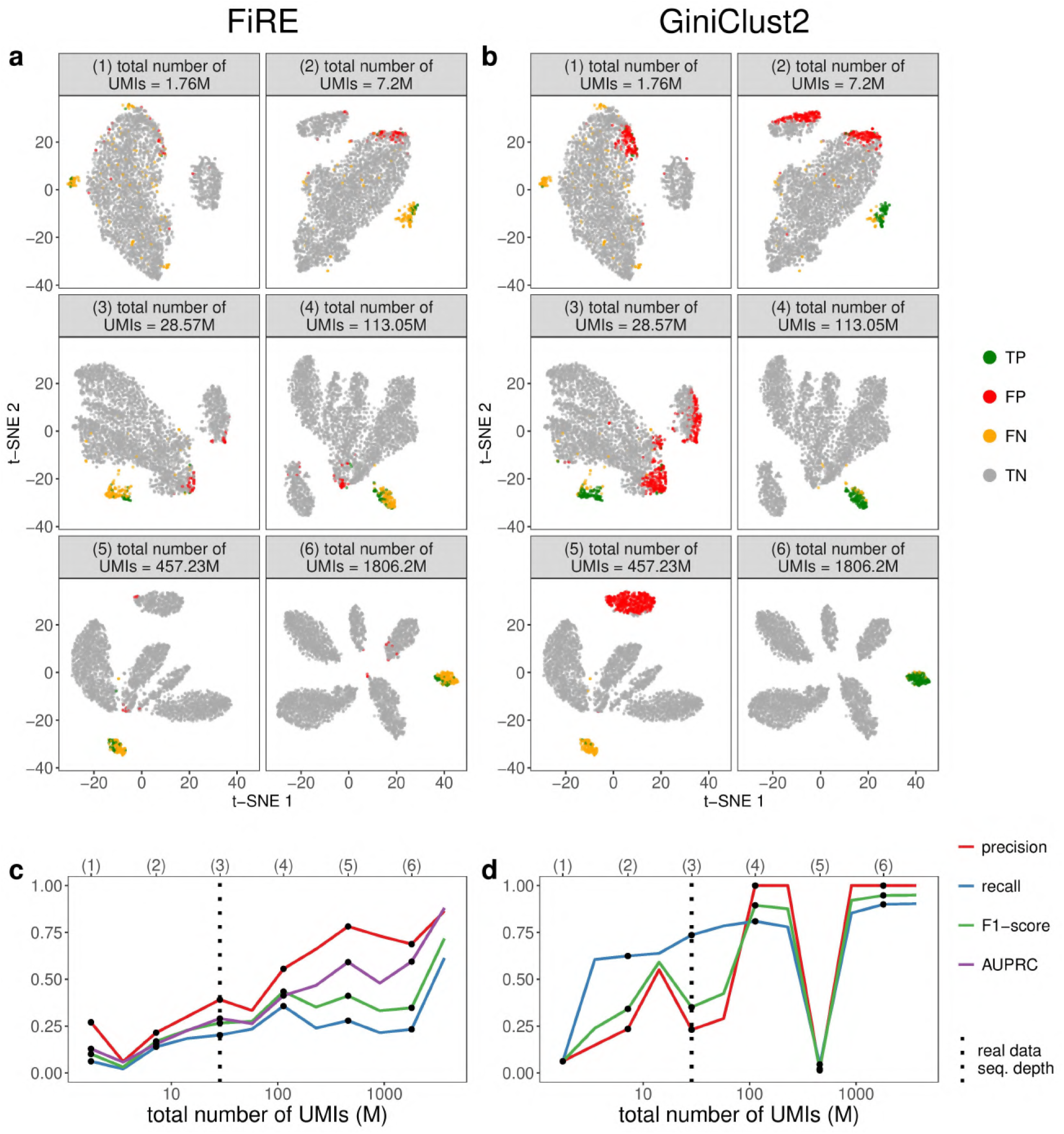
scDesign2 guides the choice of sequencing depth in rare cell type detection. scDesign2 generates synthetic 10x Genomics data with twelve sequencing depths. Two rare-cell-type detection methods—FiRE and GiniClust2—are applied to each synthetic dataset to detect rare cell types. (a) t-SNE visualization of six synthetic datasets and identification results—true positive (TP), false positive (FP), false negative (FN), and true negative (TN) cells—of FiRE in each dataset. (b) t-SNE visualization of the same six synthetic datasets and identification results of GiniClust2 in each dataset. (c) Four identification accuracy measures by FiRE (precision, recall, F1-score, and AUPRC) vs. sequencing depth. (d) Three identification accuracy measures by GiniClust2 (precision, recall, and F1-score) vs. sequencing depth. In (c) and (d), the results of the six sequencing depths in (a) and (b) are marked as dots and in the top, and the sequencing depth of the real dataset [76] is marked as vertical dashed lines.

For the second, varying-cell-number scenario, we expect the detection accuracy to first increase and then decrease as the cell number increases, similar to our expectation for cell clustering. Again, the detection accuracy of FiRE and GiniClust2 confirm our expectation. Across thirteen cell numbers ranging from 29 to 121,376 (with the sequencing depth fixed as 28.57 million UMIs, the same as in real data), we observe an overall trend of detection accuracy that first increases and then decreases (Fig. 11). For both FiRE and GiniClust2, their F1-scores are optimal at 1,896 cells (Fig. 11c–d). This optimality is supported by the t-SNE visualization, which shows a plot of synthetic cells with TP, FP, FN, and TN labels for every cell number and each detection method (Fig. 11a–b). Hence, the real data cell number 3793 is not optimal for detecting tuft cells given the total sequencing depth of 28.57 million UMIs. If the detection of tuft cells is a primary goal and the sequencing depth cannot be increased due to budget constraints, we would recommend decreasing the cell number of 1896 cells and use GiniClust2 to detect tuft cells.

**Figure 11:**
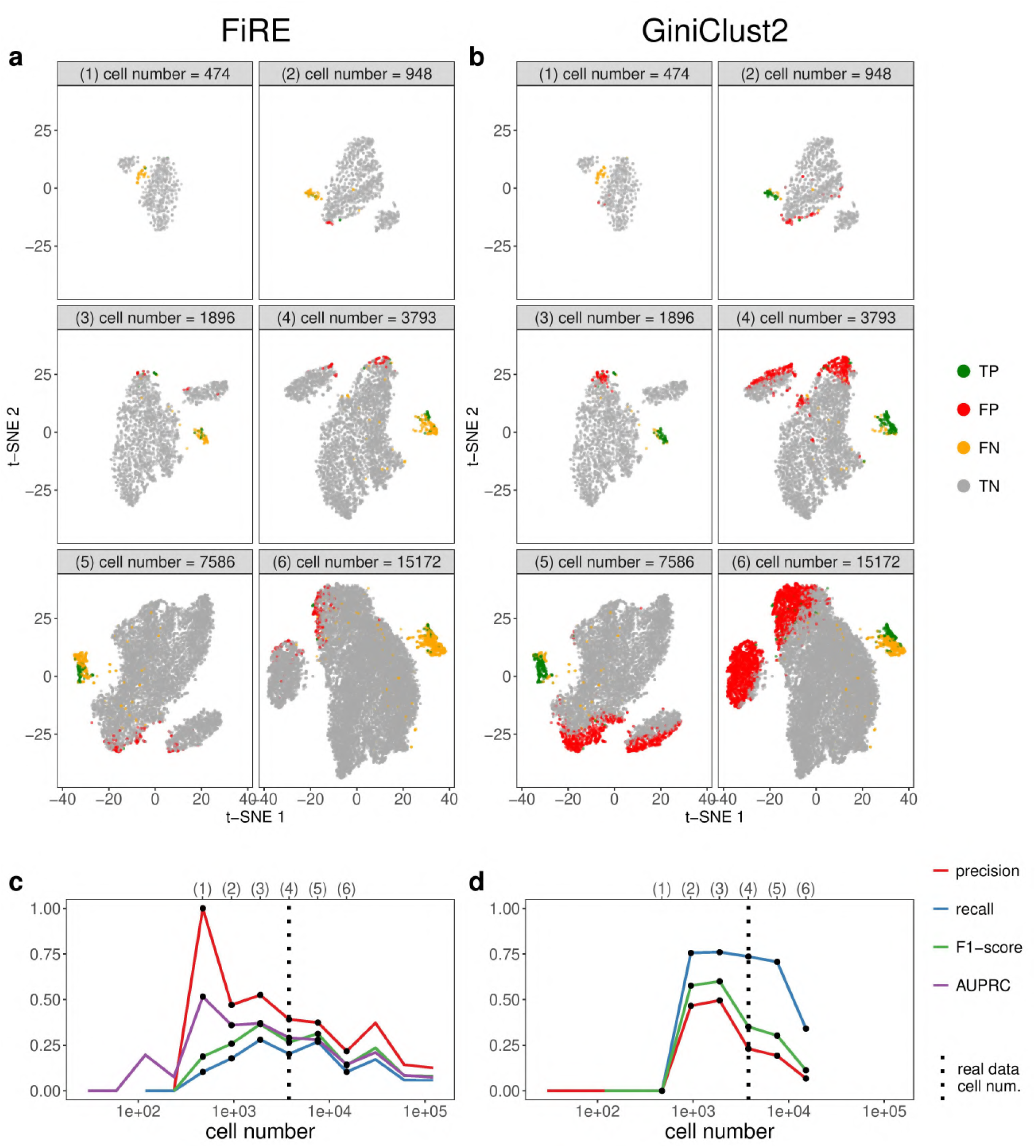
scDesign2 guides the choice of cell number in rare cell type detection, in the case where the total sequencing depth is kept as fixed. scDesign2 generates synthetic 10x Genomics data with thirteen cell numbers. Two rare-cell-type detection methods—FiRE and GiniClust2—are applied to each synthetic dataset to detect rare cell types. (a) t-SNE visualization of six synthetic datasets and identification results—true positive (TP), false positive (FP), false negative (FN), and true negative (TN) cells—of FiRE in each dataset. (b) t-SNE visualization of the same six synthetic datasets and identification results of GiniClust2 in each dataset. (c) Four identification accuracy measures by FiRE (precision, recall, F1-score, and AUPRC) vs. cell number. (d) Three identification accuracy measures by GiniClust2 (precision, recall, and F1-score) vs. cell number. In (c) and (d), the results of the six cell numbers in (a) and (b) are marked as dots and in the top, and the cell number of the real dataset [76] is marked as vertical dashed lines. Whenever there is no line for a cell number, FiRE or GiniClust2 does not detect any rare cells or fails.

For the third, fixing-average-sequencing-depth scenario, we expect the detection accuracy to first increase and then saturate as the cell number increases, similar to our expectation for cell clustering. The detection accuracy of FiRE roughly confirms our expectation, while the detection accuracy of GiniClust2 deviates from this trend (Fig. 12). For FiRE, across thirteen cell numbers ranging from 29 to 121,376 (with the average sequencing depth fixed as 7.53k UMIs per cell, the same as in real data), the F1-score reaches an early local maximum at 474 cells, and then stays relatively stable. A similar trend can be seen for the other three accuracy measures: precision, recall, and AUPRC. For GiniClust2, across nine cell numbers ranging from 29 to 7,586, the F1-score reaches a global maximum at 237 cells, and then it decreases as the cell number further increases. This is mainly due to the increasing proportion of FPs in the discovered rare cells, as indicated by the plunging precision curve. The recall, on the other hand, stays relatively stable after the optimal cell number. The t-SNE visualization supports the observed trends of these accuracy measures. For example, we can see that for GiniClust2, when the cell number reaches 1000, more cells are labelled as FP, as shown in subpanels (4)-(6) of Fig. 12b. In summary, if the goal is to detect tuft cells and the average sequencing depth is fixed as 7.53k UMIs per cell, we recommend using GiniClust2 and decreasing the number of cells to 237.

**Figure 12:**
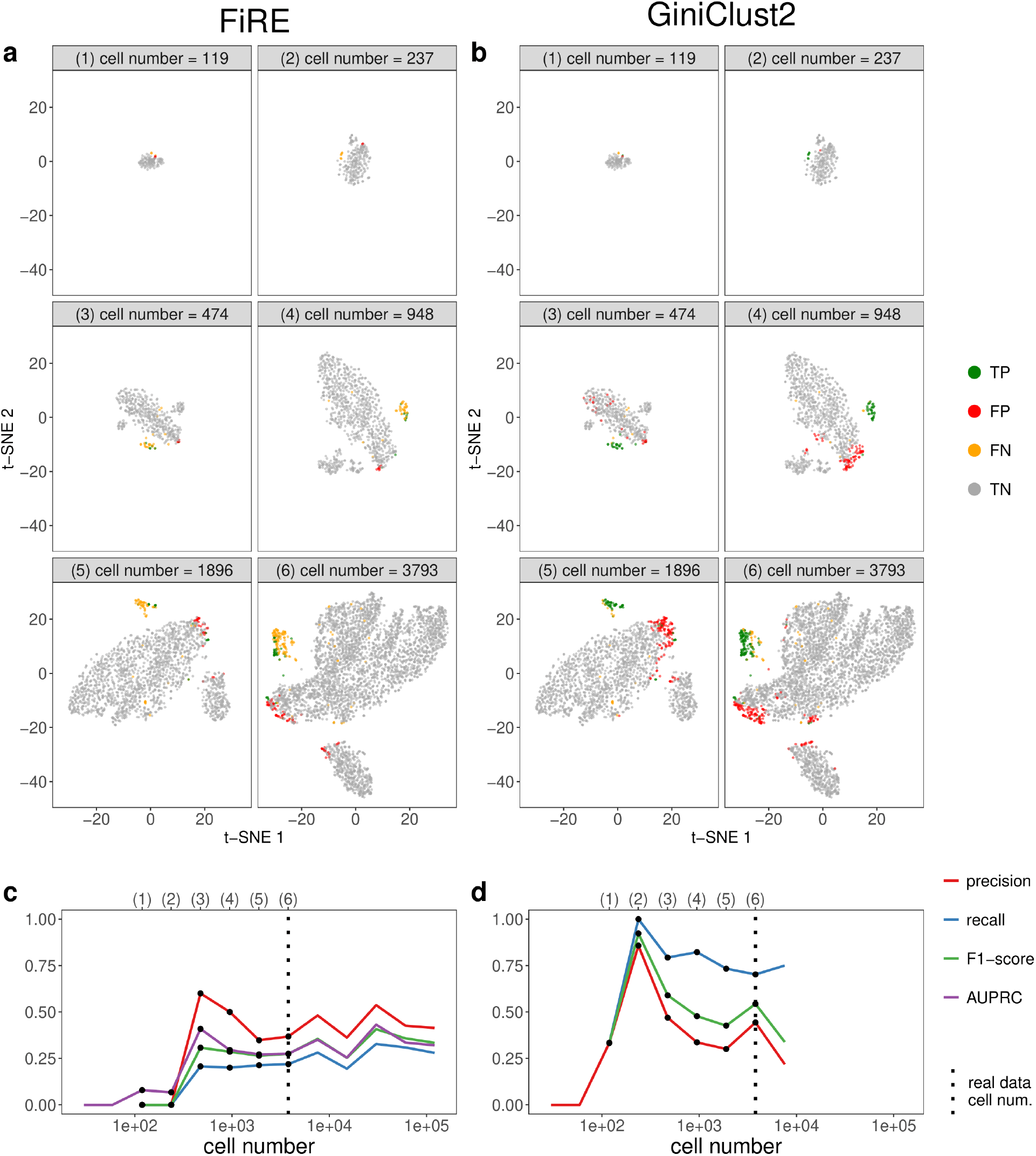
scDesign2 guides the choice of cell number in rare cell type detection, in the case where the average sequencing depth is kept as fixed. scDesign2 generates synthetic 10x Genomics data with thirteen cell numbers. Two rare-cell-type detection methods—FiRE and GiniClust2—are applied to each synthetic dataset to detect rare cell types. (a) t-SNE visualization of six synthetic datasets and identification results—true positive (TP), false positive (FP), false negative (FN), and true negative (TN) cells—of FiRE in each dataset. (b) t-SNE visualization of the same six synthetic datasets and identification results of GiniClust2 in each dataset. (c) Four identification accuracy measures by FiRE (precision, recall, F1-score, and AUPRC) vs. cell number. (d) Three identification accuracy measures by GiniClust2 (precision, recall, and F1-score) vs. cell number. In (c) and (d), the results of the six cell numbers in (a) and (b) are marked as dots and in the top, and the cell number of the real dataset [76] is marked as vertical dashed lines. Whenever there is no line for a cell number, FiRE or GiniClust2 does not detect any rare cells or fails.

In addition to assisting experimental design, scDesign2 also provides an objective comparison of FiRE and GiniClust2 across sequencing depths and cell numbers. Figs. 10–12 show that GiniClust2 has much better accuracy than FiRE at close-to-optimal sequencing depths and cell numbers. However, FiRE is a more robust method that it can successfully run at all sequencing depths and cell numbers, while GiniClust2 fails when the cell number is too small or too large (GiniClust3 may have addressed this large-cell-number issue [90]). This finding is consistent with the methodological difference between the two methods: FiRE detects rare cells via an outlier detection approach, while GiniClust2 first performs cell clustering and then identifies the cells in small clusters as rare cells. The requirement of cell clustering explains why GiniClust2 fails when the cells exhibit no clear clusters and why it works well when rare cells form small clear clusters. In contrast, outlier detection has no requirement on cluster patterns, and this explains why FiRE is robust.

## Discussion

In this article, we propose scDesign2, a transparent simulator for single-cell gene expression count data. Our development of scDesign2 is motivated by the pressing challenge to generate realistic synthetic data for various scRNA-seq protocols and other single-cell gene expression count-based technologies. Unlike existing simulators including our previous simulator scDesign, scDesign2 achieves six properties: protocol adaptiveness, gene preservation, gene correlation capture, flexible cell number and sequencing depth choices, transparency, and computational and sample efficiency. This achievement of scDesign2 is enabled by its unique use of the copula statistical framework, which combines marginal distributions of individual genes and the global correlation structure among genes. As a result, scDesign2 has the following methodological advantages that contribute to its high degree of transparency. First, it selects a marginal distribution from four options (Poisson, ZIP, NB, and ZINB) for each gene in a data-driven manner to best capture and summarize the expression characteristics of that gene. Second, it uses a Gaussian copula to estimate gene correlations, which will be used to generate synthetic single-cell gene expression counts that preserve the correlation structures. Third, it can generate gene expression counts according to user-specified sequencing depth and cell number.

We have performed a comprehensive set of benchmarking and real data studies to evaluate scDesign2 in terms of its accuracy in generating synthetic data and its efficacy in guiding experimental design and benchmarking computational methods. Based on four scRNA-seq protocols and 12 cell types, our benchmarking results demonstrate that scDesign2 better captures gene expression characteristics in real data than eight existing scRNA-seq simulators do. In particular, among the four simulators that aim to preserve gene correlations, scDesign2 achieves the best accuracy. Moreover, we demonstrate the capacity of scDesign2 in generating synthetic data of other single-cell count-based technologies including MERFISH or pciSeq, two single-cell spatial transcriptomics technologies. After validating the realistic nature of synthetic data generated by scDesign2, we use real data applications to demonstrate how scDesign2 can guide the selection of cell number and sequencing depth in experimental design, as well as how scDesign2 can benchmark computational methods for cell clustering and rare cell type identification.

Since scRNA-seq data typically contain tens of thousands of genes, the estimation of the copula gene correlation matrix is a high dimensional problem. This problem can be partially avoided by only estimating the copula correlation matrix of thousands of moderately to highly expressed genes. We use a simulation study to demonstrate why this approach is reasonable (Supplementary Figs. S42 and S43), and a more detailed discussion is in the Methods section. To summarize, the simulation results suggest that, to reach an average estimation accuracy of ±0.3 of true correlation values among the top 1000 highly expressed genes, at least 20 cells are needed. To reach an accuracy level of ±0.2 for the top 1500 highly expressed genes, at least 50 cells are needed. With 100 cells, an accuracy level of ±0.1 can be reached for the top 200 highly expressed genes, and a slightly worse accuracy level can be reached for the top 2000 genes.

In the implementation of the `scDesign2 R` package, we control the number of genes for which copula correlations need to be estimated by filtering out the genes whose zero proportions exceed a user-specified cutoff. For all the results in this paper, the cutoff is set as 0.8. In Supplementary Table S1, we summarize the number of cells (*n*), i.e., the sample size, and the number of genes included for copula correlation estimation (*p*) in each of the 12 datasets used for benchmarking simulators. Based on Supplementary Figs. S42 and S43, we see that *p* appears to be too large for the CEL-Seq2, Fluidigm C1, and Smart-Seq2 datasets. This suggests that the results in this paper may be further improved by setting a more stringent cutoff for gene selection.

For future methodological improvement, there are other ways to address this high-dimensional estimation problem. For example, we can consider implementing sparse estimation (e.g., [91]) for the copula correlation matrix. Moreover, we can build a hierarchical model to borrow information across cell types/clusters. This will be useful for improving the model fitting for small cell types/clusters that may share similar gene correlation structures.

The current implementation of scDesign2 is restricted to single-cell datasets composed of discrete cell types, because the generative model of scDesign2 assumes that cells of the same type follow the same distribution of gene expression. However, many single-cell datasets exhibit continuous cell trajectories instead of discrete cell types. A nice property of the probabilistic model used in scDesign2 is that it is generalizable to account for continuous cell trajectories. First, we can use the Generalized Additive Model (GAM) [92–94] to model each gene’s marginal distribution of expression as a function of cell pseudotime, which can be computationally inferred from real data [50, 51, 53]. Second, the copula framework can be used to incorporate gene correlation structures along the cell pseudotime. Combining these two steps into a generative model, this extension of scDesign2 has the potential to overcome the current challenge in preserving gene correlations encountered by existing simulators for single-cell trajectory data, such as Splatter Path [65], dyngen [95], and PROSSTT [64]. Another note is that scDesign2 does not generate synthetic cells based on outlier cells that do not cluster well with any cells in well-formed clusters. This is not necessarily a disadvantage, neither is it a unique feature to scDesign2. In fact, all model-based simulators that learn a generative model from real data must ignore certain outlier cells that do not fit well to their model. Some outlier cells could either represent an extremely rare cell type or are just “doublets” [96–99], artifacts resulted from single-cell sequencing experiments. Hence, our stance is that ignorance of outlier cells is a sacrifice that every simulator has to make; the open question is the degree to which outlier cells should be ignored, and proper answers to this question must resort to statistical model selection principles.

Regarding the use of scDesign2 to guide the design of scRNA-seq experiments, although scDesign2 can model and simulate data from different scRNA-seq protocols and other singlecell expression count-based technologies, the current scDesign2 implementation is not yet applicable to cross-protocol data generation (i.e., training scDesign2 on real data of one protocol and generating synthetic data for another protocol) because of complicated differences in data characteristics among protocols. To demonstrate this issue, we use a multi-protocol dataset of peripheral blood mononuclear cells (PBMCs) generated for benchmarking purposes [19]. We select data of five cell types measured by three protocols, 10x Genomics, Drop-Seq, and Smart-Seq2, and we train scDesign2 on the 10x Genomics data. Then we adjust the fitted scDesign2 model for the Drop-Seq and Smart-Seq2 protocols by rescaling the mean parameters in the fitted model to account for the total sequencing depth and cell number, which are protocol-specific (see Methods for details). After the adjustment, we use the model for each protocol to generate synthetic data. Supplementary Fig. S44 illustrates the comparison of real data and synthetic data for each protocol. From the comparison, we observe that the synthetic cells do not mix well with the real cells for the two cross-protocol scenarios; only for 10x Genomics, the same-protocol scenario, do the synthetic cells mix well with the real cells.

To further illustrate the different data characteristics of different protocols, we compare individual genes’ mean expression levels in the aforementioned three protocols. We refer to Drop-Seq and Smart-Seq2 as the target protocols, and 10x Genomics as the reference protocol. First, we randomly partition the two target-protocol datasets and the reference-protocol dataset into two halves each; we repeat the partitions for 100 times and collect 100 sets of partial datasets, with each set containing two target-protocol partial datasets (one Drop-Seq and one Smart-Seq2) and two reference-protocol partial datasets (split from the 10x-Genomics dataset)—one of the latter is randomly picked and referred to as the “reference data.” Second, For every gene in each cell type, we take each set of partial datasets and compute two cross-protocol ratios, defined as the gene’s mean expression levels in the target-protocol partial datasets divided by its mean expression level in the reference data, and a within-protocol ratio, defined as the ratio of the gene’s mean expression level in the other reference-protocol partial dataset divided by that in the reference data; together, with the 100 sets of partial dataset, every gene in each cell type has 100 ratios for each of the two cross-protocol comparisons and 100 ratios for the within-protocol comparison. We apply this procedure to the top 50 and 2000 highly expressed genes in five cell types. Supplementary Figs. S45 and S46 show that, with the within-protocol ratios as a baseline control for each cell type and each target protocol, the cross-protocol ratios exhibit a strongly gene-specific pattern; moreover, there is no monotone relationship between the cross-protocol ratios and the mean expression levels of genes. This result confirms that there does not exist a single scaling factor to convert all genes’ expression levels from one protocol to another. However, an interesting phenomenon is that, for each target protocol, the cross-protocol ratios have similar patterns across cell types. This phenomenon sheds light on a future research direction of cross-protocol simulation for the cell types that exist in only one protocol, if the two protocols have shared cell types. In this scenario, we may train a model for each cell type in each protocol, learn a gene-specific but cell-type-invariant scaling factor from the shared cell types, and simulate data for the cell types missing in one protocol.

We note that the above analysis is only conducted for the genes’ mean expression levels. The difficulty of cross-protocol simulation is in fact even larger because realistic simulation requires the rescaling of the other distributional parameter(s) in a two-parameter distribution such as NB and ZIP or a three-parameter distribution such as ZINB. Existing work has provided extensive empirical evidence on the vast differences between protocols in terms of data characteristics [41, 80].

In applications 2 and 3, we have demonstrated how to use scDesign2 to guide experimental design and benchmark computational methods for the tasks of cell clustering and rare cell type detection. Note that in these analyses, the optimized sequencing depths and cell numbers are only applicable to the same experimental protocols and biological samples. Yet, this limitation does not disqualify scDesign2 as a useful tool to guide experimental design. For example, researchers usually perform a coarse-grained, low-budget experiment to obtain a preliminary dataset, and then they may use scDesign2 to guide the optimal design of the later, more refined experiment. As another example, if scRNA-seq data need to be collected from many individuals, researchers usually first perform a pilot study on a small number of individuals. Then they may train scDesign2 using the pilot data to guide the design of the subsequent, large-scale experiments. Moreover, in addition to guiding the experimental design, scDesign2 is useful as a general benchmarking tool for various experimental protocols and computational methods. For example, the analyses we performed in applications 2 and 3 are easily generalizable to other computational methods for a more comprehensive benchmarking.

Although we only use cell clustering and rare cell type detection to demonstrate scDesign2’s use in guiding experimental design and benchmarking computational methods, we want to emphasize that scDesign2 has broad applications beyond these two tasks. Inheriting the flexible and transparent modeling nature of our previous simulator scDesign, scDesign2 can also benchmark other computational analyses we have demonstrated in our scDesign paper [34], including differential gene expression analysis and cell dimensionality reduction. Moreover, beyond its role as a simulator, scDesign2 may benefit single-cell gene expression data analysis by providing its estimated parameters about gene expression and gene correlations. Here we discuss three potential directions. First, scDesign2 can assist differential gene expression analysis. Its estimated marginal distributions of individual genes in different cell types can be used to investigate more general patterns of differential expression (such as different variances and different zero proportions), in addition to comparing gene expression means between two groups of cells [100]. Second, its estimated gene correlation structures can be used to construct cell-type-specific gene networks [101] and incorporated into gene set enrichment analysis to enhance statistical power [102, 103]. Third, scDesign2 has the potential to improve the alignment of cells from multiple single-cell datasets [104]. Its estimated gene expression parameters can guide the calculation of cell type or cluster similarities between batches, and its estimated gene correlation structures can be used to align cell types or clusters across batches based on the similarity in gene correlation structures. [105].

## Methods

### The statistical framework of scDesign2

#### Fitting a generative model of single-cell gene expression count data with gene correlations

Given an scRNA-seq count matrix 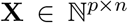 with *p* genes and *n* cells, we assume that the *n* cells belong to *K* cell types and that the cell memberships have been assigned by clustering, labeled by marker genes, or known in advance. (For input data without pre-defined cell types, our recommendation for cell clustering is in two subsections.) Our goal is to fit a parametric count model to characterize the joint distribution of genes’ counts in each cell type. For cell type *k*, We denote its number of cells by *n*^(*k*)^, its count sub-matrix by **X**^(*k*)^, and its set of model parameters by Θ^(*k*)^, *k* = 1,…,*K*. For simplicity of notation, we drop the superscript (*k*) in the following discussion about the generative model for one single cell type.

We denote *X_.j_* = (*X*_1*j*_,…, *X_pj_*)^⊤^ ∈ ℝ^*p*^ as a random *p*-dimensional gene count vector in cell *j, j* = 1,…,*n*. We denote its realization, i.e., the observed gene count vector as the *j*-th column in **X**, by *x_.j_* = (*x*_1*j*_,…, *x_pj_*)^⊤^. Jointly for the *p* genes, we assume that *X_.j_* independently follows a *p*-dimensional distribution *F*, which we will specify by a copula in the next paragraph. Marginally for each gene *i*, we assume that *X_ij_* independently follows a univariate count distribution *F_i_*. For example, if *F_i_*, is the ZINB distribution, we write 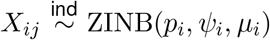, which can be interpreted as a hierarchical model: (1) 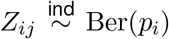 is a hidden latent variable indicating whether gene i drops out in cell *j*; (2) 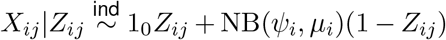, where 1_0_ indicates a point mass at 0. That is,

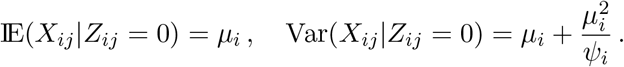

Note that the *Z_ij_*’s are unobserved and introduced only to describe the zero-inflation component. The Poisson, the zero-inflated Poisson (ZIP), and the negative binomial (NB) distributions are three special cases of the ZINB distribution, where *p_i_*, = 0 for Poisson and NB, and *ψ_i_*, = ∞ for Poisson and ZIP. From these four distributions, scDesign2 automatically chooses the one that best fits to gene *i*’s observed counts. Specifically, for the *i*-th row of **X**, *x_i._* = (*x*_*i*1_,…,*x_in_*)^⊤^, if its sample mean 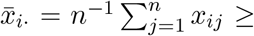 its sample variance 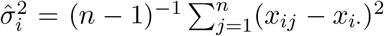, i.e., there is no over-dispersion, scDesign2 fits the Poisson and the ZIP distributions separately to *x_i._* by maximum likelihood estimation (MLE), and then performs a likelihood ratio test with 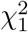 as the null distribution to determine if zero-inflation is significant, i.e., the ZIP distribution should be chosen over the Poisson distribution. Otherwise if there is over-dispersion, i.e., 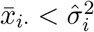, scDesign2 fits the NB and the ZINB distributions separately to *x_i._* by MLE and then performs a likelihood ratio test with 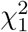 as the null distribution to determine if zero-inflation is significant, i.e., the ZINB distribution should be chosen over the NB distribution. The default significance level (i.e., p-value cutoff) for both tests is 0.05.

After estimating the marginal distributions of the *p* genes, i.e., *F*_1_,…,*F_p_*, scDesign2 uses a copula to model the joint *p*-dimensional distribution *F*. A copula is defined as a joint cumulative distribution function (CDF), *C*(·) : [0,1]^*p*^ → [0,1], which includesp uniform marginal distributions on [0,1]. That is, *C* is the CDF of a random vector *U* = (*U*_1_,…, *U_p_*)^⊤^ ∈ [0,1]^*p*^, with *U_i_* ~ Uniform[0,1], *i* = 1,…,*p*. For cell *j*’s gene count vector *X_.j_* ∈ ℝ^*p*^, although its *i*-th component *X_ij_* may not follow the Uniform[0,1] distribution, we can transform *X_ij_* by applying the marginal CDF *F_i_* so that *F_i_*(*X_ij_*) ∼ Uniform[0, 1]. This allows us to write the joint CDF *F* as

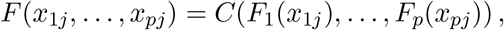

which is decomposable into the copula *C* and the marginal distributions *F*_1_,…, *F_p_*. Sklar’s theorem states that such a decomposition exists uniquely for any continuous distribution *F* [106]. If *F* is discrete in any dimension, the copula *C* still exists but may not be unique, i.e., not identifiable [107, 108]. To resolve this unidentifiability issue, scDesign2 uses the technique of distributional transform [109]: first draw *V_ij_* ~ Uniform[0,1] independently for *i* = 1,…,*p* and *j* = 1,…,*n*; second define *U_ij_* as

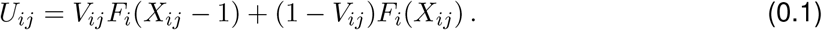

The effect of this transform is illustrated in Supplementary Fig. S47. Essentially, for a discrete random variable *X_ij_* with CDF *F_i_*, this transform distributes the non-zero probability mass *X_ij_* has at every value *x* uniformly to the interval [*x,x* + 1), thus transforming the discrete CDF *F_i_* to a continuous CDF 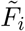 as

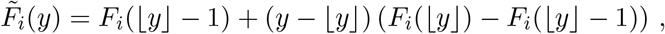

where ⌊*y*⌋ denotes the largest integer no greater than *y*.

With *V_ij_* and *X_ij_*, if we define

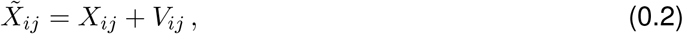

then the probability density function of 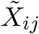 is

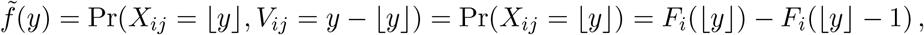

and the CDF of 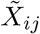 is

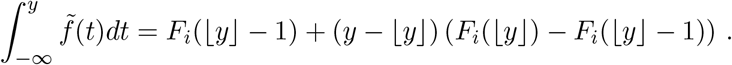

Hence, 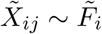; that is, the continuous random variable 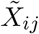 constructed from *X_ij_* and *V_ij_* follows 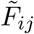. Defining 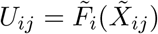, we have *U_ij_* ~ Uniform[0,1] and

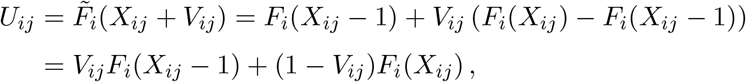

which is (0.1). This proves that *U_ij_* constructed by (0.1) follows Uniform[0,1] and is thus desirable.

After this transform, the CDF *F* of *X_.j_* is defined as the copula *C* of *U_.j_* = (*U*_1*j*_,…, *U_pj_*)^⊤^:

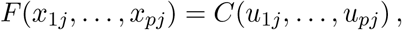

where (*u*_1*j*_,…, *u_pj_*)^⊤^ is a realization of (*U*_1*j*_,…, *U_pj_*)^⊤^. In scDesign2, we choose *C* as the Gaussian copula. Denoting by Φ the CDF of a standard Gaussian distribution, we define *F* as

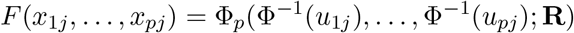

where Φ_*p*_(·; **R**) is the CDF of a *p*-dimensional Gaussian distribution with a zero mean vector and a covariance matrix that is equal to the correlation matrix **R**. If we denote *R_hl_* as the Gaussian copula correlation between genes *h* and *l*, i.e., the (*h,l*)-th entry of **R**, and *τ_hl_* as the Kendall’s tau between the same two genes on the original scale, i.e., *τ_hl_* = *τ*(*X_hj_,X_lh_*), then we have the following relationship [110, 111],

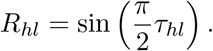

This relationship links the copula correlation with the Kendall’s tau of the two original variables, thus providing an interpretation of the copula correlation. It also suggests that **R** can be estimated by plugging the sample tau matrix into the above formula; however, this estimate of *R* may not always be positive semidefinite [112, 113]. Therefore, we use another procedure to estimate **R**.

Denote by 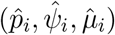 the estimated parameters of *F_i_*, which specify a fitted marginal distribution 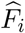. We sample 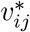 from Uniform[0,1] independently for *i* = 1,…,*p* and *j* = 1,…,*n*, and we calculate 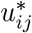 as

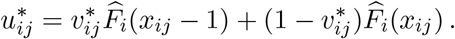

Then we estimate **R** by the sample covariance matrix 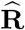 of 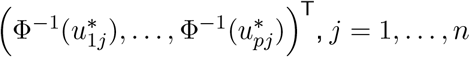, *j* = l,…,*n*.

As a side note, since this estimation procedure requires the random sampling of 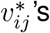, it introduces additional randomness into the estimation of **R**; that is, 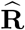 is not a deterministic function of data. However, this additional randomness has a negligible effect on the synthetic data. As demonstrated in Supplementary Fig. S48, the gene correlation matrices estimated from synthetic data generated by scDesign2, with 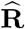 estimated under two different random samples of 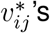, are very similar to each other.

To summarize, scDesign2 first estimate the marginal distributions *F*_1_,…,*F_p_* as 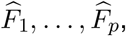, each of which may be a fitted Poisson, ZIP, NB, or ZINB distribution. Then scDesign2 calculates 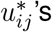 as described above and estimates a *p* × *p* covariance matrix as 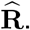 Finally, scDesign2 estimates the *p*-dimensional joint distribution *F* as

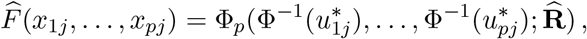

whose estimated model parameters are 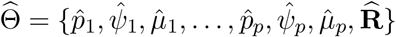.

As a practical note, since the data matrix **X** typically contains tens of thousands of genes, if the sample size, i.e., the number of cells is not large enough, the estimation of the copula correlation matrix can be problematic [91]. Moreover, many genes are too lowly expressed to be detected in scRNA-seq data, making their correlations uninteresting to estimate. For these two reasons, we argue that the copula correlations should only be estimated for a subset of moderately to highly expressed genes.

In Supplementary Figs. S42 and S43, we analyze how *n* (the sample size, i.e., the number of cells) and p (the number of top expressed genes included) affect the estimation of the copula correlation matrix. We use two example datasets: the stem cell data generated by the 10x Genomics protocol and the dendrocyte (subtype 1) data generated by the Smart-Seq2 protocol. For each dataset, we extract the fitted Gaussian copula model for the top 2000 genes with the highest mean expression levels, and we use this model as the ground truth model to generate 1000 samples with a varying *n*. Then we estimate the copula correlation matrix of a varying *p* from each sample. For computational efficiency, we use the plug-in estimation method based on sample tau values: 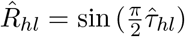. Finally, we calculate the mean squared error (MSE) between the estimated copula correlations and the true copula correlations. That is, for each *n* and *p*, we have 1000 MSE values.

In Supplementary Figs. S42 and S43, from panel (a), we can see that MSEs decrease as *n* increases. From panel (b), we can see that MSEs increase as *p* increases, i.e., more lowly expressed genes are included. To ease the interpretation of the results, we mark three horizontal lines at MSE = 0.09, 0.04, and 0.01 to represent three levels of estimation quality. On the scale of correlation values, these three levels indicate that on average the estimated values are within ±0.3, ±0.2, and ±0.1 of the true values. The results suggest that to reach the ±0.3 level of estimation quality, a reasonable choice of *n* is at least 20, and the top 1000 highly expressed genes can be included. To reach the ±0.2 level, a reasonable choice of *n* is at least 50, and the top 1500 highly expressed genes can be included. For *n* = 100, the ±0.1 level can be reached for the top 100-200 highly expressed genes, and even the error level for the top 2000 is close to this level. The results confirm that sample size is not a concern for single-cell data because most cell types contain at least a hundred cells that can be measured by current protocols.

In the implementation of the `scDesign2 R` package, before fitting the above generative model for each cell type, scDesign2 partitions the genes into three groups: the first group containing genes with zero proportions less than a cutoff (default 0.8, but can be changed according to the discussion above), the second group containing genes with zero proportions between the cutoff and (*n*–2)/*n*, where *n* is the number of cells, and the last group including the remaining genes, i.e., genes expressed in fewer than three cells. For the first group, scDesign2 fits the above generative model jointly for its genes. For the second group, scDesign2 fits a marginal distribution for each individual gene. For the last group, scDesign2 only generates zero counts for all its genes.

#### Generation of synthetic single-cell gene expression count data

To generate synthetic scRNA-seq data for *K* cell types, scDesign2 first estimates the proportions of *K* cell types from the real scRNA-seq count matrix **X**, for which we denote the number of reads mapped to the *n*^(*k*)^ cells of type *k* as *N*^(*k*)^, and the total number of reads mapped to all the *n* cells as 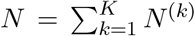. Denoting the cell type proportions as *π* = (*π*^(1)^,…,*π*^(*K*)^)^⊤^ such that 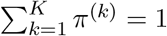, scDesign2 estimates them by 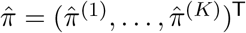, where

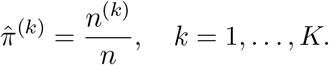

We denote the synthetic scRNA-seq data to be generated as **X**′, which contains *n*′ cells and *N*′ expected number of reads, with *n*′ and *N*′ as user-specified input parameters of scDesign2. Denoting the number of synthetic cells of type *k* as *n*^(*k*)′^, scDesign2 draws the numbers of synthetic cells of all *K* cell types from a multinomial distribution, i.e., 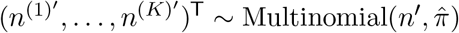. Then given *n*^(*k*)′^, the expected number of reads assigned to cell type *k* in **X**′ should be

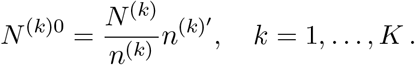

However, given the constraint that the expected total number of reads in **X**′ is *N*′, we need to rescale *N*^(*k*)0^ to

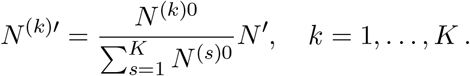

As a result, the scaling factor is

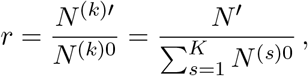

which does not depend on the cell type, and scDesign2 uses this scaling factor to rescale the mean parameter of every gene.

Given the fitted generative model 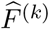 for cell type *k* with parameters

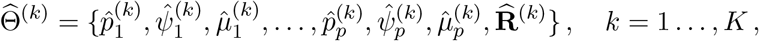

and the scaling factor *r*, scDesign2 generates *n*^(*k*)′^ synthetic cells from a rescaled model 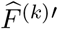, which is defined by parameters

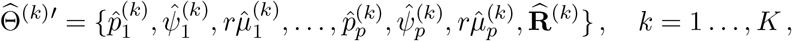

Concretely, how the data generation works is that scDesign2 first draws *n*^(*k*)′^ vectors, denoted as 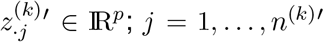, independently from 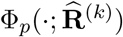. Then scDesign2 converts 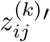 to 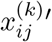 by setting 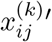 to be the 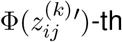 quantile of 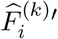 i.e., 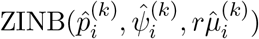 (including the Poisson, ZIP, and NB distributions as special cases), *i* = 1,…,*p*. Finally, scDesign2 outputs the synthetic count matrix **X**′ = [**X**^(1)^′ ⋯ **X**^(*K*)^′], where 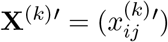 is a *p* × *n*^(*k*)′^ matrix for cell type *k*.

Note that the synthetic count matrix **X**′ does not contain exactly *N*′ reads; rather, *N*′ is the expected total number of reads. We think this setting mimics a real sequencing experiment, where the total number of sequenced reads would not be exactly the same as the preset sequencing depth *N*′ due to experimental randomness.

#### Recommendation for cell clustering when input data do not have labelled cell types

If users would like to train scDesign2 on a gene-by-cell count matrix without cell type labels, a necessary preceding step is cell clustering. We recommend users to choose a state-of-the-art cell clustering method such as Seurat and SC3. For the resulting clusters, we recommend users to visualize them by t-SNE or UMAP and use a goodness-of-fit measure (e.g., Pearson’s chi-square statistic and ROGUE score [81]) to check whether each gene approximately follows a NB or ZINB distribution in a cell cluster. This check will guide users to decide on an appropriate number of cell clusters in a data-driven way.

### The scDesign2 variant without copula

The only difference between this variant “w/o copula” and scDesign2 is that this variant assumes the p genes to have independent marginal distributions *F*_1_,…,*F_p_*. The fitting of the p marginal distributions and the generation of synthetic data is the same as those in scDesign2.

### Existing simulators

- **scDesign**: The R package scDesign version 1.0.0 is used for the analysis.
- **scGAN**: This method is executed from this github repository https://github.com/imsb-uke/scGAN, downloaded around March 29, 2020.
- **splat, splat simple, kersplat**: These methods are executed from the R packge splatter version 1.10.1.
- **SPARSim**: The R package SPARSim version 0.9.5 is used for the analysis.
- **SymSim**: The R package SymSim version 0.0.0.9000 is used for the analysis.
- **ZINB-WaVE**: The ZINB-WaVE method is used from the wrapper functions in the R package splatter version 1.10.1.
- **scDesign**: The R package scDesign version 1.0.0 is used for the analysis.

### Dimensionality reduction methods

- **t-SNE**: The R package Rtsne version 0.15 is used for generating t-SNE plots. The function Rtsne is used, with all parameters set to default, except check duplicate = FALSE and perplexity is changed from the default value of 30 to one third of the sample size when the sample size (total number of cells) is less than 90.
- **PCA**: The R function prcomp() is used for generating PCA plots, with parameters set as default.

### Cell clustering methods

- **Seurat**: The Seurat clustering method is executed by the following instruction in this tutorial https://satijalab.org/seurat/v3.2/pbmc3k_tutorial.html. R package Seurat version 3.1.5 is used for the analysis.
- **SC3**: The SC3 clustering method is executed by the following instruction in this tutorial https://www.bioconductor.org/packages/release/bioc/vignettes/SC3/inst/doc/SC3.html. R package SC3 version 1.14.0 is used for the analysis.

### Rare cell type detection methods

- **FiRE**: The FiRE method is executed by the following instruction in this tutorial https://github.com/princethewinner/FiRE. R package FiRE version 1.0 is used for the analysis.
- **GiniClust2**: This method is executed from this github repository https://github.com/dtsoucas/GiniClust2 downloaded around March 4, 2020. It is executed based on the reference manual in this repository, except no cells are filtered.

### Datasets

- **10x Genomics**: The 10x Genomics dataset measures the mouse intestinal epithelial tissue [76]. The raw count dataset is downloaded from Gene Expression Omnibus (GEO) with accession number GSE92332. Data for cell types Stem, Goblet, Tuft, Transit Amplifying Early (TA Early), Enterocyte Progenitor and Enterocyte Progenitor Early were selected for analysis. Spike-in RNA counts were filtered. The resulting count matrix contains 15962 genes and 3793 cells.
- **CEL-Seq2**: The CEL-Seq2 dataset measures the human pancreas [77]. The raw count dataset is downloaded from GEO with accession number GSE85241. Data for cell types alpha, beta, acinar, delta, duct, endothelial, mesenchymal and pancreatic polypeptide cell (pp) were selected for analysis. Spike-in RNA counts were filtered. The resulting count matrix contains 19049 genes and 2279 cells.
- **Fluidigm C1**: The Fluidigm C1 dataset measures human brain cells [78]. The raw count dataset is downloaded from GEO with accession number GSE67835. Data for cell types astrocytes, endothelial, fetal quiescent, hybrid neurons, oligodendrocytes and oligodendrocyte precursor cell (OPC) were selected for analysis. The resulting count matrix contains 22088 genes and 317 cells.
- **Smart-Seq2**: The Smart-Seq2 dataset measures human blood dendritic cells [79]. The raw count dataset is downloaded from GEO with accession number GSE94820. Data for dendrocyte subtypes 1–6 and monocyte subtypes 1–4 were selected for analysis. Spike-in RNA counts were filtered. The resulting count matrix contains 26586 genes and 1078 cells.
- **MERFISH**: The MERFISH dataset measures the mouse hypothalamic preoptic region [85]. The raw count dataset is downloaded from Dryad (https://datadryad.org/stash/dataset/doi:10.5061/dryad.8t8s248). It contains 155 genes and 6412 cells. Cell subtypes are combined into cell types, e.g., “Endothelial 1” and “Endothelial 2” are combined as “Endothelial”, resulting in nine cell types in total.
- **pciSeq**: The pciSeq dataset measures the mouse hippocampal area CA1 [86]. The raw data “cθlls_lθft_CA1 _3-1” are downloaded from https://su.figshare.com/articles/pciSeq_files_in_csv_format/10318610/1. Gene expression values are rounded as integers, and cell subtypes are combined into cell types, e.g., “Astro.1” to “Astro.5” are combined as “Astro”. The cell type “Zero” is removed as it contains cells with almost no genes expressed, so seven cell types are retained. The processed data contain 84 genes and 2253 cells.

## Software and code

The `scDesign2 R` package is available at https://github.com/JSB-UCLA/scDesign2. The source code and data for reproducing the results are available at https://doi.org/10.5281/zenodo.4011311.

## Competing interests

The authors declare no competing interests.

## Acknowledgements

The authors would like to thank Dr. Roy Wollman and his Ph.D. student Zach Hemminger for bringing our attention to MERFISH and pciSeq data. The authors also appreciate the comments and feedback from the members of the Junction of Statistics and Biology at UCLA (http://jsb.ucla.edu).

## Funding

This work was supported by the following grants: National Science Foundation DBI-1846216, NIH/NIGMS R01GM120507, Johnson & Johnson WiSTEM2D Award, Sloan Research Fellowship, and UCLA David Geffen School of Medicine W.M. Keck Foundation Junior Faculty Award (to J.J.L); Rutgers School of Public Health Pilot Grant and NJ ACTS BERD Mini-Methods Grant (to W.V.L).

## Footnotes

1 ZINB-WaVE was not proposed as a simulator in its original publication [38] but was later implemented as a simulator in the splatter package [65].

2 A quote from the SERGIO paper [72]: “It is worth noting here that several existing single-cell expression simulators employ a probabilistic model whose parameters are directly estimated from a real dataset and then sample synthetic data from the model. This approach is not feasible in SERGIO since the true GRN underlying the real dataset is unknown and notoriously hard to reconstruct, and the explicit use of a GRN is a crucial distinguishing feature of SERGIO. As such, SERGIO uses a randomly generated GRN to first synthesize clean expression data and uses the real dataset only in the second phase, to determine the extent of technical noise to add to the clean data.”

3 The training of scGAN takes 1-2 days (with NVIDIA GeFore GTX 2080 Ti GPU) on 255 cells and 15926 genes, in contrast to the other simulators that take at most minutes to train with CPU.

## Notes

### Competing Interest Statement

The authors have declared no competing interest.

### Summary of Updates

1. We have included the ROGUE score analysis as a preprocessing step before the model fitting of scDesign2. Users can assess the homogeneity of every cell type or cell cluster, before using scDesign2 to fit cell-type-specific models. We have summarized the ROGUE score analysis results in Fig. 5. 2. We have shown in Fig. 3b, S6b-S16b, and S17-S28 that scDesign2 outperforms existing simulators in preserving correlation statistics of the top highly expressed genes, which are key to the resemblance between synthetic data and real data. The revised text is in the Results section on Pages 6-7. 3. We have added explanations about the distributional transform used in the estimation of copula correlations. The new text is in the Methods section on Pages 20-21, and a new figure Supplementary Fig. S47 has been added for illustration purposes. 4. We have found an analytical relationship between Kendall's tau and the Gaussian copula correlation from the literature. This relationship offers an interpretation of the copula correlation matrix in the scDesign2 model and is summarized in the Methods section on Page 21. 5. We have designed simulation studies to analyze the effects of n (the sample size, i.e., the number of cells) and p (the number of genes) on the estimation accuracy of the copula correlation matrix. Based on the results, we have added practical suggestions for scDesign2 users and outlined future improvement directions for the scDesign2 method. We have summarized the analysis results in Supplementary Figs. S42 and S43, and we have added text to the Discussion section on Pages 15-16 and the Methods section on Page 22. 6. We have included the median integration LISI (miLISI) as a quantitative metric to assess how well the synthetic data mix with the real data in Figs. 4, 6, and S29-S32. The analysis is summarized in the Results section on Pages 7-8. 7. We have added a scenario for the experimental design application of scDesign2: with the per-cell sequencing depth fixed, the total number of cells sequenced changes. The results are summarized in the Results section on Pages 12 and 14, and Figs. 9 and 12 and Figs. S35, S38, and S41. 8. We have demonstrated and explained why the current implementation of scDesign2 cannot be directly used for cross-protocol data simulation. The results are summarized in the Discussion section on Pages 17-18 and in Figs. S44-S46.

