## Supplementary for "scDesign2: a transparent simulator that generates high-fidelity single-cell gene expression count data with gene correlations captured"

### Supplementary Tables

**Table S1: Summary of the sample size ($n$) and the number of genes included for copula correlation estimation ($p$), for each of the 12 datasets used for the benchmarking of simulators.**

| protocol | cell type | number of cells used for model fitting ($n$) | number of genes included for copula correlation estimation ($p$) |
|---|---|---|---|
| 10x Genomics | goblet | 255 | 3022 |
| 10x Genomics | stem | 634 | 2465 |
| 10x Genomics | tuft | 83 | 2981 |
| CEL-Seq2 | alpha | 426 | 8285 |
| CEL-Seq2 | beta | 233 | 8498 |
| CEL-Seq2 | acinar | 134 | 9612 |
| Fluidigm C1 | astrocytes | 25 | 7574 |
| Fluidigm C1 | neurons | 61 | 9440 |
| Fluidigm C1 | oligodendrodytes | 19 | 7232 |
| Smart-Seq2 | dendrocyte 1 | 83 | 8880 |
| Smart-Seq2 | dendrocyte 2 | 47 | 8483 |
| Smart-Seq2 | monocyte 2 | 61 | 8459 |

#### Supplementary Figures

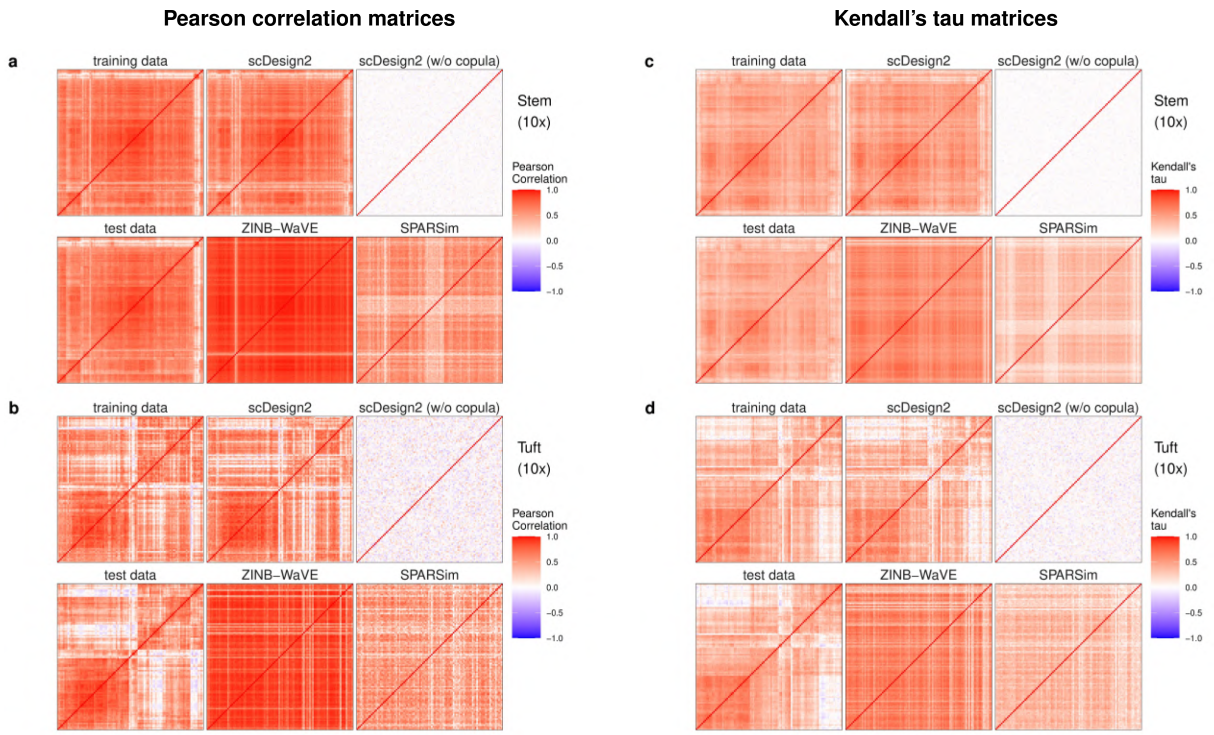


**Figure S1: Heatmaps of gene correlation matrices estimated from real data and synthetic data generated by scDesign2, its variant without copula, ZINB-WaVE, and SPARSim.** (a)-(b) Pearson correlation matrices; (c)-(d) Kendall's tau matrices. In (a) and (c), training and test data contain stem cells measured by 10x Genomics [76]; In (b) and (d), training and test data contain tuft cells measured by 10x Genomics [76]. For each cell type, the Pearson correlation matrices and Kendall's tau matrices are shown for the 100 genes with the highest mean expression values in the test data; the rows and columns (i.e., genes) of all the matrices are ordered by the complete-linkage hierarchical clustering of genes (using Pearson correlation as the similarity in (a)-(b) and Kendall's tau in (c)-(d)) in the test data. We find that the correlation matrices estimated from the sythetic data generated by scDesign2 most resemble those of training and test data.

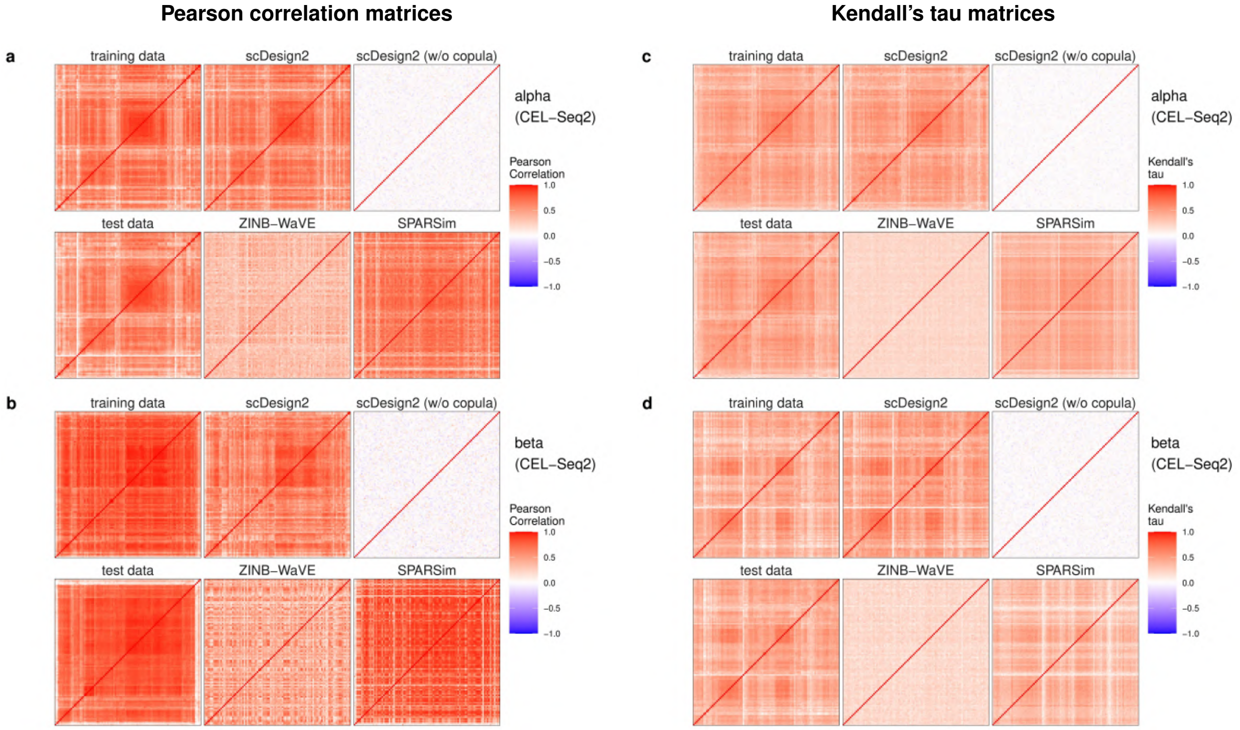


**Figure S2: Heatmaps of gene correlation matrices estimated from real data and synthetic data generated by scDesign2, its variant without copula, ZINB-WaVE, and SPARSim.** (a)-(b) Pearson correlation matrices; (c)-(d) Kendall's tau matrices. In (a) and (c), training and test data contain alpha cells measured by CEL-Seq2 [77]; In (b) and (d), training and test data contain beta cells measured by CEL-Seq2 [77]. For each cell type, the Pearson correlation matrices and Kendall's tau matrices are shown for the 100 genes with the highest mean expression values in the test data; the rows and columns (i.e., genes) of all the matrices are ordered by the complete-linkage hierarchical clustering of genes (using Pearson correlation as the similarity in (a)-(b) and Kendall's tau in (c)-(d)) in the test data. We find that the correlation matrices estimated from the sythetic data generated by scDesign2 most resemble those of training and test data.

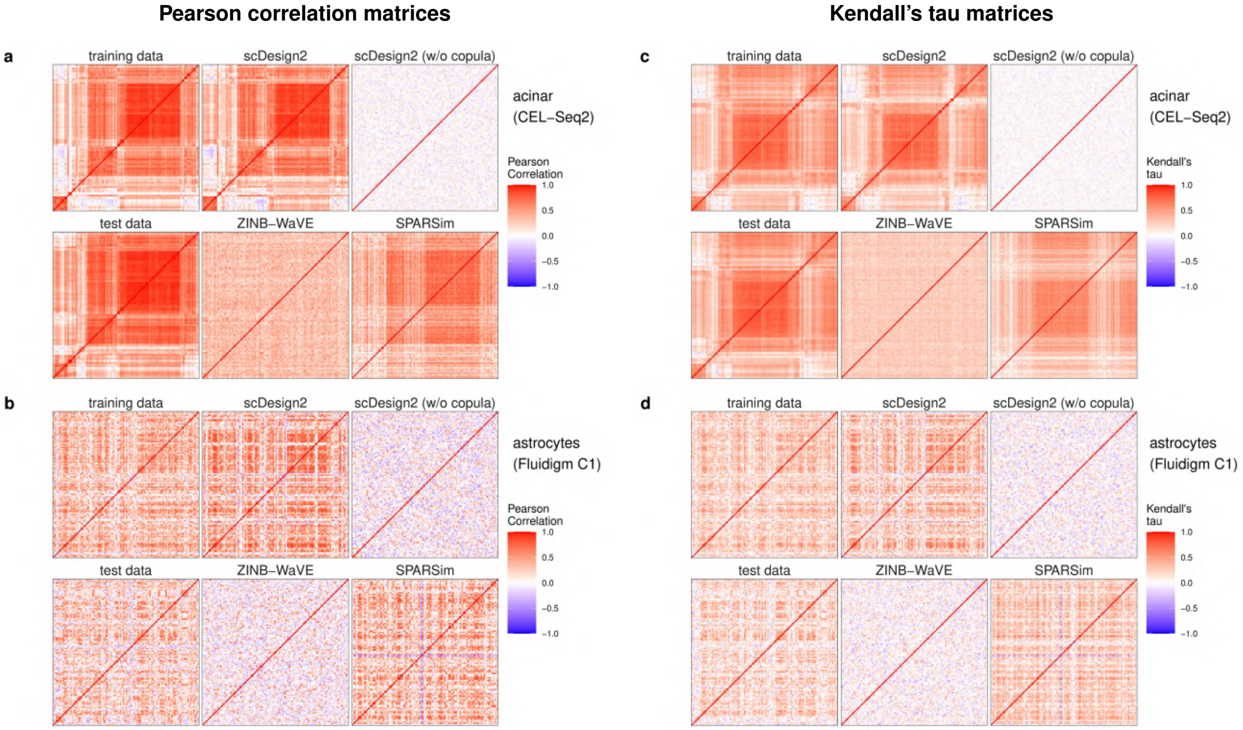


**Figure S3: Heatmaps of gene correlation matrices estimated from real data and synthetic data generated by scDesign2, its variant without copula, ZINB-WaVE, and SPARSim.** (a)-(b) Pearson correlation matrices; (c)-(d) Kendall's tau matrices. In (a) and (c), training and test data contain acinar cells measured by CEL-Seq2 [77]; In (b) and (d), training and test data contain astrocytes measured by Fluidigm C1 [78]. For each cell type, the Pearson correlation matrices and Kendall's tau matrices are shown for the 100 genes with the highest mean expression values in the test data; the rows and columns (i.e., genes) of all the matrices are ordered by the complete-linkage hierarchical clustering of genes (using Pearson correlation as the similarity in (a)-(b) and Kendall's tau in (c)-(d)) in the test data. We find that the correlation matrices estimated from the sythetic data generated by scDesign2 most resemble those of training and test data.

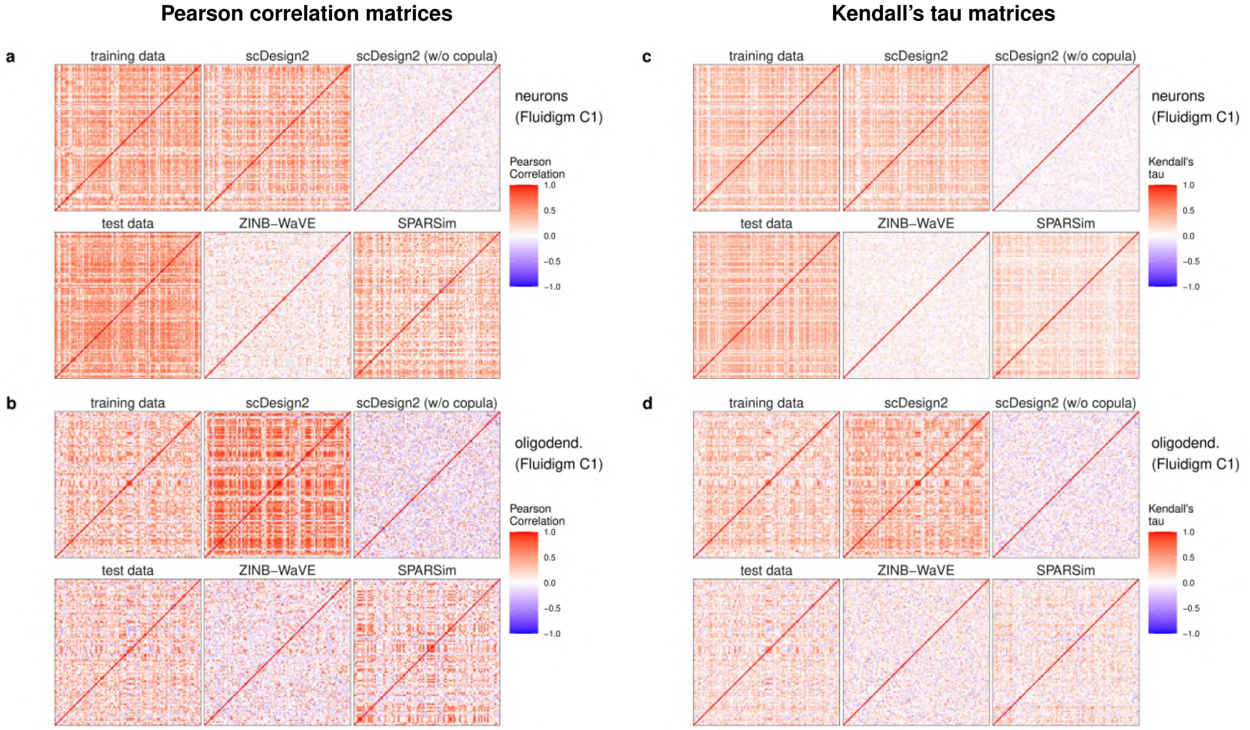


**Figure S4: Heatmaps of gene correlation matrices estimated from real data and synthetic data generated by scDesign2, its variant without copula, ZINB-WaVE, and SPARSim.** (a)-(b) Pearson correlation matrices; (c)-(d) Kendall's tau matrices. In (a) and (c), training and test data contain neurons measured by Fluidigm C1 [78]; In (b) and (d), training and test data contain oligodendrocytes measured by Fluidigm C1 [78]. For each cell type, the Pearson correlation matrices and Kendall's tau matrices are shown for the 100 genes with the highest mean expression values in the test data; the rows and columns (i.e., genes) of all the matrices are ordered by the complete-linkage hierarchical clustering of genes (using Pearson correlation as the similarity in (a)-(b) and Kendall's tau in (c)-(d)) in the test data. We find that the correlation matrices estimated from the sythetic data generated by scDesign2 most resemble those of training and test data.

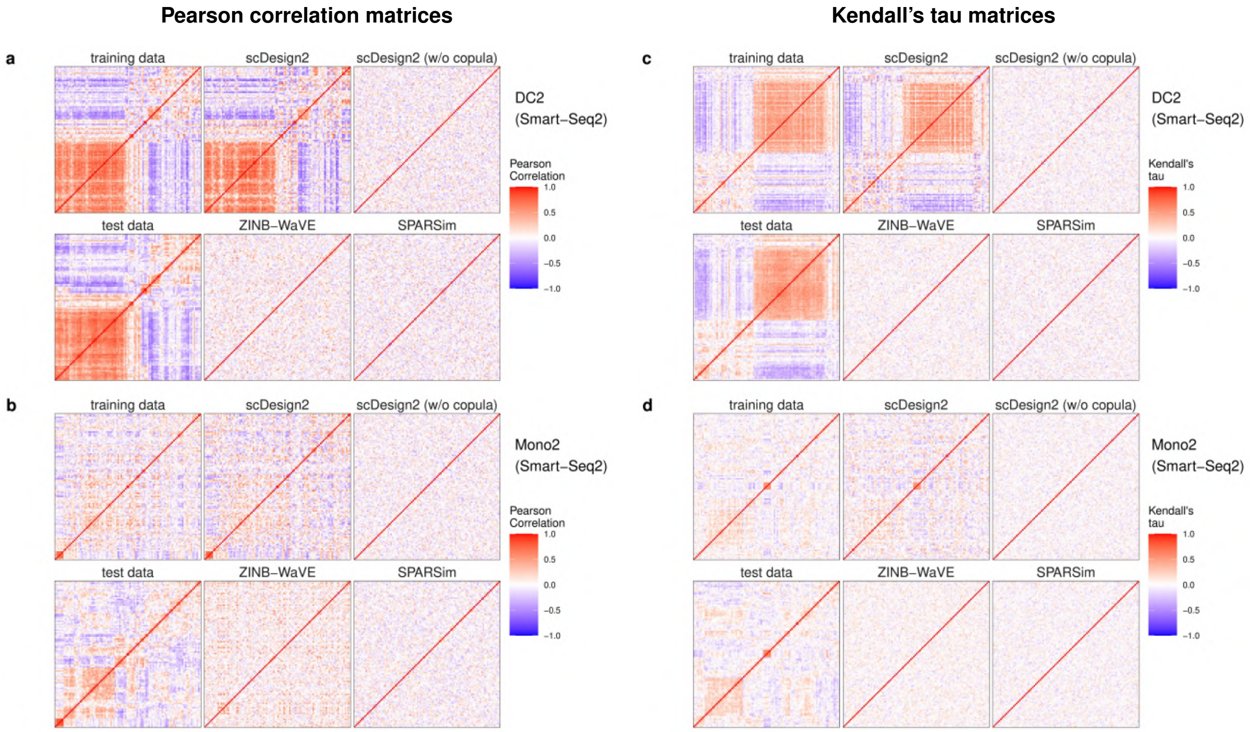


**Figure S5: Heatmaps of gene correlation matrices estimated from real data and synthetic data generated by scDesign2, its variant without copula, ZINB-WaVE, and SPARSim.** (a)-(b) Pearson correlation matrices; (c)-(d) Kendall's tau matrices. In (a) and (c), training and test data contain cells of dendrocytes subtype 2 (DC2) measured by Smart-Seq2 [79]; In (b) and (d), training and test data contain cells of monocytes subtype 2 (Mono2) measured by Smart-Seq2 [79]. For each cell type, the Pearson correlation matrices and Kendall's tau matrices are shown for the 100 genes with the highest mean expression values in the test data; the rows and columns (i.e., genes) of all the matrices are ordered by the complete-linkage hierarchical clustering of genes (using Pearson correlation as the similarity in (a)-(b) and Kendall's tau in (c)-(d)) in the test data. We find that the correlation matrices estimated from the sythetic data generated by scDesign2 most resemble those of training and test data.

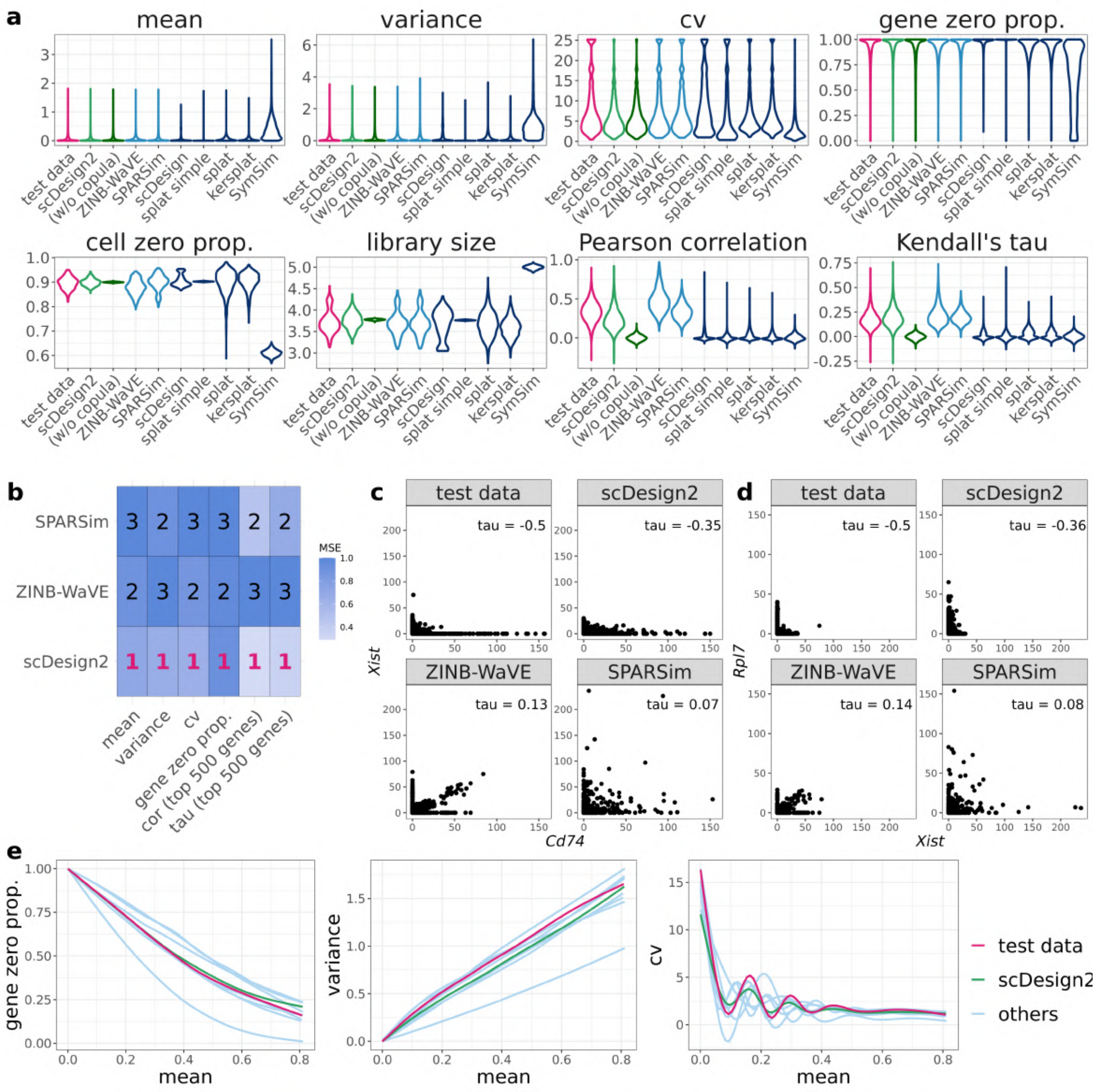


**Figure S6: Benchmarking scDesign2 against its variant without copula and seven existing scRNA-seq simulators for generating stem cells measured by 10x Genomics.** (a) Distributions of eight summary statistics (gene-wise expression mean, variance, coefficient of variation (cv), and zero proportion; cell-wise zero proportion and library size; gene-pair-wise Pearson correlation and Kendall's tau) are plotted based on the real data (test data unused for training simulators) and the synthetic data generated by scDesign2, scDesign2 without copula (w/o copula), ZINB-WaVE, SPARSim, scDesign, three variants of the splatter package (splat simple, splat, and kersplat), and SymSim. (b) Ranking (with 1 being the best-performing method) of scDesign2, ZINB-WaVE, and SPARSim, the only three methods that preserve genes, in terms of the mean-squared error (MSE) of each of six summary statistics (four gene-wise and two gene-pair-wise) between the statistic values in the real data and the synthetic data generated by each simulator. Note that the color scale shows the normalized MSE: for each statistic (column in the table), the normalized MSEs are the MSEs divided by the largest MSE of that statistic. scDesign2 is ranked the top for six out of the six statistics. For the two gene-pair-wise statistics, we focus on the top 500 highly expressed genes, because as analyzed in the text, they are more meaningful, both biologically and statistically, than the correlations of the lowly expressed genes. (c)-(d) Scatterplots of two example gene pairs—*Xist* vs. *CD74* and *Rpl7* vs. *Xist*—based on the real data and the synthetic data generated by scDesign2, ZINB-WaVE, and SPARSim. Only scDesign2 captures the negative gene correlations in the real data. (e) Smoothed relationships between three pairs of gene-wise statistics (zero proportion vs. mean, variance vs. mean, and cv vs. mean) across all genes (curves plotted by the R function `geom_smooth()`) in the real data and the synthetic data generated by scDesign2 and the seven existing simulators (others). Note that ZINB-WaVE and SymSim filter out certain genes when simulating new data; Pearson correlation and Kendall's tau are only calculated between the genes whose zero proportions are less than 50%; gene-wise mean and variance and cell-wise library size are transformed to the $\log_{10}(1+x)$ scale (where $x$ represents a statistic's value).

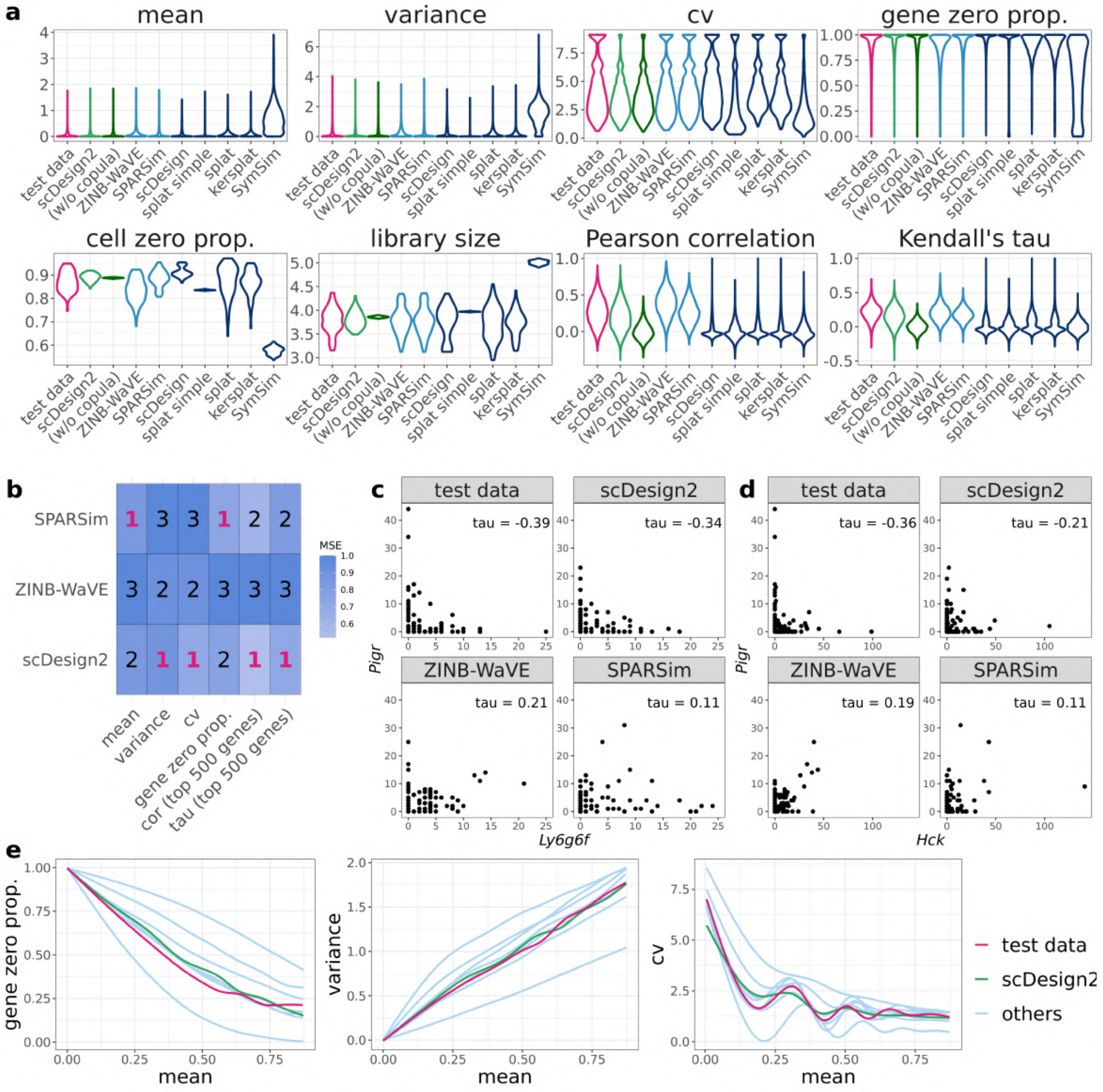


**Figure S7: Benchmarking scDesign2 against its variant without copula and seven existing scRNA-seq simulators for generating tuft cells measured by 10x Genomics.** (a) Distributions of eight summary statistics (gene-wise expression mean, variance, coefficient of variation (cv), and zero proportion; cell-wise zero proportion and library size; gene-pair-wise Pearson correlation and Kendall's tau) are plotted based on the real data (test data unused for training simulators) and the synthetic data generated by scDesign2, scDesign2 without copula (w/o copula), ZINB-WaVE, SPARSim, scDesign, three variants of the splatter package (splat simple, splat, and kersplat), and SymSim. (b) Ranking (with 1 being the best-performing method) of scDesign2, ZINB-WaVE, and SPARSim, the only three methods that preserve genes, in terms of the mean-squared error (MSE) of each of six summary statistics (four gene-wise and two gene-pair-wise) between the statistic values in the real data and the synthetic data generated by each simulator. Note that the color scale shows the normalized MSE: for each statistic (column in the table), the normalized MSEs are the MSEs divided by the largest MSE of that statistic. scDesign is ranked the top for four out of the six statistics. For the two gene-pair-wise statistics, we focus on the top 500 highly expressed genes, because as analyzed in the text, they are more meaningful, both biologically and statistically, than the correlations of the lowly expressed genes. (c)-(d) Scatterplots of two example gene pairs—*Ly6g6f* vs. *Pigr* and *Hck* vs. *Pigr*—based on the real data and the synthetic data generated by scDesign2, ZINB-WaVE, and SPARSim. Only scDesign2 captures the negative gene correlations in the real data. (e) Smoothed relationships between three pairs of gene-wise statistics (zero proportion vs. mean, variance vs. mean, and cv vs. mean) across all genes (curves plotted by the R function `geom_smooth()`) in the real data and the synthetic data generated by scDesign2 and the seven existing simulators (others). Note that ZINB-WaVE and SymSim filter out certain genes when simulating new data; Pearson correlation and Kendall's tau are only calculated between the genes whose zero proportions are less than 50%; gene-wise mean and variance and cell-wise library size are transformed to the $\log_{10}(1+x)$ scale (where $x$ represents a statistic's value).

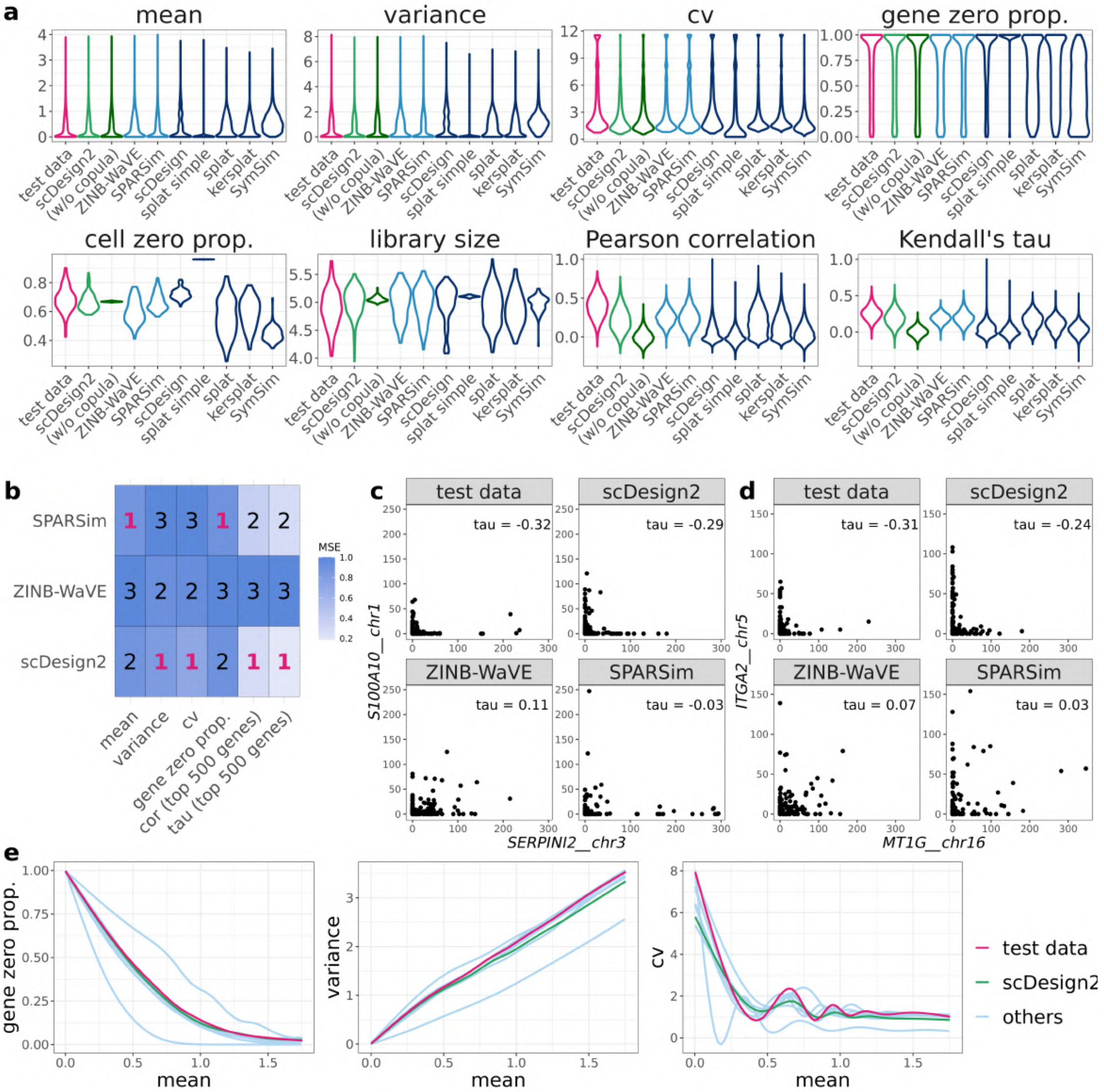


**Figure S8: Benchmarking scDesign2 against its variant without copula and seven existing scRNA-seq simulators for generating acinar cells measured by CEL-Seq2.** (a) Distributions of eight summary statistics (gene-wise expression mean, variance, coefficient of variation (cv), and zero proportion; cell-wise zero proportion and library size; gene-pair-wise Pearson correlation and Kendall's tau) are plotted based on the real data (test data unused for training simulators) and the synthetic data generated by scDesign2, scDesign2 without copula (w/o copula), ZINB-WaVE, SPARSim, scDesign, three variants of the splatter package (splat simple, splat, and kersplat), and SymSim. (b) Ranking (with 1 being the best-performing method) of scDesign2, ZINB-WaVE, and SPARSim, the only three methods that preserve genes, in terms of the mean-squared error (MSE) of each of six summary statistics (four gene-wise and two gene-pair-wise) between the statistic values in the real data and the synthetic data generated by each simulator. Note that the color scale shows the normalized MSE: for each statistic (column in the table), the normalized MSEs are the MSEs divided by the largest MSE of that statistic. scDesign is ranked the top for four out of the six statistics. For the two gene-pair-wise statistics, we focus on the top 500 highly expressed genes, because as analyzed in the text, they are more meaningful, both biologically and statistically, than the correlations of the lowly expressed genes. (c)-(d) Scatterplots of two example gene pairs—*SERPINI2* vs. *S100A10* and *MT1G* vs. *ITGA2*—based on the real data and the synthetic data generated by scDesign2, ZINB-WaVE, and SPARSim. (e) Smoothed relationships between three pairs of gene-wise statistics (zero proportion vs. mean, variance vs. mean, and cv vs. mean) across all genes (curves plotted by the R function `geom_smooth()`) in the real data and the synthetic data generated by scDesign2 and the seven existing simulators (others). Note that ZINB-WaVE and SymSim filter out certain genes when simulating new data; Pearson correlation and Kendall's tau are only calculated between the genes whose zero proportions are less than 50%; gene-wise mean and variance and cell-wise library size are transformed to the $\log_{10}(1+x)$ scale (where $x$ represents a statistic's value).

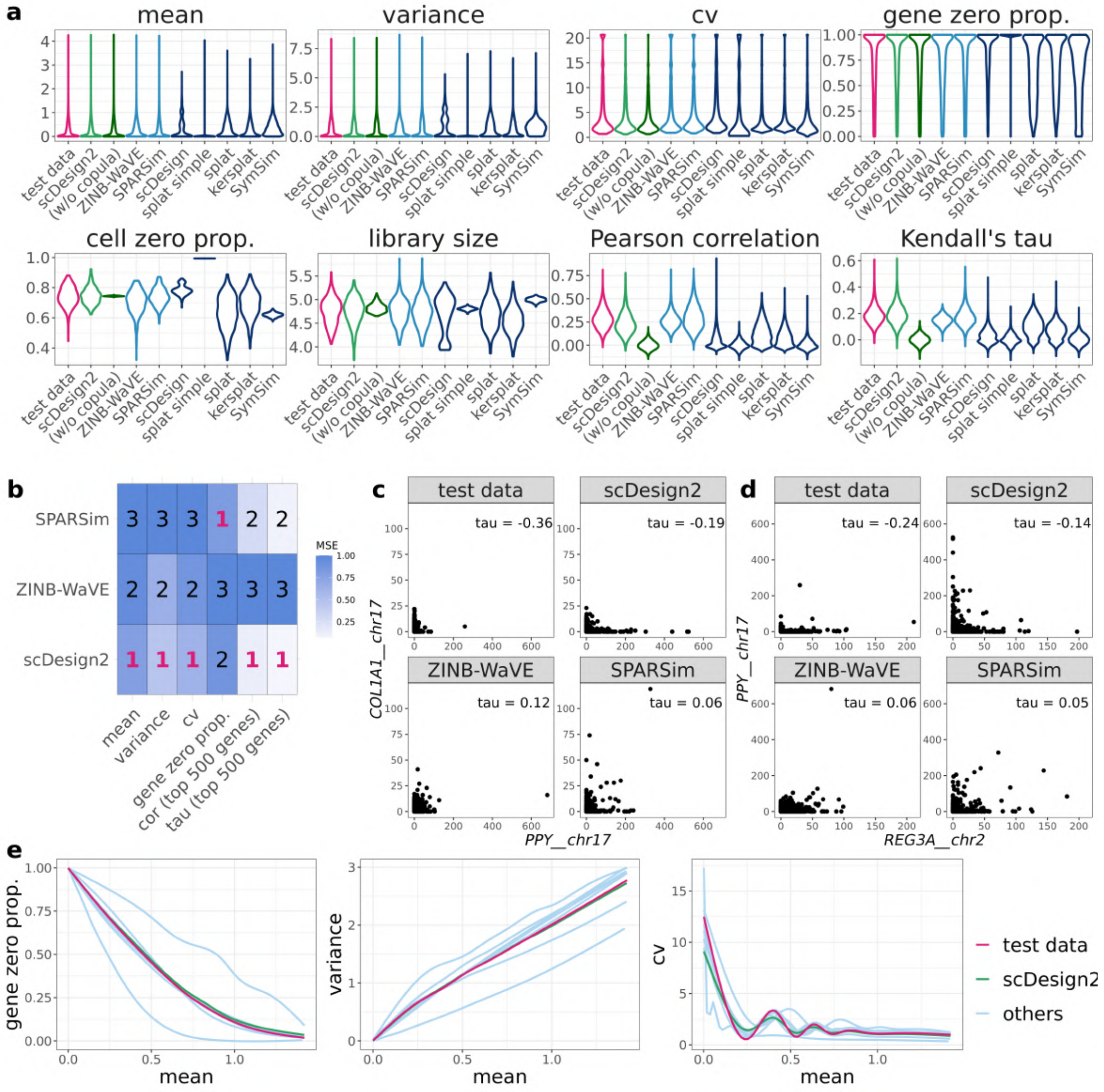


**Figure S9: Benchmarking scDesign2 against its variant without copula and seven existing scRNA-seq simulators for generating alpha cells measured by CEL-Seq2.** (a) Distributions of eight summary statistics (gene-wise expression mean, variance, coefficient of variation (cv), and zero proportion; cell-wise zero proportion and library size; gene-pair-wise Pearson correlation and Kendall's tau) are plotted based on the real data (test data unused for training simulators) and the synthetic data generated by scDesign2, scDesign2 without copula (w/o copula), ZINB-WaVE, SPARSim, scDesign, three variants of the splatter package (splat simple, splat, and kersplat), and SymSim. (b) Ranking (with 1 being the best-performing method) of scDesign2, ZINB-WaVE, and SPARSim, the only three methods that preserve genes, in terms of the mean-squared error (MSE) of each of six summary statistics (four gene-wise and two gene-pair-wise) between the statistic values in the real data and the synthetic data generated by each simulator. Note that the color scale shows the normalized MSE: for each statistic (column in the table), the normalized MSEs are the MSEs divided by the largest MSE of that statistic. scDesign is ranked the top for five out of the six statistics. For the two gene-pair-wise statistics, we focus on the top 500 highly expressed genes, because as analyzed in the text, they are more meaningful, both biologically and statistically, than the correlations of the lowly expressed genes. (c)-(d) Scatterplots of two example gene pairs—*PPY* vs. *COL1A1* and *REG3A* vs. *PPY*—based on the real data and the synthetic data generated by scDesign2, ZINB-WaVE, and SPARSim. Only scDesign2 captures the negative gene correlations in the real data. (e) Smoothed relationships between three pairs of gene-wise statistics (zero proportion vs. mean, variance vs. mean, and cv vs. mean) across all genes (curves plotted by the R function `geom_smooth()`) in the real data and the synthetic data generated by scDesign2 and the seven existing simulators (others). Note that ZINB-WaVE and SymSim filter out certain genes when simulating new data; Pearson correlation and Kendall's tau are only calculated between the genes whose zero proportions are less than 50%; gene-wise mean and variance and cell-wise library size are transformed to the $\log_{10}(1+x)$ scale (where $x$ represents a statistic's value).

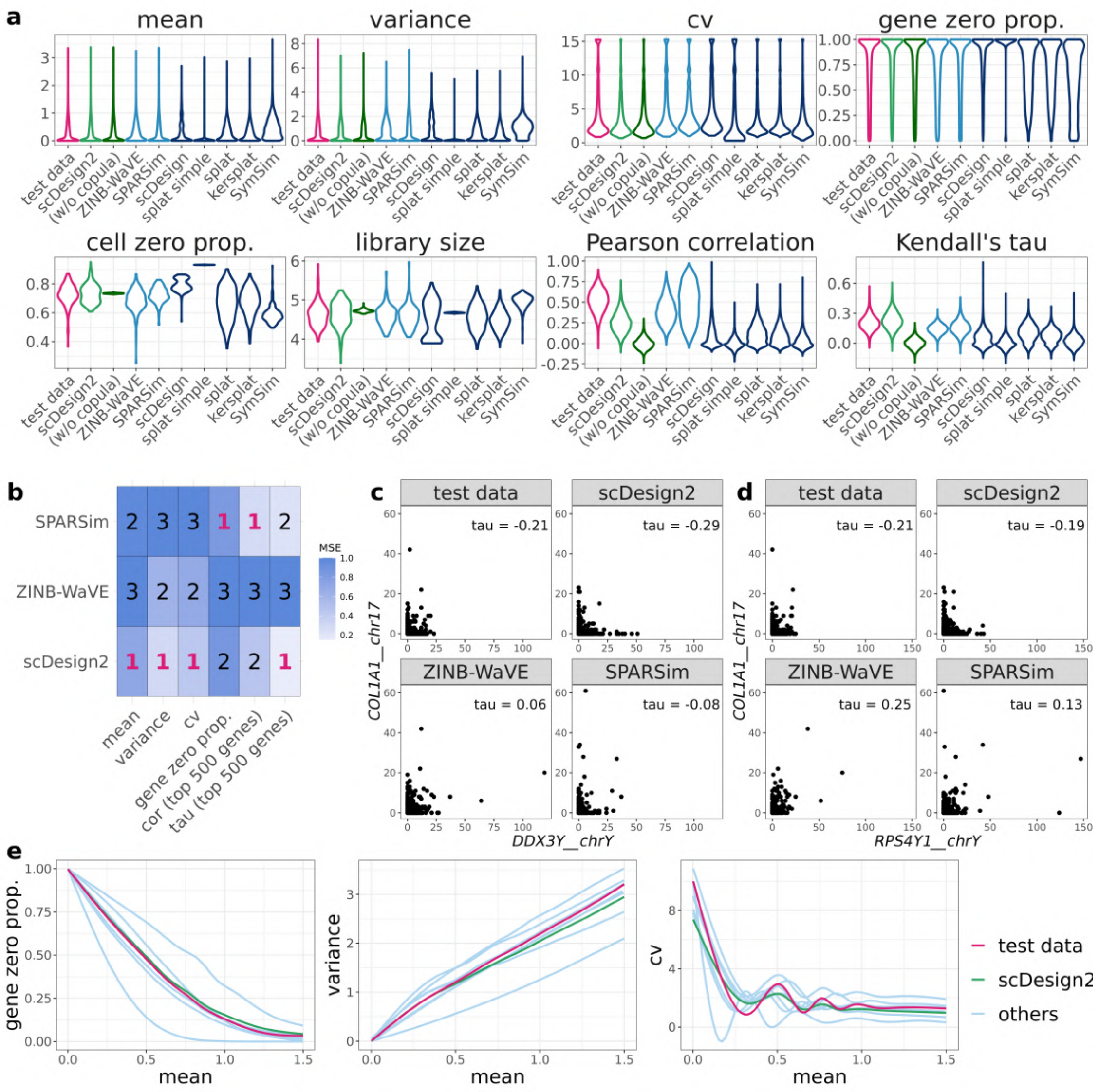


**Figure S10: Benchmarking scDesign2 against its variant without copula and seven existing scRNA-seq simulators for generating beta cells measured by CEL-Seq2.** (a) Distributions of eight summary statistics (gene-wise expression mean, variance, coefficient of variation (cv), and zero proportion; cell-wise zero proportion and library size; gene-pair-wise Pearson correlation and Kendall's tau) are plotted based on the real data (test data unused for training simulators) and the synthetic data generated by scDesign2, scDesign2 without copula (w/o copula), ZINB-WaVE, SPARSim, scDesign, three variants of the splatter package (splat simple, splat, and kersplat), and SymSim. (b) Ranking (with 1 being the best-performing method) of scDesign2, ZINB-WaVE, and SPARSim, the only three methods that preserve genes, in terms of the mean-squared error (MSE) of each of six summary statistics (four gene-wise and two gene-pair-wise) between the statistic values in the real data and the synthetic data generated by each simulator. Note that the color scale shows the normalized MSE: for each statistic (column in the table), the normalized MSEs are the MSEs divided by the largest MSE of that statistic. scDesign is ranked the top for four out of the six statistics. For the two gene-pair-wise statistics, we focus on the top 500 highly expressed genes, because as analyzed in the text, they are more meaningful, both biologically and statistically, than the correlations of the lowly expressed genes. (c)-(d) Scatterplots of two example gene pairs—*DDX3Y* vs. *COL1A1* and *RPS4Y1* vs. *COL1A1*—based on the real data and the synthetic data generated by scDesign2, ZINB-WaVE, and SPARSim. (e) Smoothed relationships between three pairs of gene-wise statistics (zero proportion vs. mean, variance vs. mean, and cv vs. mean) across all genes (curves plotted by the R function `geom_smooth()`) in the real data and the synthetic data generated by scDesign2 and the seven existing simulators (others). Note that ZINB-WaVE and SymSim filter out certain genes when simulating new data; Pearson correlation and Kendall's tau are only calculated between the genes whose zero proportions are less than 50%; gene-wise mean and variance and cell-wise library size are transformed to the $\log_{10}(1+x)$ scale (where $x$ represents a statistic's value).

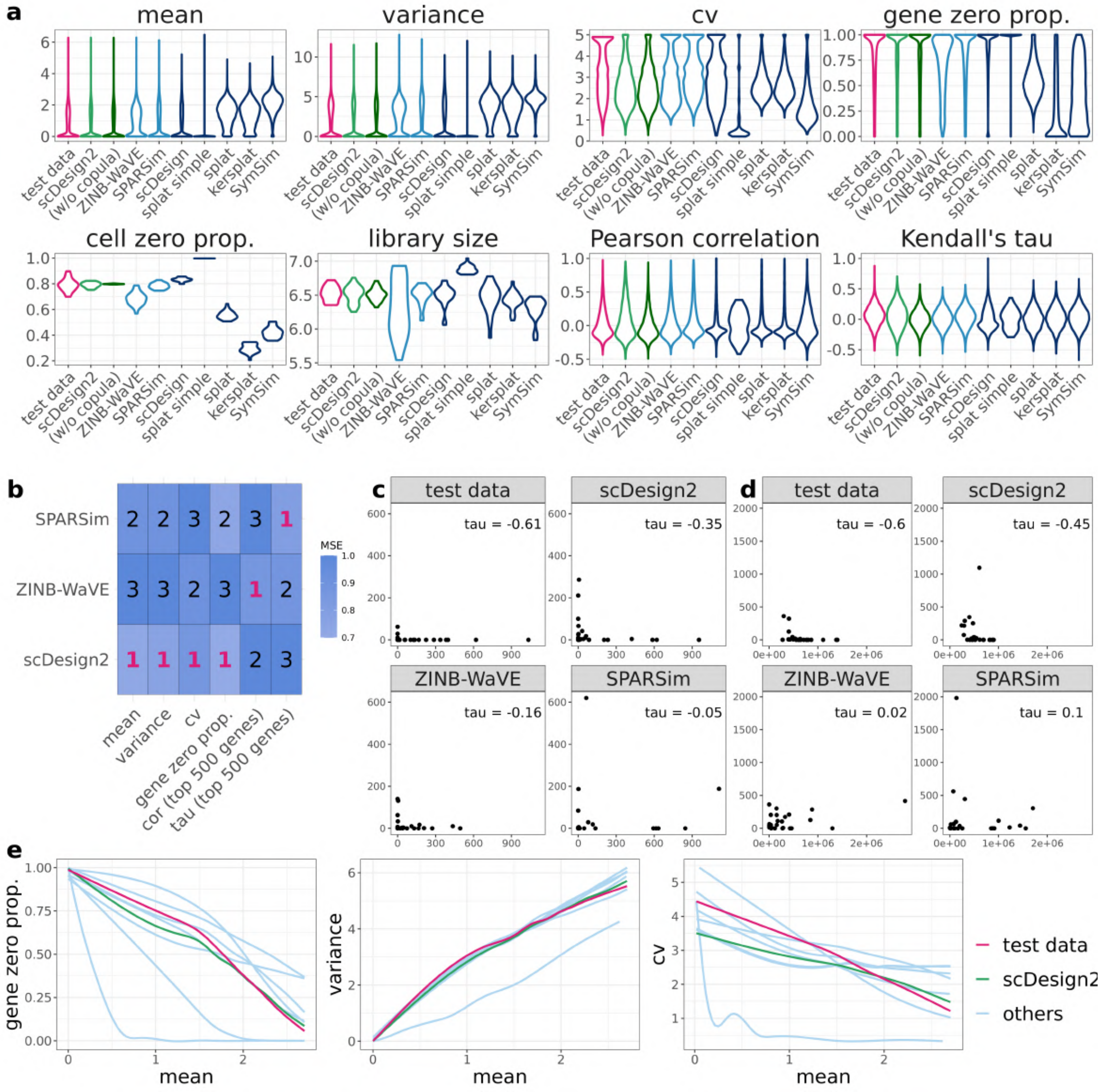


**Figure S11: Benchmarking scDesign2 against its variant without copula and seven existing scRNA-seq simulators for generating astrocytes measured by Fluidigm C1 (SMARTer).** (a) Distributions of eight summary statistics (gene-wise expression mean, variance, coefficient of variation (cv), and zero proportion; cell-wise zero proportion and library size; gene-pair-wise Pearson correlation and Kendall's tau) are plotted based on the real data (test data unused for training simulators) and the synthetic data generated by scDesign2, scDesign2 without copula (w/o copula), ZINB-WaVE, SPARSim, scDesign, three variants of the splatter package (splat simple, splat, and kersplat), and SymSim. (b) Ranking (with 1 being the best-performing method) of scDesign2, ZINB-WaVE, and SPARSim, the only three methods that preserve genes, in terms of the mean-squared error (MSE) of each of six summary statistics (four gene-wise and two gene-pair-wise) between the statistic values in the real data and the synthetic data generated by each simulator. Note that the color scale shows the normalized MSE: for each statistic (column in the table), the normalized MSEs are the MSEs divided by the largest MSE of that statistic. scDesign is ranked the top for four out of the six statistics. For the two gene-pair-wise statistics, we focus on the top 500 highly expressed genes, because as analyzed in the text, they are more meaningful, both biologically and statistically, than the correlations of the lowly expressed genes. (c)-(d) Scatterplots of two example gene pairs based on the real data and the synthetic data generated by scDesign2, ZINB-WaVE, and SPARSim. (e) Smoothed relationships between three pairs of gene-wise statistics (zero proportion vs. mean, variance vs. mean, and cv vs. mean) across all genes (curves plotted by the R function `geom_smooth()`) in the real data and the synthetic data generated by scDesign2 and the seven existing simulators (others). Note that ZINB-WaVE and SymSim filter out certain genes when simulating new data; Pearson correlation and Kendall's tau are only calculated between the genes whose zero proportions are less than 50%; gene-wise mean and variance and cell-wise library size are transformed to the $\log_{10}(1+x)$ scale (where $x$ represents a statistic's value).

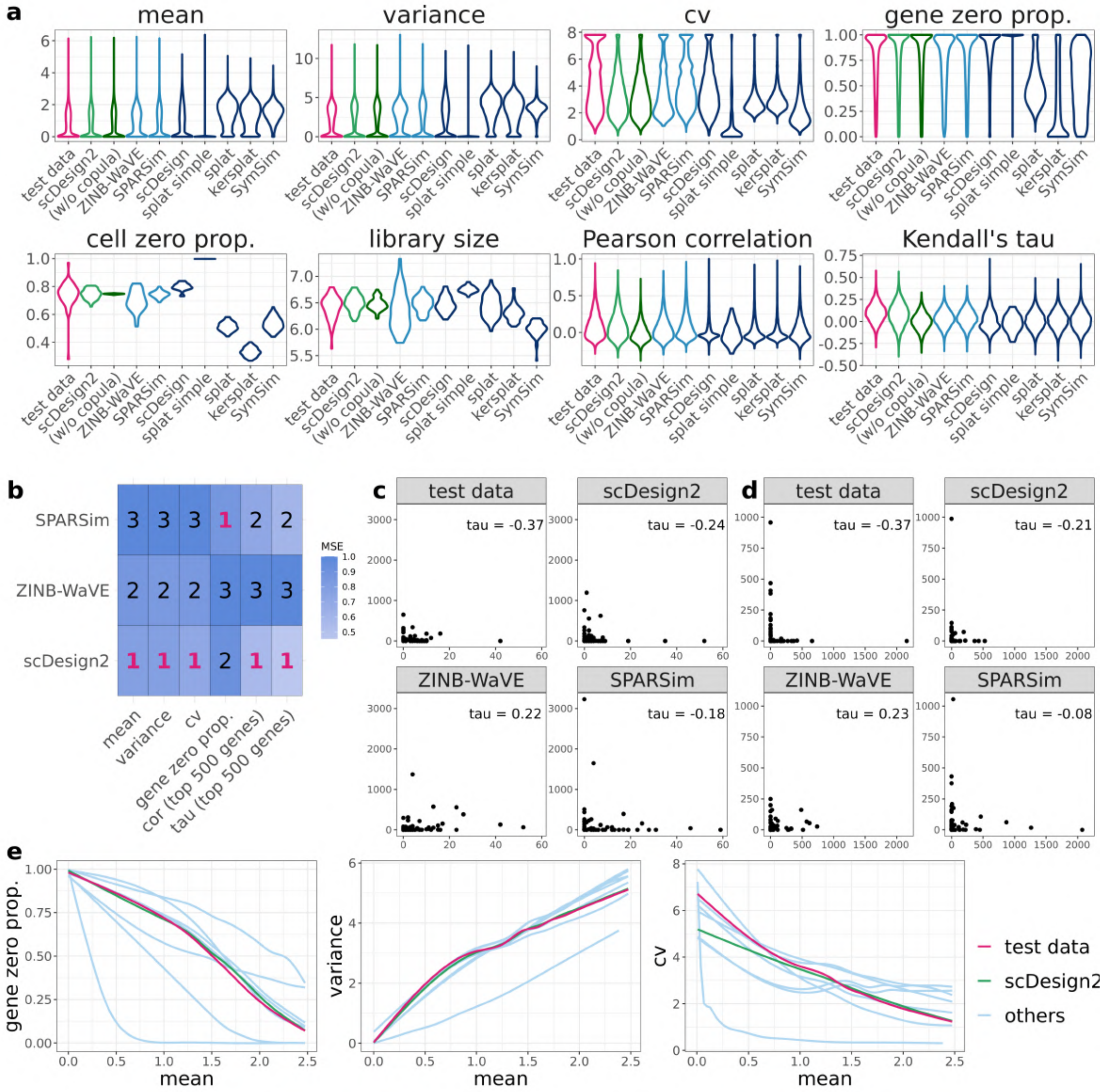


**Figure S12: Benchmarking scDesign2 against its variant without copula and seven existing scRNA-seq simulators for generating neurons measured by Fluidigm C1 (SMARTer).** (a) Distributions of eight summary statistics (gene-wise expression mean, variance, coefficient of variation (cv), and zero proportion; cell-wise zero proportion and library size; gene-pair-wise Pearson correlation and Kendall's tau) are plotted based on the real data (test data unused for training simulators) and the synthetic data generated by scDesign2, scDesign2 without copula (w/o copula), ZINB-WaVE, SPARSim, scDesign, three variants of the splatter package (splat simple, splat, and kersplat), and SymSim. (b) Ranking (with 1 being the best-performing method) of scDesign2, ZINB-WaVE, and SPARSim, the only three methods that preserve genes, in terms of the mean-squared error (MSE) of each of six summary statistics (four gene-wise and two gene-pair-wise) between the statistic values in the real data and the synthetic data generated by each simulator. Note that the color scale shows the normalized MSE: for each statistic (column in the table), the normalized MSEs are the MSEs divided by the largest MSE of that statistic. scDesign is ranked the top for five out of the six statistics. For the two gene-pair-wise statistics, we focus on the top 500 highly expressed genes, because as analyzed in the text, they are more meaningful, both biologically and statistically, than the correlations of the lowly expressed genes. (c)-(d) Scatterplots of two example gene pairs based on the real data and the synthetic data generated by scDesign2, ZINB-WaVE, and SPARSim. (e) Smoothed relationships between three pairs of gene-wise statistics (zero proportion vs. mean, variance vs. mean, and cv vs. mean) across all genes (curves plotted by the R function `geom_smooth()`) in the real data and the synthetic data generated by scDesign2 and the seven existing simulators (others). Note that ZINB-WaVE and SymSim filter out certain genes when simulating new data; Pearson correlation and Kendall's tau are only calculated between the genes whose zero proportions are less than 50%; gene-wise mean and variance and cell-wise library size are transformed to the $\log_{10}(1+x)$ scale (where $x$ represents a statistic's value).

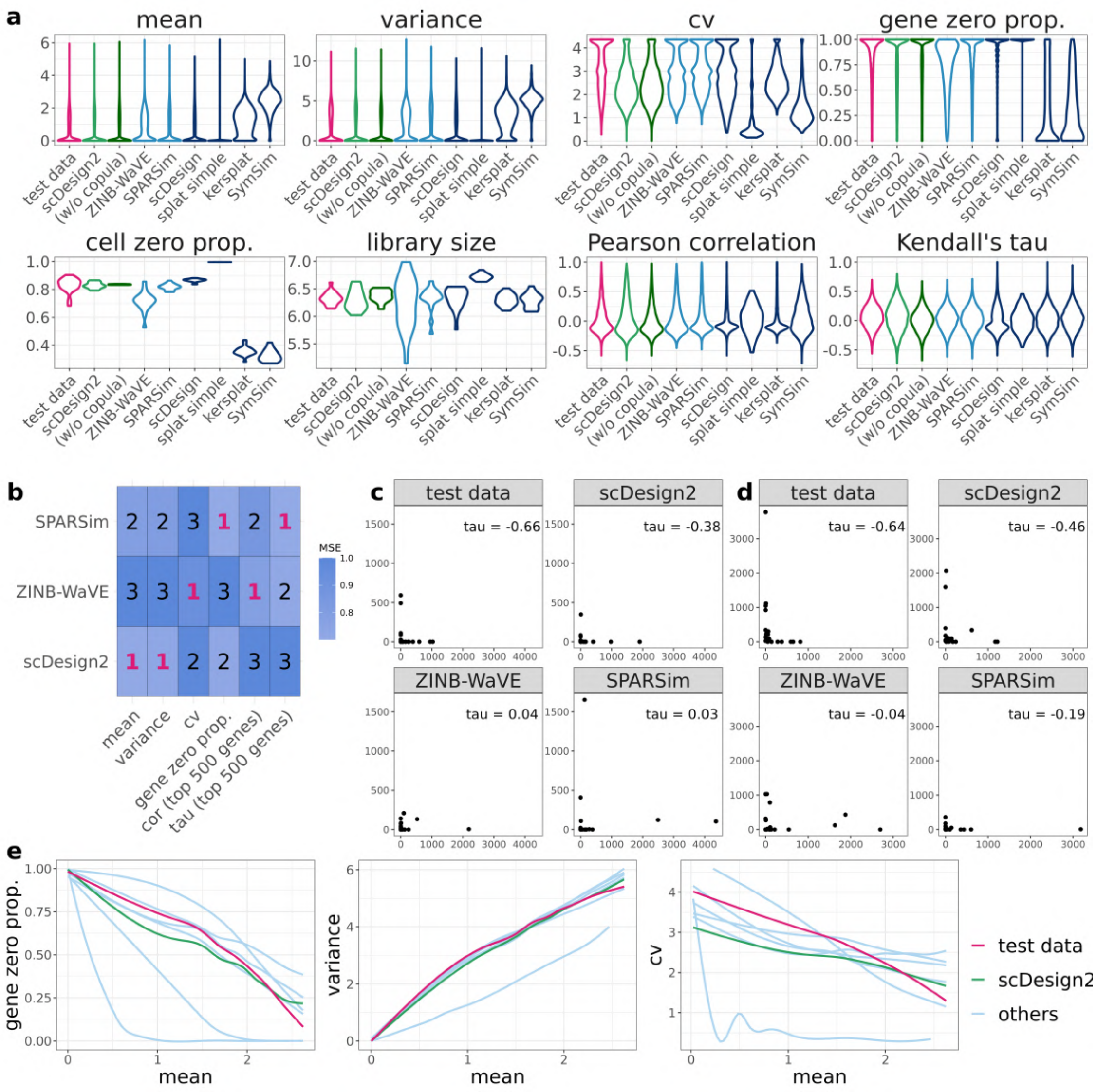


**Figure S13: Benchmarking scDesign2 against its variant without copula and seven existing scRNA-seq simulators for generating oligodendrocytes measured by Fluidigm C1 (SMARTer).** The input data is a oligodendrocytes dataset. (a) Distributions of eight summary statistics (gene-wise expression mean, variance, coefficient of variation (cv), and zero proportion; cell-wise zero proportion and library size; gene-pair-wise Pearson correlation and Kendall's tau) are plotted based on the real data (test data unused for training simulators) and the synthetic data generated by scDesign2, scDesign2 without copula (w/o copula), ZINB-WaVE, SPARSim, scDesign, three variants of the splatter package (splat simple, splat, and kersplat), and SymSim. (b) Ranking (with 1 being the best-performing method) of scDesign2, ZINB-WaVE, and SPARSim, the only three methods that preserve genes, in terms of the mean-squared error (MSE) of each of six summary statistics (four gene-wise and two gene-pair-wise) between the statistic values in the real data and the synthetic data generated by each simulator. Note that the color scale shows the normalized MSE: for each statistic (column in the table), the normalized MSEs are the MSEs divided by the largest MSE of that statistic. scDesign is ranked the top for two out of the six statistics. For the two gene-pair-wise statistics, we focus on the top 500 highly expressed genes, because as analyzed in the text, they are more meaningful, both biologically and statistically, than the correlations of the lowly expressed genes. (c)-(d) Scatterplots of two example gene pairs based on the real data and the synthetic data generated by scDesign2, ZINB-WaVE, and SPARSim. (e) Smoothed relationships between three pairs of gene-wise statistics (zero proportion vs. mean, variance vs. mean, and cv vs. mean) across all genes (curves plotted by the R function `geom_smooth()`) in the real data and the synthetic data generated by scDesign2 and the seven existing simulators (others). Note that ZINB-WaVE and SymSim filter out certain genes when simulating new data; Pearson correlation and Kendall's tau are only calculated between the genes whose zero proportions are less than 50%; gene-wise mean and variance and cell-wise library size are transformed to the $\log_{10}(1+x)$ scale (where $x$ represents a statistic's value).

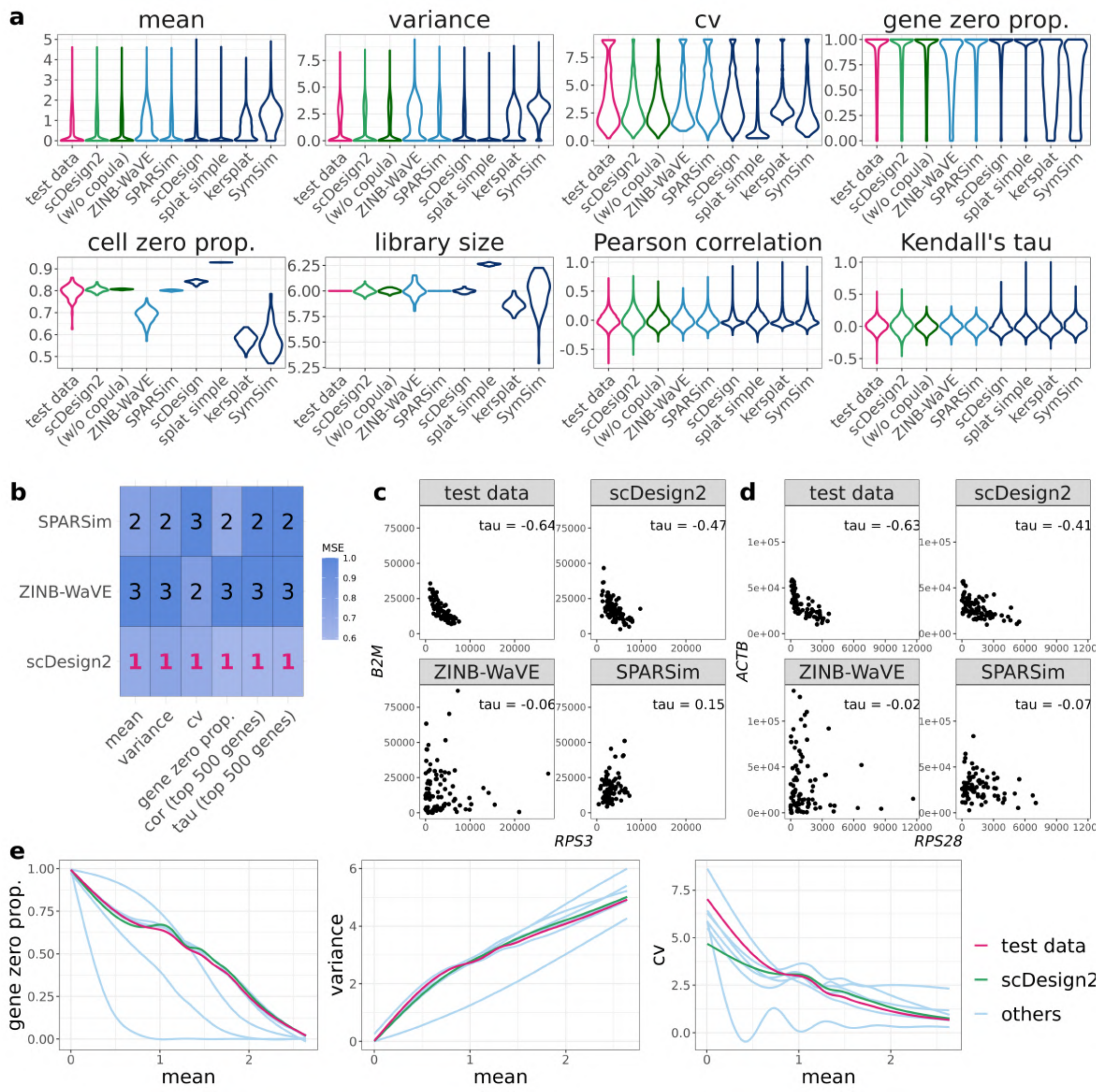


**Figure S14: Benchmarking scDesign2 against its variant without copula and seven existing scRNA-seq simulators for generating dendrocytes (subtype 1) measured by Smart-Seq2.** (a) Distributions of eight summary statistics (gene-wise expression mean, variance, coefficient of variation (cv), and zero proportion; cell-wise zero proportion and library size; gene-pair-wise Pearson correlation and Kendall's tau) are plotted based on the real data (test data unused for training simulators) and the synthetic data generated by scDesign2, scDesign2 without copula (w/o copula), ZINB-WaVE, SPARSim, scDesign, three variants of the splatter package (splat simple, splat, and kersplat), and SymSim. (b) Ranking (with 1 being the best-performing method) of scDesign2, ZINB-WaVE, and SPARSim, the only three methods that preserve genes, in terms of the mean-squared error (MSE) of each of six summary statistics (four gene-wise and two gene-pair-wise) between the statistic values in the real data and the synthetic data generated by each simulator. Note that the color scale shows the normalized MSE: for each statistic (column in the table), the normalized MSEs are the MSEs divided by the largest MSE of that statistic. scDesign is ranked the top for six out of the six statistics. For the two gene-pair-wise statistics, we focus on the top 500 highly expressed genes, because as analyzed in the text, they are more meaningful, both biologically and statistically, than the correlations of the lowly expressed genes. (c)-(d) Scatterplots of two example gene pairs—*RPS3* vs. *B2M* and *RPS28* vs. *ACTB*—based on the real data and the synthetic data generated by scDesign2, ZINB-WaVE, and SPARSim. (e) Smoothed relationships between three pairs of gene-wise statistics (zero proportion vs. mean, variance vs. mean, and cv vs. mean) across all genes (curves plotted by the R function `geom_smooth()`) in the real data and the synthetic data generated by scDesign2 and the seven existing simulators (others). Note that ZINB-WaVE and SymSim filter out certain genes when simulating new data; Pearson correlation and Kendall's tau are only calculated between the genes whose zero proportions are less than 50%; gene-wise mean and variance and cell-wise library size are transformed to the $\log_{10}(1+x)$ scale (where $x$ represents a statistic's value).

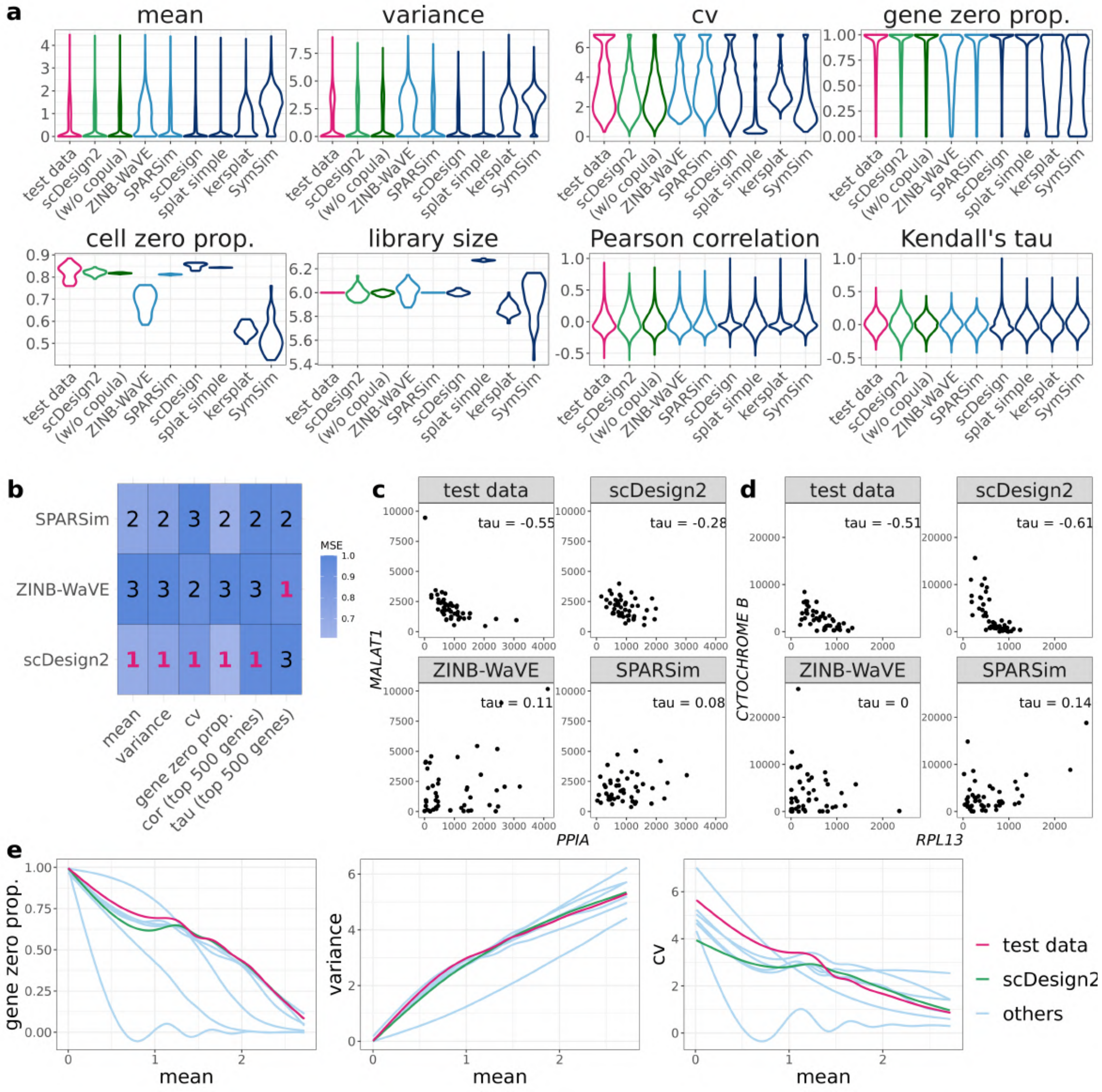


**Figure S15: Benchmarking scDesign2 against its variant without copula and seven existing scRNA-seq simulators for generating dendrocytes (subtype 2) measured by Smart-Seq2.** (a) Distributions of eight summary statistics (gene-wise expression mean, variance, coefficient of variation (cv), and zero proportion; cell-wise zero proportion and library size; gene-pair-wise Pearson correlation and Kendall's tau) are plotted based on the real data (test data unused for training simulators) and the synthetic data generated by scDesign2, scDesign2 without copula (w/o copula), ZINB-WaVE, SPARSim, scDesign, three variants of the splatter package (splat simple, splat, and kersplat), and SymSim. (b) Ranking (with 1 being the best-performing method) of scDesign2, ZINB-WaVE, and SPARSim, the only three methods that preserve genes, in terms of the mean-squared error (MSE) of each of six summary statistics (four gene-wise and two gene-pair-wise) between the statistic values in the real data and the synthetic data generated by each simulator. Note that the color scale shows the normalized MSE: for each statistic (column in the table), the normalized MSEs are the MSEs divided by the largest MSE of that statistic. scDesign is ranked the top for five out of the six statistics. For the two gene-pair-wise statistics, we focus on the top 500 highly expressed genes, because as analyzed in the text, they are more meaningful, both biologically and statistically, than the correlations of the lowly expressed genes. (c)-(d) Scatterplots of two example gene pairs—*PPIA* vs. *MALAT1* and *RPL13* vs. *CYTOCHROME B*—based on the real data and the synthetic data generated by scDesign2, ZINB-WaVE, and SPARSim. Only scDesign2 captures the negative gene correlations in the real data. (e) Smoothed relationships between three pairs of gene-wise statistics (zero proportion vs. mean, variance vs. mean, and cv vs. mean) across all genes (curves plotted by the R function `geom_smooth()`) in the real data and the synthetic data generated by scDesign2 and the seven existing simulators (others). Note that ZINB-WaVE and SymSim filter out certain genes when simulating new data; Pearson correlation and Kendall's tau are only calculated between the genes whose zero proportions are less than 50%; gene-wise mean and variance and cell-wise library size are transformed to the $\log_{10}(1+x)$ scale (where $x$ represents a statistic's value).

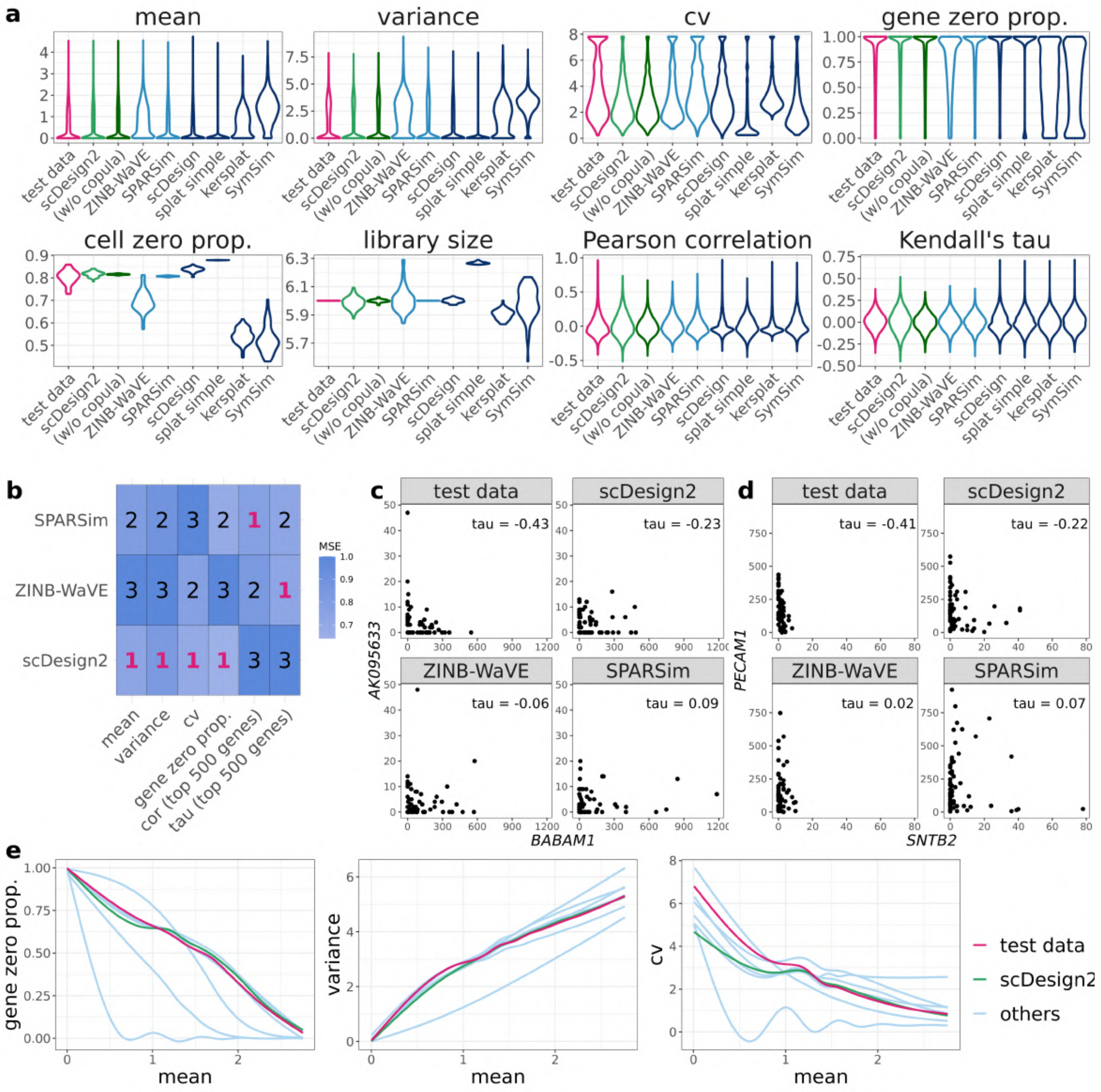


**Figure S16: Benchmarking scDesign2 against its variant without copula and seven existing scRNA-seq simulators for generating monocytes (subtype 2) measured by Smart-Seq2.** (a) Distributions of eight summary statistics (gene-wise expression mean, variance, coefficient of variation (cv), and zero proportion; cell-wise zero proportion and library size; gene-pair-wise Pearson correlation and Kendall's tau) are plotted based on the real data (test data unused for training simulators) and the synthetic data generated by scDesign2, scDesign2 without copula (w/o copula), ZINB-WaVE, SPARSim, scDesign, three variants of the splatter package (splat simple, splat, and kersplat), and SymSim. (b) Ranking (with 1 being the best-performing method) of scDesign2, ZINB-WaVE, and SPARSim, the only three methods that preserve genes, in terms of the mean-squared error (MSE) of each of six summary statistics (four gene-wise and two gene-pair-wise) between the statistic values in the real data and the synthetic data generated by each simulator. Note that the color scale shows the normalized MSE: for each statistic (column in the table), the normalized MSEs are the MSEs divided by the largest MSE of that statistic. scDesign is ranked the top for four out of the six statistics. For the two gene-pair-wise statistics, we focus on the top 500 highly expressed genes, because as analyzed in the text, they are more meaningful, both biologically and statistically, than the correlations of the lowly expressed genes. (c)-(d) Scatterplots of two example gene pairs—*BABAM1* vs. *AK095633* and *SNTB2* vs. *PECAM1*—based on the real data and the synthetic data generated by scDesign2, ZINB-WaVE, and SPARSim. (e) Smoothed relationships between three pairs of gene-wise statistics (zero proportion vs. mean, variance vs. mean, and cv vs. mean) across all genes (curves plotted by the R function `geom_smooth()`) in the real data and the synthetic data generated by scDesign2 and the seven existing simulators (others). Note that ZINB-WaVE and SymSim filter out certain genes when simulating new data; Pearson correlation and Kendall's tau are only calculated between the genes whose zero proportions are less than 50%; gene-wise mean and variance and cell-wise library size are transformed to the $\log_{10}(1+x)$ scale (where $x$ represents a statistic's value).

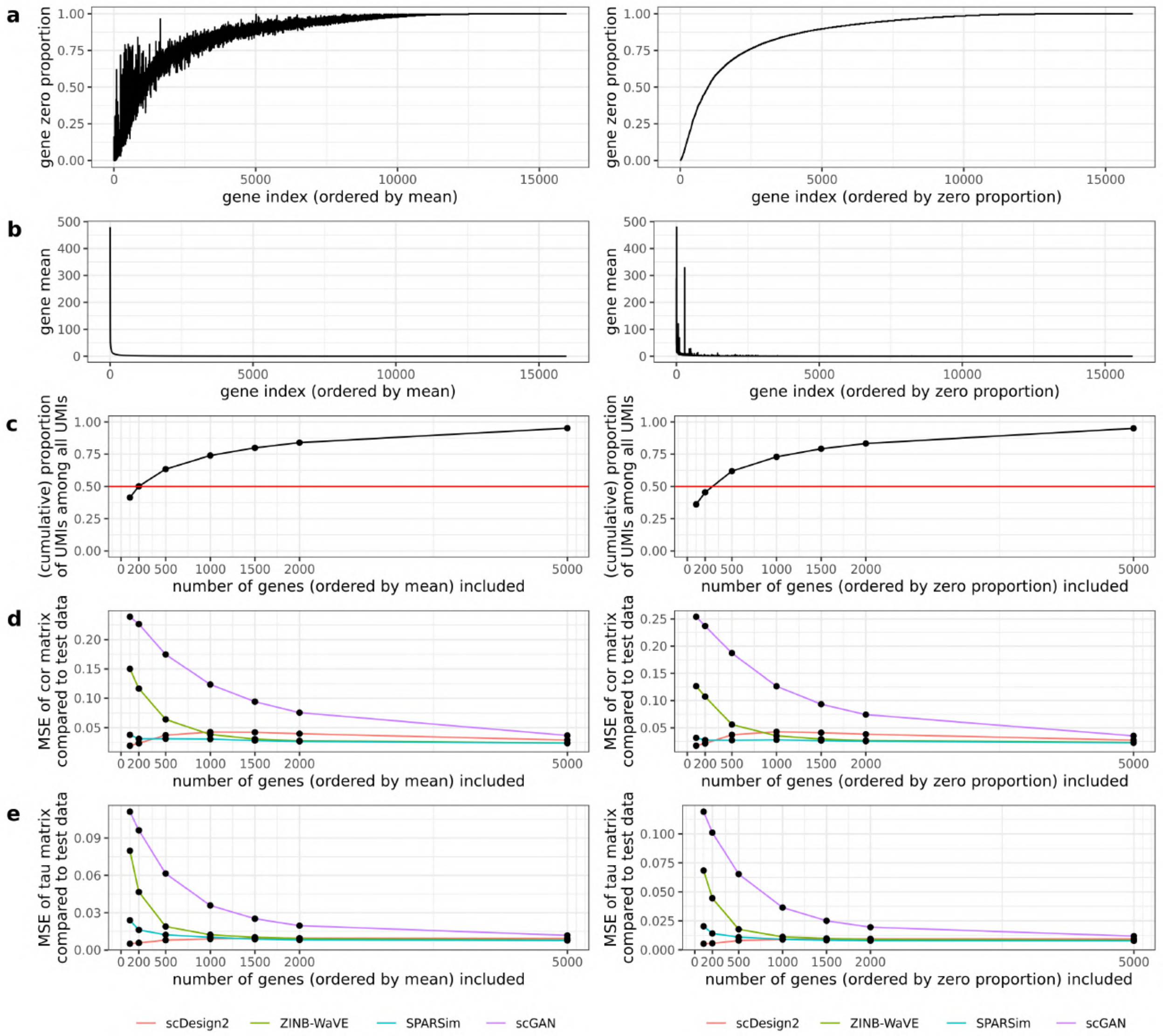


**Figure S17: Relationships of the mean squared error (MSE) vs. the dimension (i.e., number of genes) of the (Pearson or Kendall's tau) gene correlation matrices, which are estimated from the synthetic data generated by four simulators trained on the 10x Genomics goblet cell data.** The genes are ordered in two ways, by their mean expression from high to low (left) or by their zero proportion from low to high (right). The plots are shown for the top 100, 200, 500, 1000, 1500, and 2000 genes ordered in either way. (a) The zero proportion of each top gene. (b) The mean expression of each top gene. (c) The relationship of the cumulative proportion of the top genes' UMIs among all UMIs vs. the number of top genes. (d)–(e) The MSE is defined as the average per-entry squared difference between the correlation matrices estimated from each synthetic dataset and the test data. Each plot shows the relationship of the MSE of (d) the Pearson correlation matrix or (e) the Kendall's tau matrix for the top genes estimated from each simulator's synthetic data vs. the number of top genes.

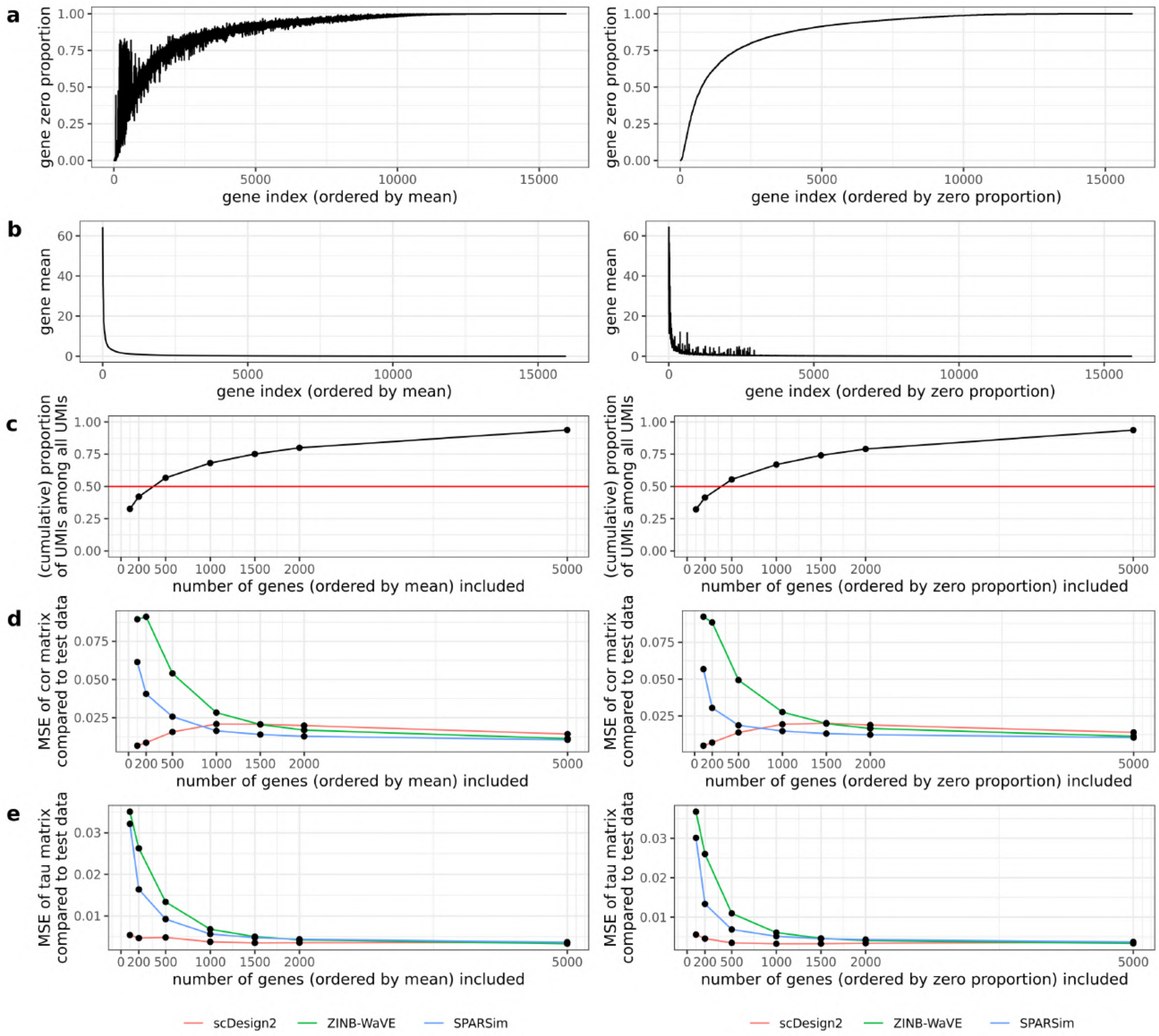


**Figure S18: Relationships of the mean squared error (MSE) vs. the dimension (i.e., number of genes) of the (Pearson or Kendall's tau) gene correlation matrices, which are estimated from the synthetic data generated by three simulators trained on the 10x Genomics stem cell data.** The genes are ordered in two ways, by their mean expression from high to low (left) or by their zero proportion from low to high (right). The plots are shown for the top 100, 200, 500, 1000, 1500, and 2000 genes ordered in either way. (a) The zero proportion of each top gene. (b) The mean expression of each top gene. (c) The relationship of the cumulative proportion of the top genes' UMIs among all UMIs vs. the number of top genes. (d)–(e) The MSE is defined as the average per-entry squared difference between the correlation matrices estimated from each synthetic dataset and the test data. Each plot shows the relationship of the MSE of (d) the Pearson correlation matrix or (e) the Kendall's tau matrix for the top genes estimated from each simulator's synthetic data vs. the number of top genes.

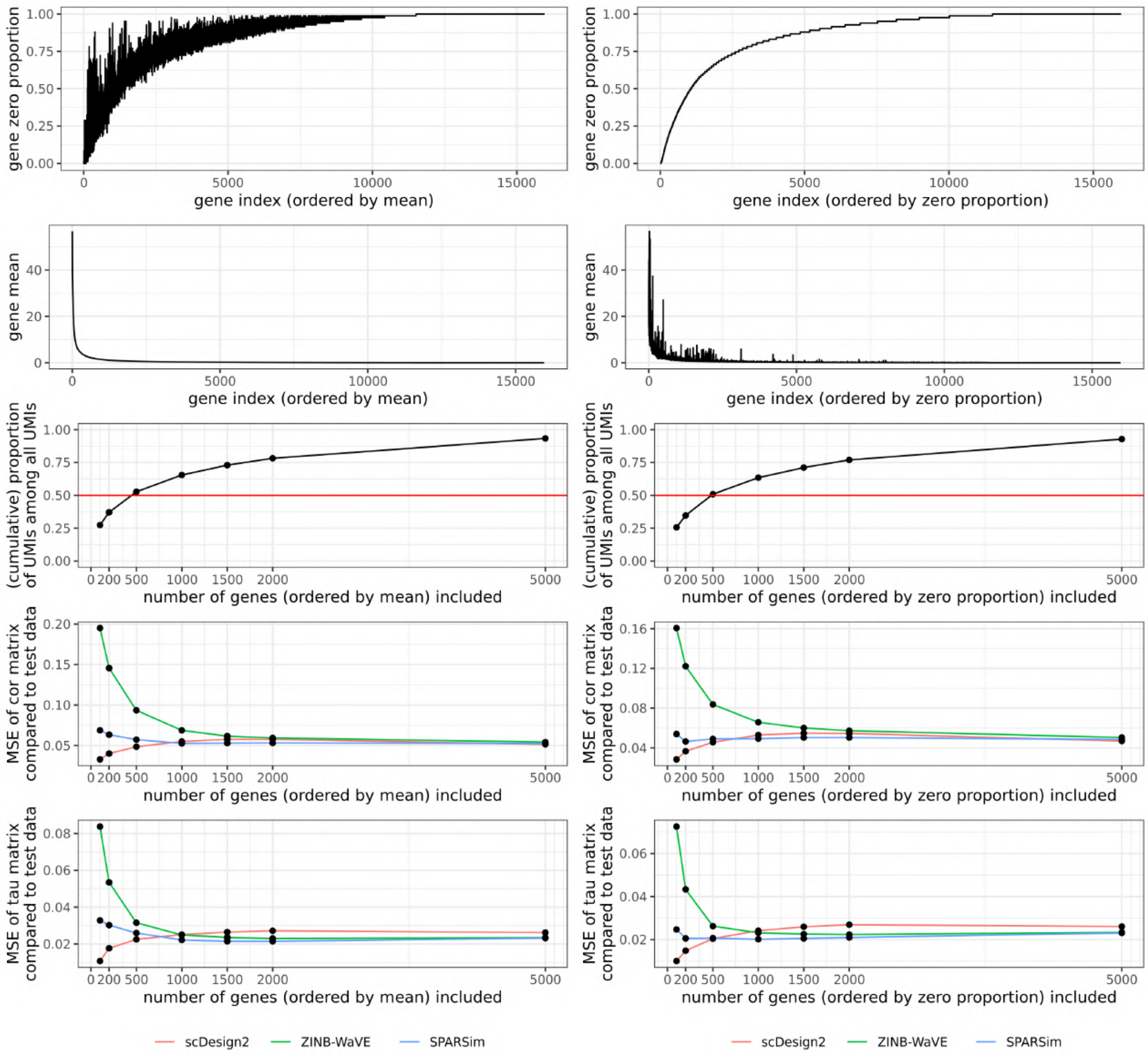


**Figure S19: Relationships of the mean squared error (MSE) vs. the dimension (i.e., number of genes) of the (Pearson or Kendall's tau) gene correlation matrices, which are estimated from the synthetic data generated by three simulators trained on the 10x Genomics tuft cell data.** The genes are ordered in two ways, by their mean expression from high to low (left) or by their zero proportion from low to high (right). The plots are shown for the top 100, 200, 500, 1000, 1500, and 2000 genes ordered in either way. (a) The zero proportion of each top gene. (b) The mean expression of each top gene. (c) The relationship of the cumulative proportion of the top genes' UMIs among all UMIs vs. the number of top genes. (d)–(e) The MSE is defined as the average per-entry squared difference between the correlation matrices estimated from each synthetic dataset and the test data. Each plot shows the relationship of the MSE of (d) the Pearson correlation matrix or (e) the Kendall's tau matrix for the top genes estimated from each simulator's synthetic data vs. the number of top genes.

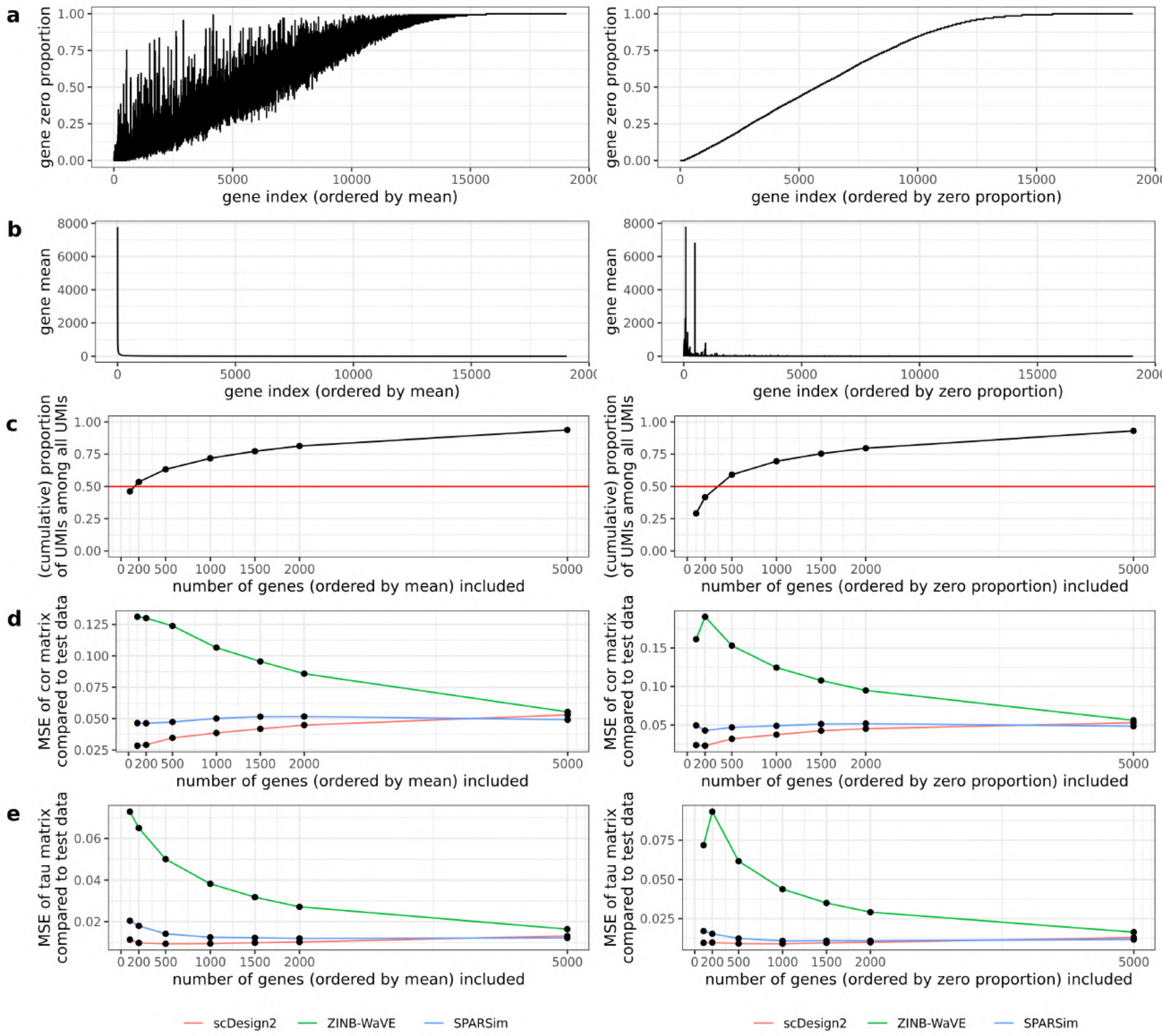


**Figure S20: Relationships of the mean squared error (MSE) vs. the dimension (i.e., number of genes) of the (Pearson or Kendall's tau) gene correlation matrices, which are estimated from the synthetic data generated by three simulators trained on the CEL-Seq2 acinar cell data.** The genes are ordered in two ways, by their mean expression from high to low (left) or by their zero proportion from low to high (right). The plots are shown for the top 100, 200, 500, 1000, 1500, and 2000 genes ordered in either way. (a) The zero proportion of each top gene. (b) The mean expression of each top gene. (c) The relationship of the cumulative proportion of the top genes' UMIs among all UMIs vs. the number of top genes. (d)–(e) The MSE is defined as the average per-entry squared difference between the correlation matrices estimated from each synthetic dataset and the test data. Each plot shows the relationship of the MSE of (d) the Pearson correlation matrix or (e) the Kendall's tau matrix for the top genes estimated from each simulator's synthetic data vs. the number of top genes.

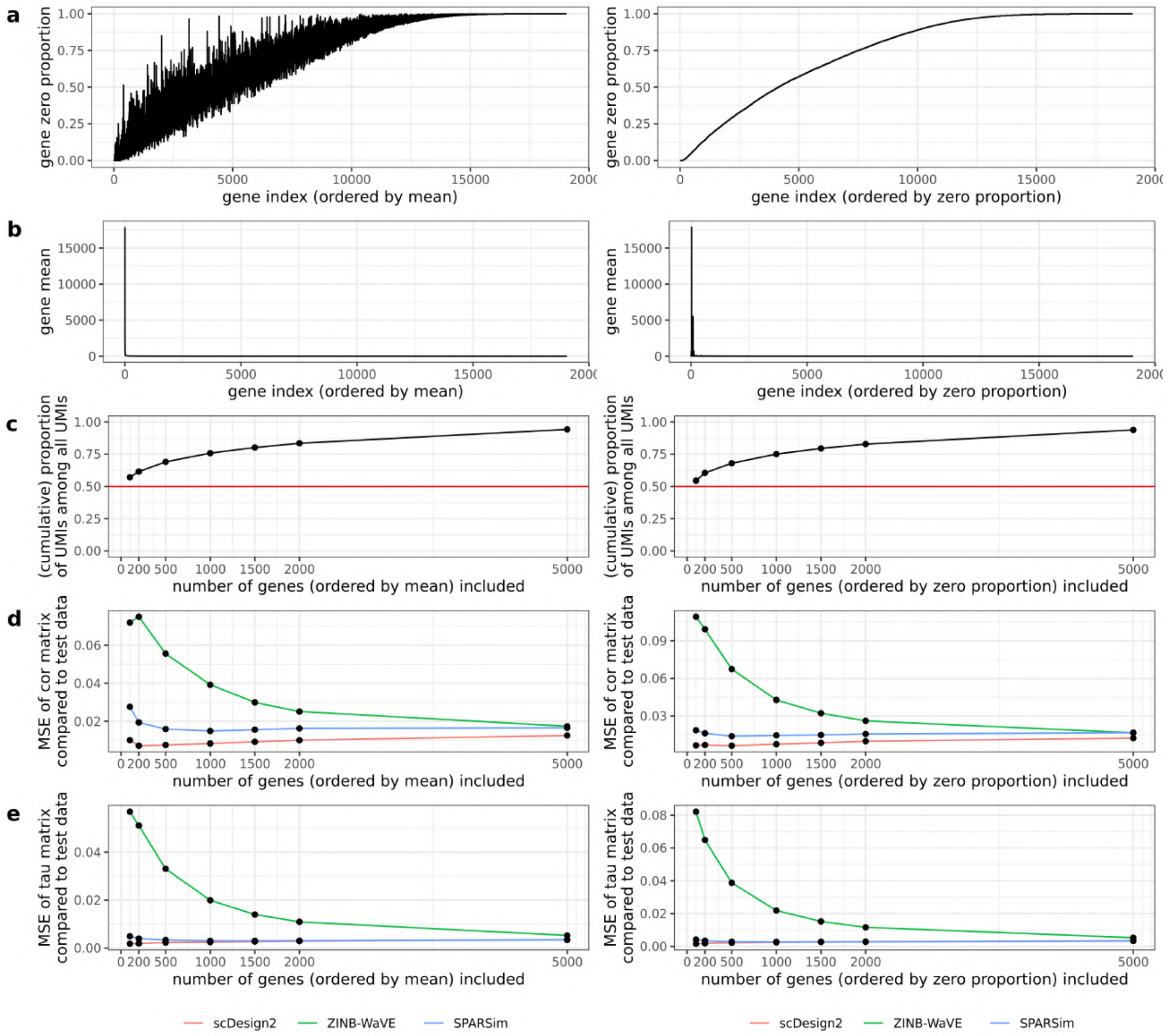


**Figure S21: Relationships of the mean squared error (MSE) vs. the dimension (i.e., number of genes) of the (Pearson or Kendall's tau) gene correlation matrices, which are estimated from the synthetic data generated by three simulators trained on the CEL-Seq2 alpha cell data.** The genes are ordered in two ways, by their mean expression from high to low (left) or by their zero proportion from low to high (right). The plots are shown for the top 100, 200, 500, 1000, 1500, and 2000 genes ordered in either way. (a) The zero proportion of each top gene. (b) The mean expression of each top gene. (c) The relationship of the cumulative proportion of the top genes' UMIs among all UMIs vs. the number of top genes. (d)–(e) The MSE is defined as the average per-entry squared difference between the correlation matrices estimated from each synthetic dataset and the test data. Each plot shows the relationship of the MSE of (d) the Pearson correlation matrix or (e) the Kendall's tau matrix for the top genes estimated from each simulator's synthetic data vs. the number of top genes.

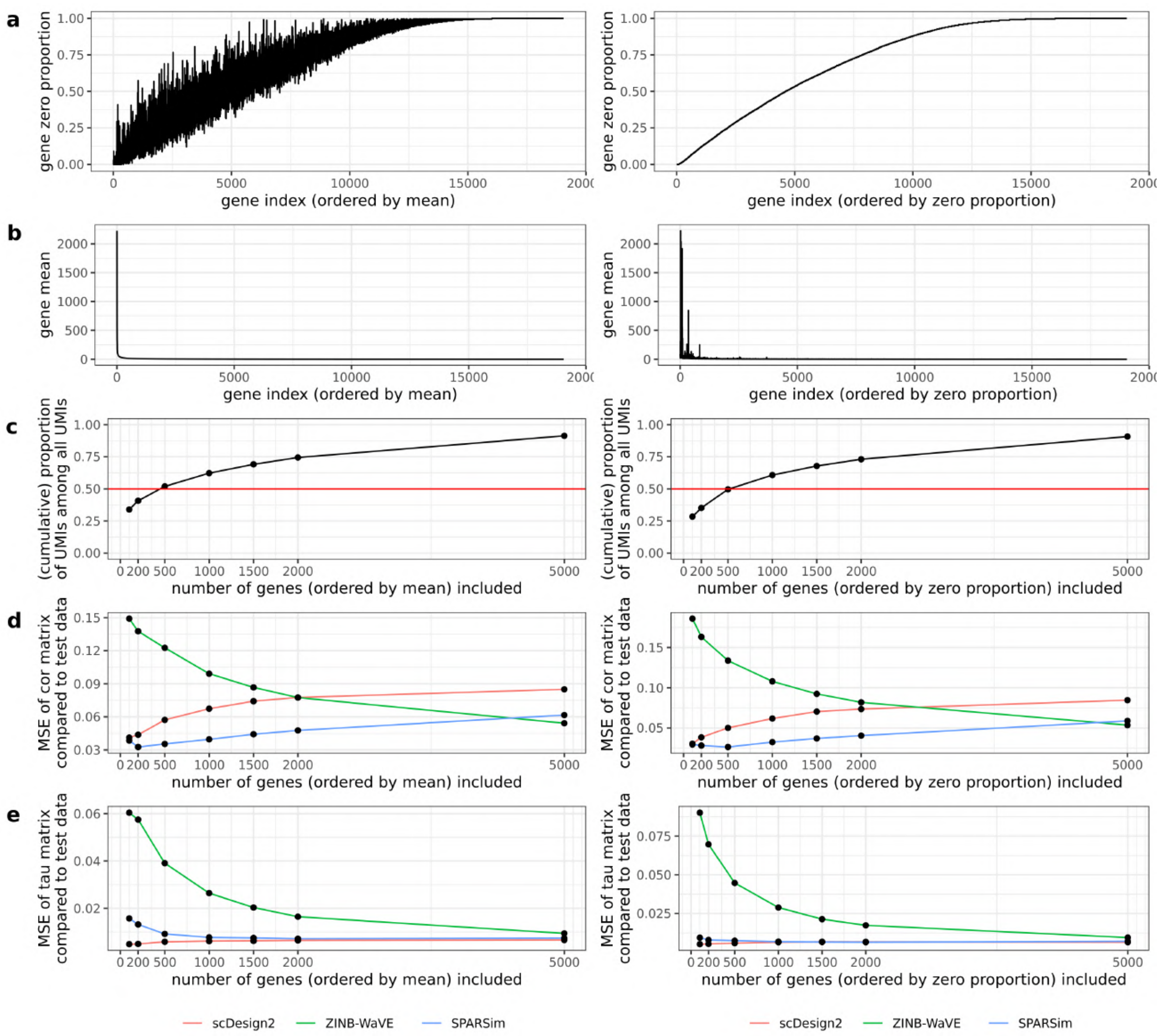


**Figure S22: Relationships of the mean squared error (MSE) vs. the dimension (i.e., number of genes) of the (Pearson or Kendall's tau) gene correlation matrices, which are estimated from the synthetic data generated by three simulators trained on the CEL-Seq2 beta cell data.** The genes are ordered in two ways, by their mean expression from high to low (left) or by their zero proportion from low to high (right). The plots are shown for the top 100, 200, 500, 1000, 1500, and 2000 genes ordered in either way. (a) The zero proportion of each top gene. (b) The mean expression of each top gene. (c) The relationship of the cumulative proportion of the top genes' UMIs among all UMIs vs. the number of top genes. (d)–(e) The MSE is defined as the average per-entry squared difference between the correlation matrices estimated from each synthetic dataset and the test data. Each plot shows the relationship of the MSE of (d) the Pearson correlation matrix or (e) the Kendall's tau matrix for the top genes estimated from each simulator's synthetic data vs. the number of top genes.

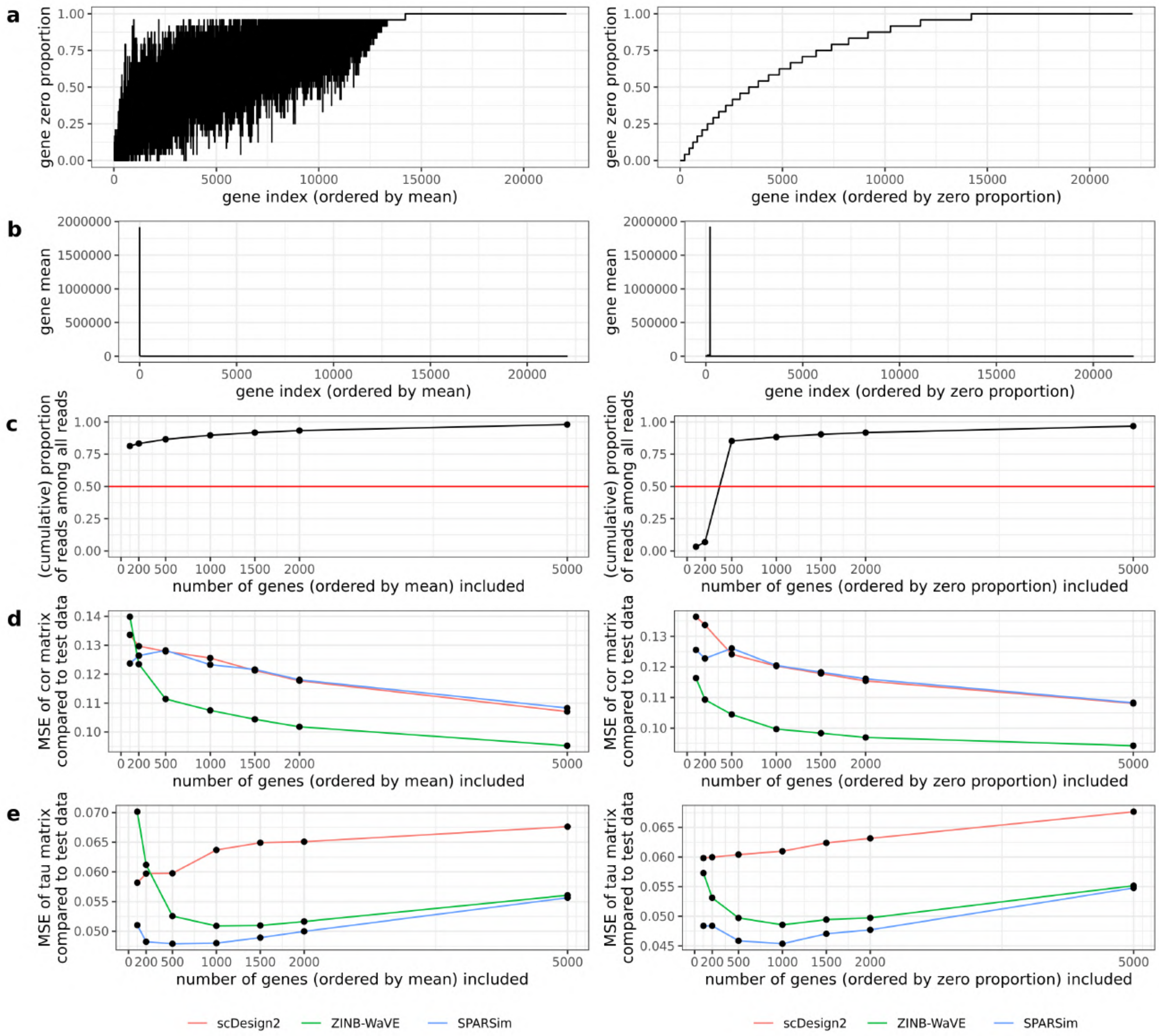


**Figure S23: Relationships of the mean squared error (MSE) vs. the dimension (i.e., number of genes) of the (Pearson or Kendall's tau) gene correlation matrices, which are estimated from the synthetic data generated by three simulators trained on the Fluidigm C1 astrocytes data.** The genes are ordered in two ways, by their mean expression from high to low (left) or by their zero proportion from low to high (right). The plots are shown for the top 100, 200, 500, 1000, 1500, and 2000 genes ordered in either way. (a) The zero proportion of each top gene. (b) The mean expression of each top gene. (c) The relationship of the cumulative proportion of the top genes' UMIs among all UMIs vs. the number of top genes. (d)–(e) The MSE is defined as the average per-entry squared difference between the correlation matrices estimated from each synthetic dataset and the test data. Each plot shows the relationship of the MSE of (d) the Pearson correlation matrix or (e) the Kendall's tau matrix for the top genes estimated from each simulator's synthetic data vs. the number of top genes.

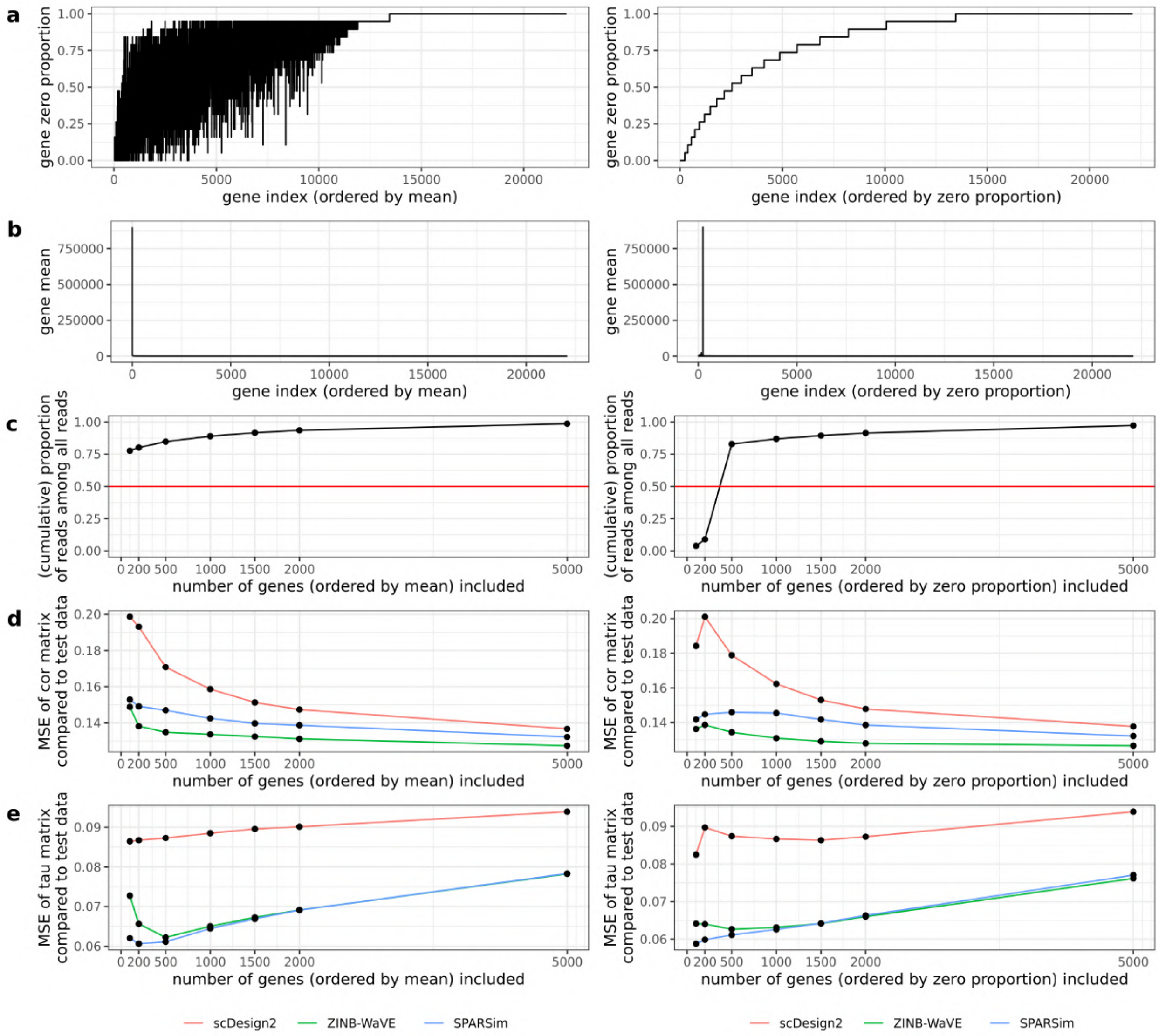


**Figure S24: Relationships of the mean squared error (MSE) vs. the dimension (i.e., number of genes) of the (Pearson or Kendall's tau) gene correlation matrices, which are estimated from the synthetic data generated by three simulators trained on the Fluidigm C1 oligodendrocytes data.** The genes are ordered in two ways, by their mean expression from high to low (left) or by their zero proportion from low to high (right). The plots are shown for the top 100, 200, 500, 1000, 1500, and 2000 genes ordered in either way. (a) The zero proportion of each top gene. (b) The mean expression of each top gene. (c) The relationship of the cumulative proportion of the top genes' UMIs among all UMIs vs. the number of top genes. (d)–(e) The MSE is defined as the average per-entry squared difference between the correlation matrices estimated from each synthetic dataset and the test data. Each plot shows the relationship of the MSE of (d) the Pearson correlation matrix or (e) the Kendall's tau matrix for the top genes estimated from each simulator's synthetic data vs. the number of top genes.

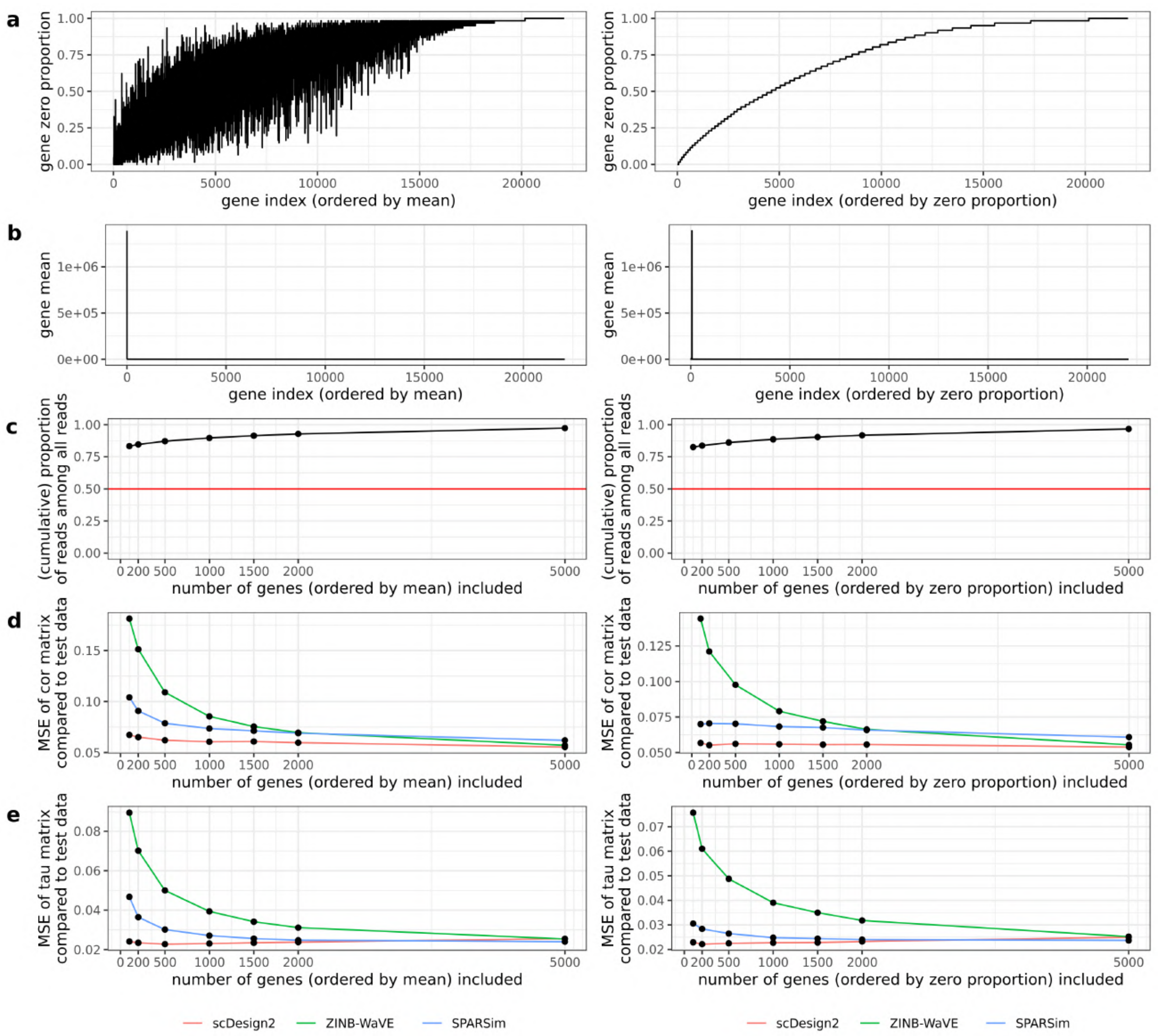


**Figure S25: Relationships of the mean squared error (MSE) vs. the dimension (i.e., number of genes) of the (Pearson or Kendall's tau) gene correlation matrices, which are estimated from the synthetic data generated by three simulators trained on the Fluidigm C1 neurons data.** The genes are ordered in two ways, by their mean expression from high to low (left) or by their zero proportion from low to high (right). The plots are shown for the top 100, 200, 500, 1000, 1500, and 2000 genes ordered in either way. (a) The zero proportion of each top gene. (b) The mean expression of each top gene. (c) The relationship of the cumulative proportion of the top genes' UMIs among all UMIs vs. the number of top genes. (d)–(e) The MSE is defined as the average per-entry squared difference between the correlation matrices estimated from each synthetic dataset and the test data. Each plot shows the relationship of the MSE of (d) the Pearson correlation matrix or (e) the Kendall's tau matrix for the top genes estimated from each simulator's synthetic data vs. the number of top genes.

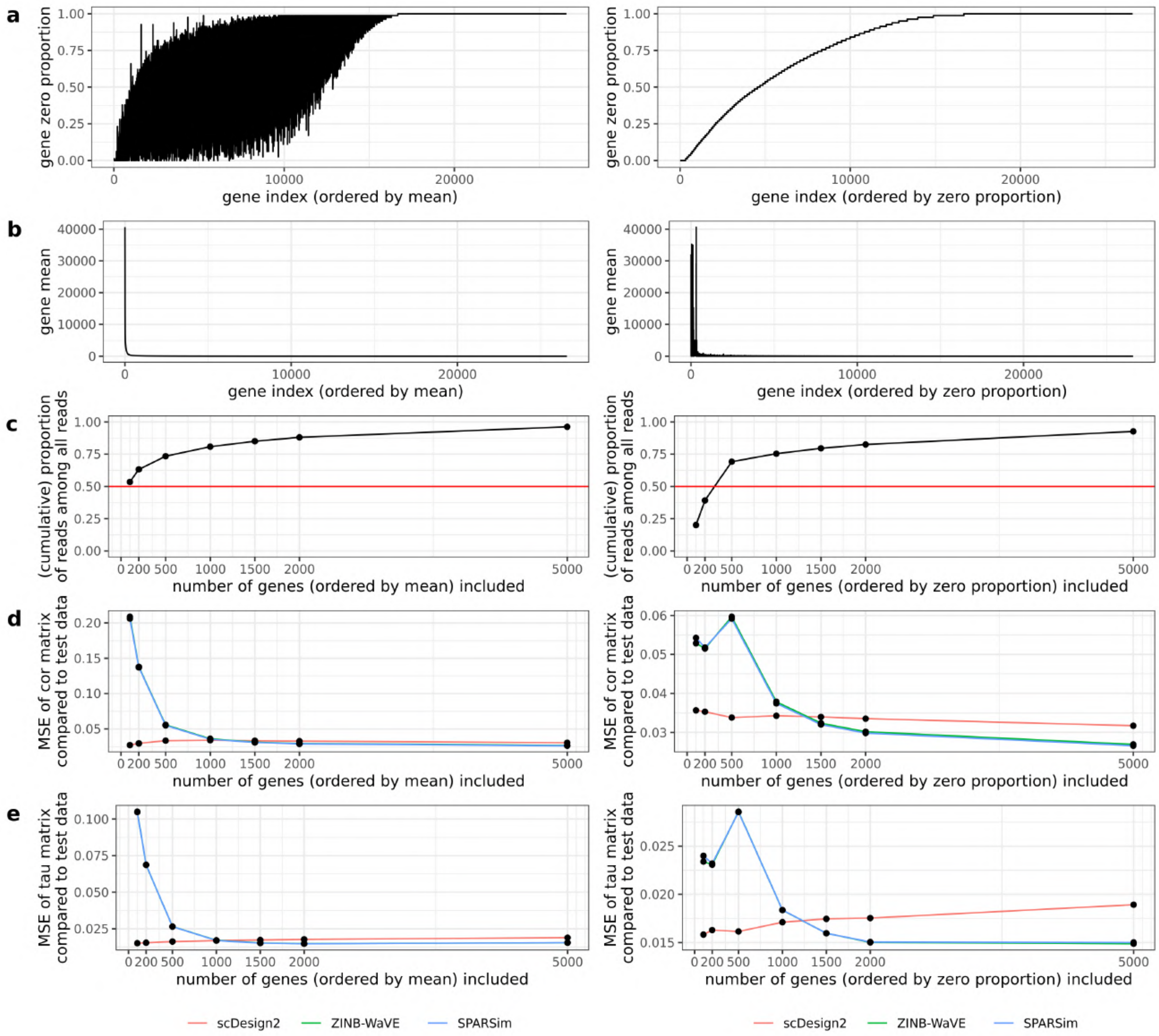


**Figure S26: Relationships of the mean squared error (MSE) vs. the dimension (i.e., number of genes) of the (Pearson or Kendall's tau) gene correlation matrices, which are estimated from the synthetic data generated by three simulators trained on the Smart-Seq2 dendrocyte (subtype 1) data.** The genes are ordered in two ways, by their mean expression from high to low (left) or by their zero proportion from low to high (right). The plots are shown for the top 100, 200, 500, 1000, 1500, and 2000 genes ordered in either way. (a) The zero proportion of each top gene. (b) The mean expression of each top gene. (c) The relationship of the cumulative proportion of the top genes' UMIs among all UMIs vs. the number of top genes. (d)–(e) The MSE is defined as the average per-entry squared difference between the correlation matrices estimated from each synthetic dataset and the test data. Each plot shows the relationship of the MSE of (d) the Pearson correlation matrix or (e) the Kendall's tau matrix for the top genes estimated from each simulator's synthetic data vs. the number of top genes.

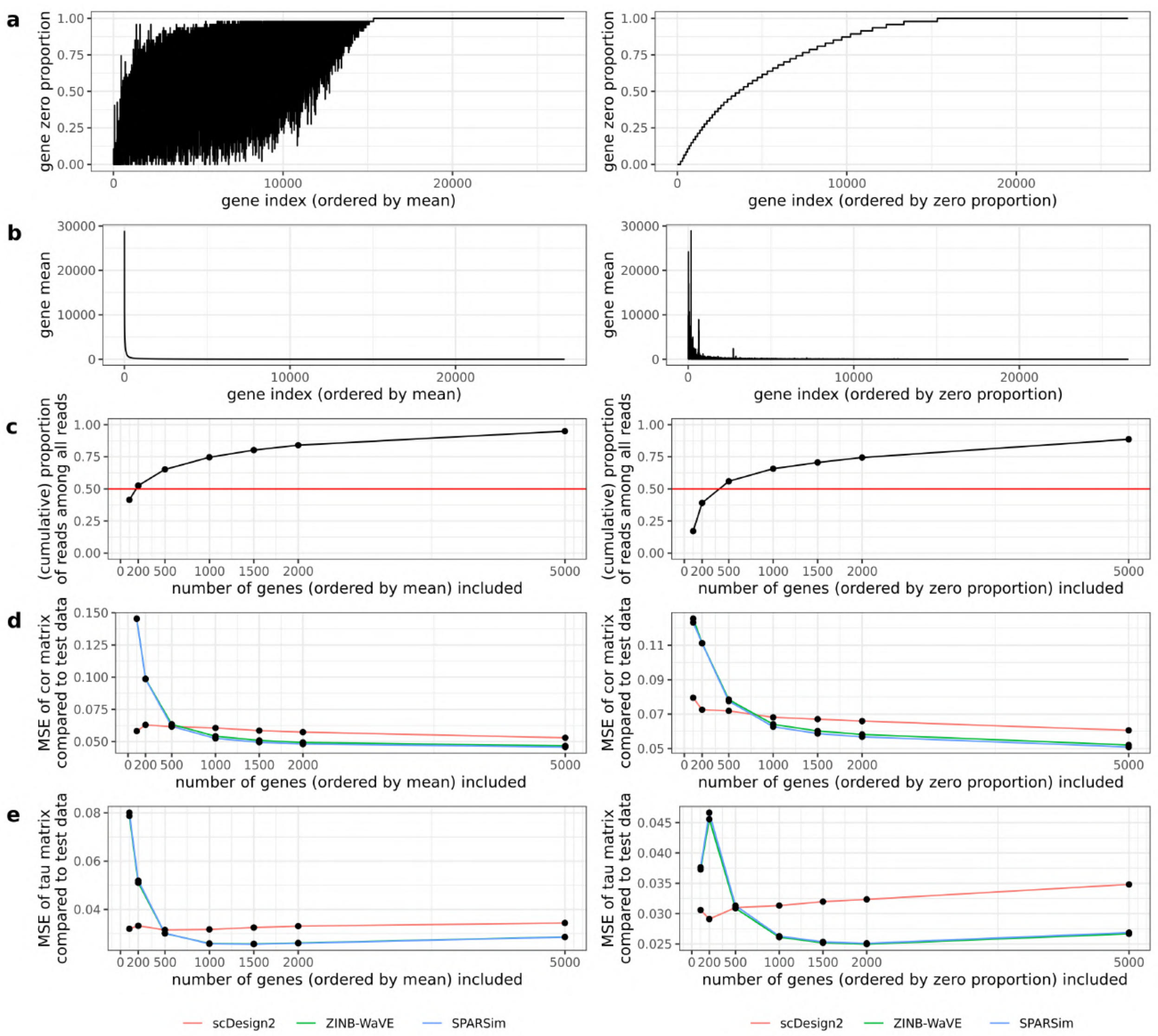


**Figure S27: Relationships of the mean squared error (MSE) vs. the dimension (i.e., number of genes) of the (Pearson or Kendall's tau) gene correlation matrices, which are estimated from the synthetic data generated by three simulators trained on the Smart-Seq2 dendrocyte (subtype 2) data.** The genes are ordered in two ways, by their mean expression from high to low (left) or by their zero proportion from low to high (right). The plots are shown for the top 100, 200, 500, 1000, 1500, and 2000 genes ordered in either way. (a) The zero proportion of each top gene. (b) The mean expression of each top gene. (c) The relationship of the cumulative proportion of the top genes' UMIs among all UMIs vs. the number of top genes. (d)–(e) The MSE is defined as the average per-entry squared difference between the correlation matrices estimated from each synthetic dataset and the test data. Each plot shows the relationship of the MSE of (d) the Pearson correlation matrix or (e) the Kendall's tau matrix for the top genes estimated from each simulator's synthetic data vs. the number of top genes.

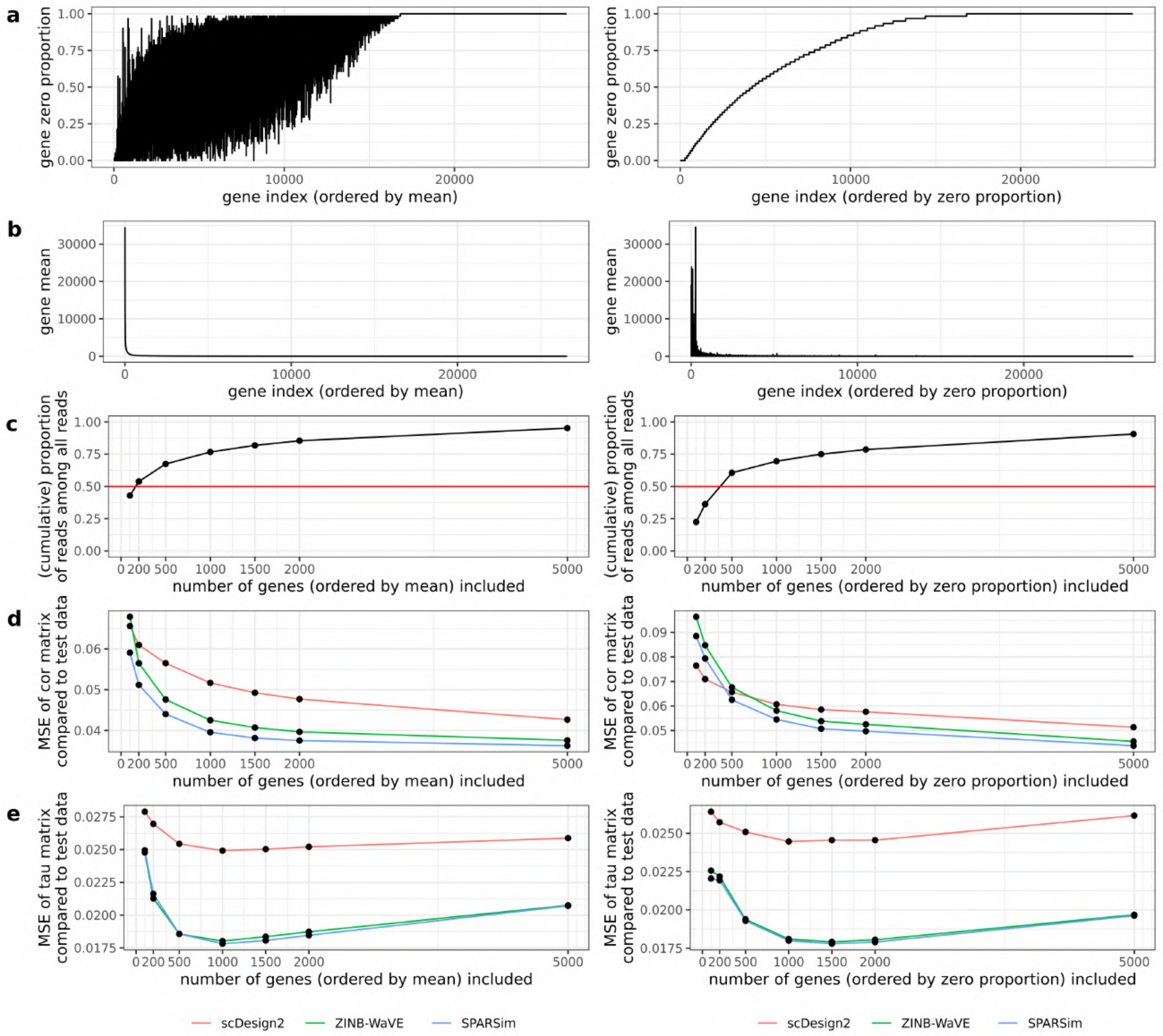


**Figure S28: Relationships of the mean squared error (MSE) vs. the dimension (i.e., number of genes) of the (Pearson or Kendall's tau) gene correlation matrices, which are estimated from the synthetic data generated by three simulators trained on the Smart-Seq2 monocyte (subtype 2) data.** The genes are ordered in two ways, by their mean expression from high to low (left) or by their zero proportion from low to high (right). The plots are shown for the top 100, 200, 500, 1000, 1500, and 2000 genes ordered in either way. (a) The zero proportion of each top gene. (b) The mean expression of each top gene. (c) The relationship of the cumulative proportion of the top genes' UMIs among all UMIs vs. the number of top genes. (d)–(e) The MSE is defined as the average per-entry squared difference between the correlation matrices estimated from each synthetic dataset and the test data. Each plot shows the relationship of the MSE of (d) the Pearson correlation matrix or (e) the Kendall's tau matrix for the top genes estimated from each simulator's synthetic data vs. the number of top genes.

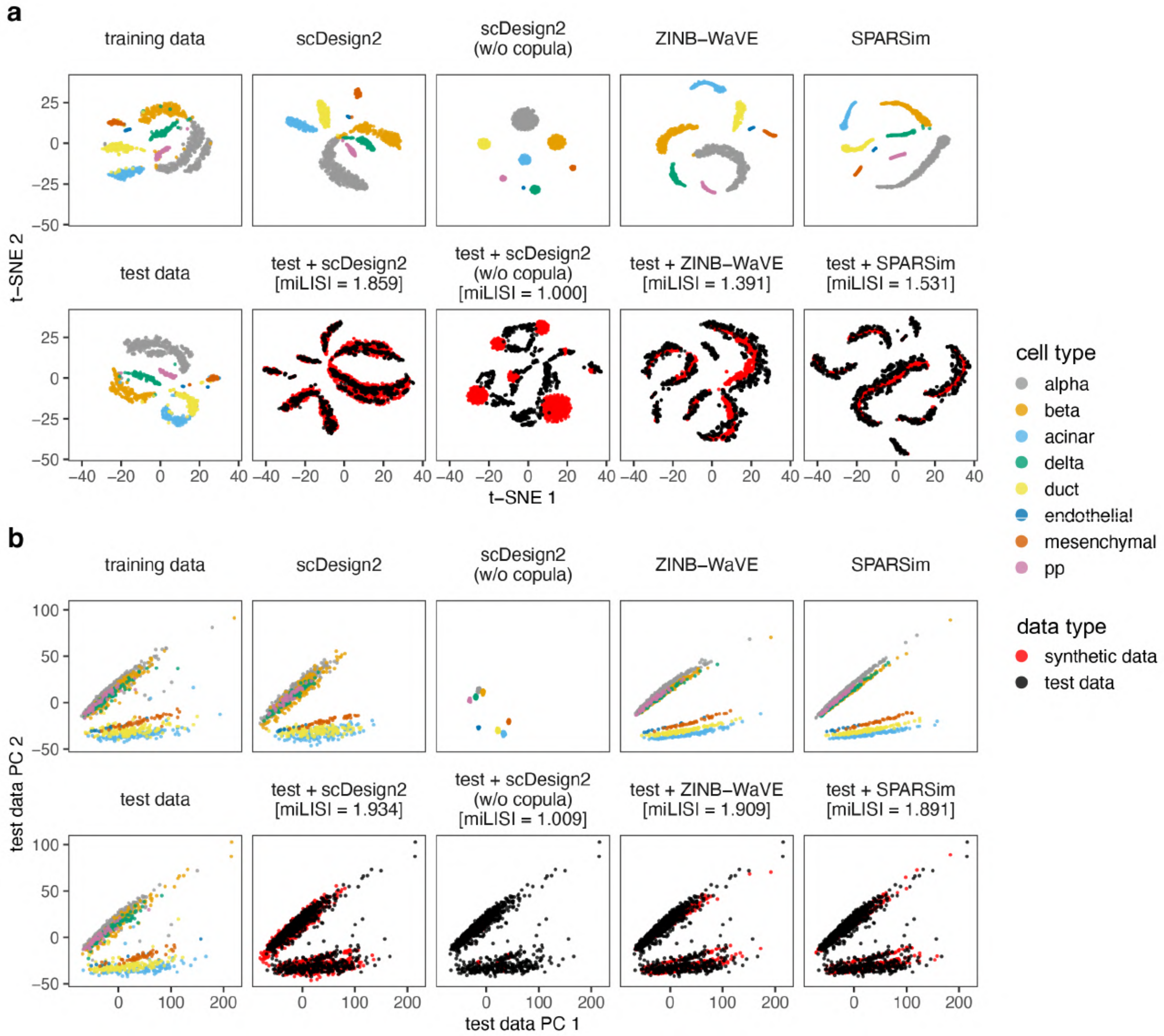


**Figure S29: Comparison of CEL-Seq2 data and synthetic data generated by scDesign2, its variant without copula, ZINB-WaVE, and SPARSim in 2D visualization.** (a) t-SNE plots and (b) principle component (PC) plots of training data, test data, synthetic data generated by each simulator, and combinations of test data and each synthetic dataset. Gene expression counts are transformed as $\log(1 + \text{count})$ before dimensionality reduction. We find that the synthetic data generated by scDesign2 most resemble the test data.

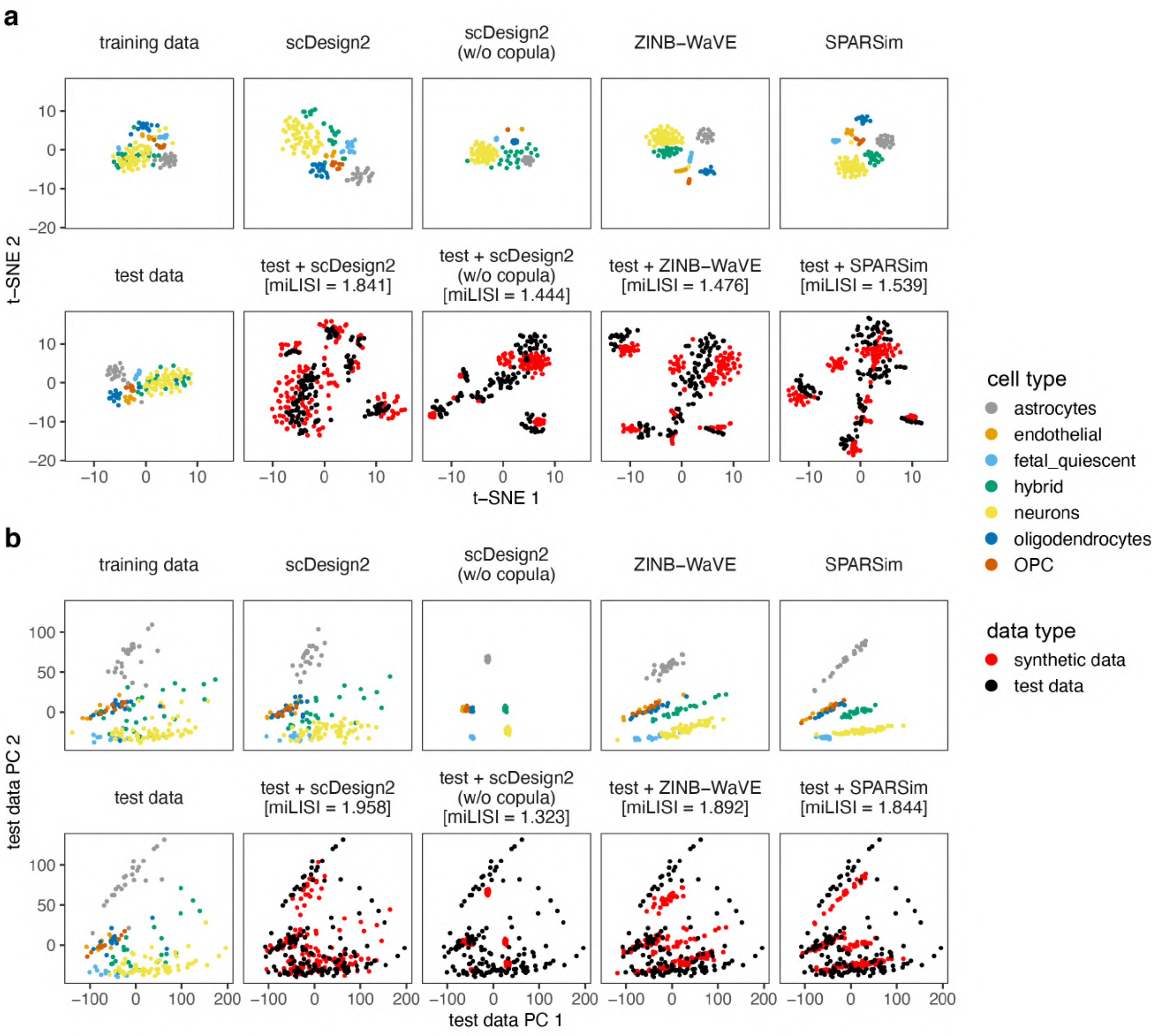


**Figure S30: Comparison of Fluidigm C1 (SMARTer) data and synthetic data generated by scDesign2, its variant without copula, ZINB-WaVE, and SPARSim in 2D visualization.** (a) t-SNE plots and (b) principle component (PC) plots of training data, test data, synthetic data generated by each simulator, and combinations of test data and each synthetic dataset. Gene expression counts are transformed as $\log(1 + \text{count})$ before dimensionality reduction. We find that the synthetic data generated by scDesign2 most resemble the test data.

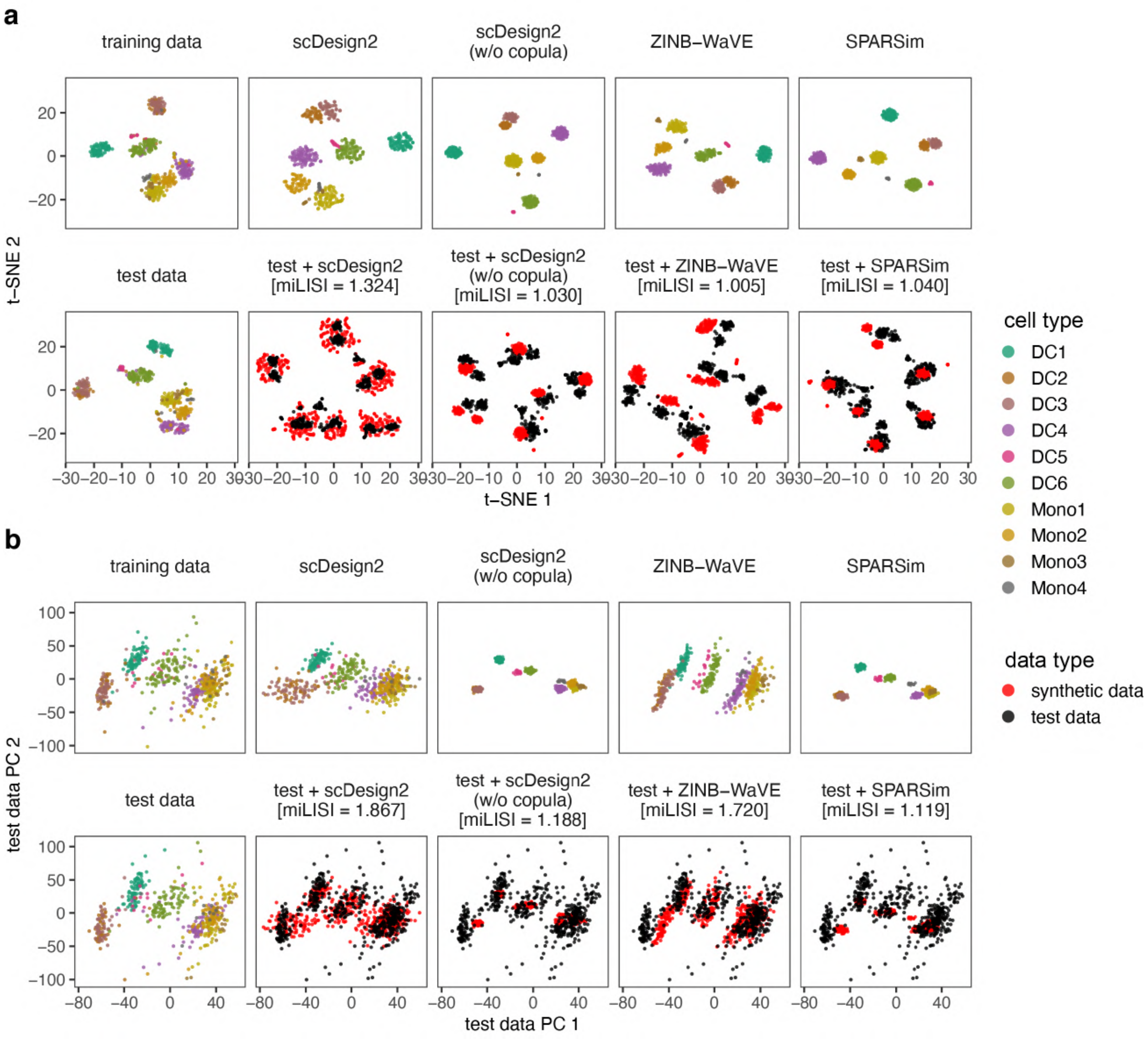


**Figure S31: Comparison of Smart-Seq2 data and synthetic data generated by scDesign2, its variant without copula, ZINB-WaVE, and SPARSim in 2D visualization.** (a) t-SNE plots and (b) principle component (PC) plots of training data, test data, synthetic data generated by each simulator, and combinations of test data and each synthetic dataset. Gene expression counts are transformed as $\log(1 + \text{count})$ before dimensionality reduction. We find that the synthetic data generated by scDesign2 most resemble the test data.

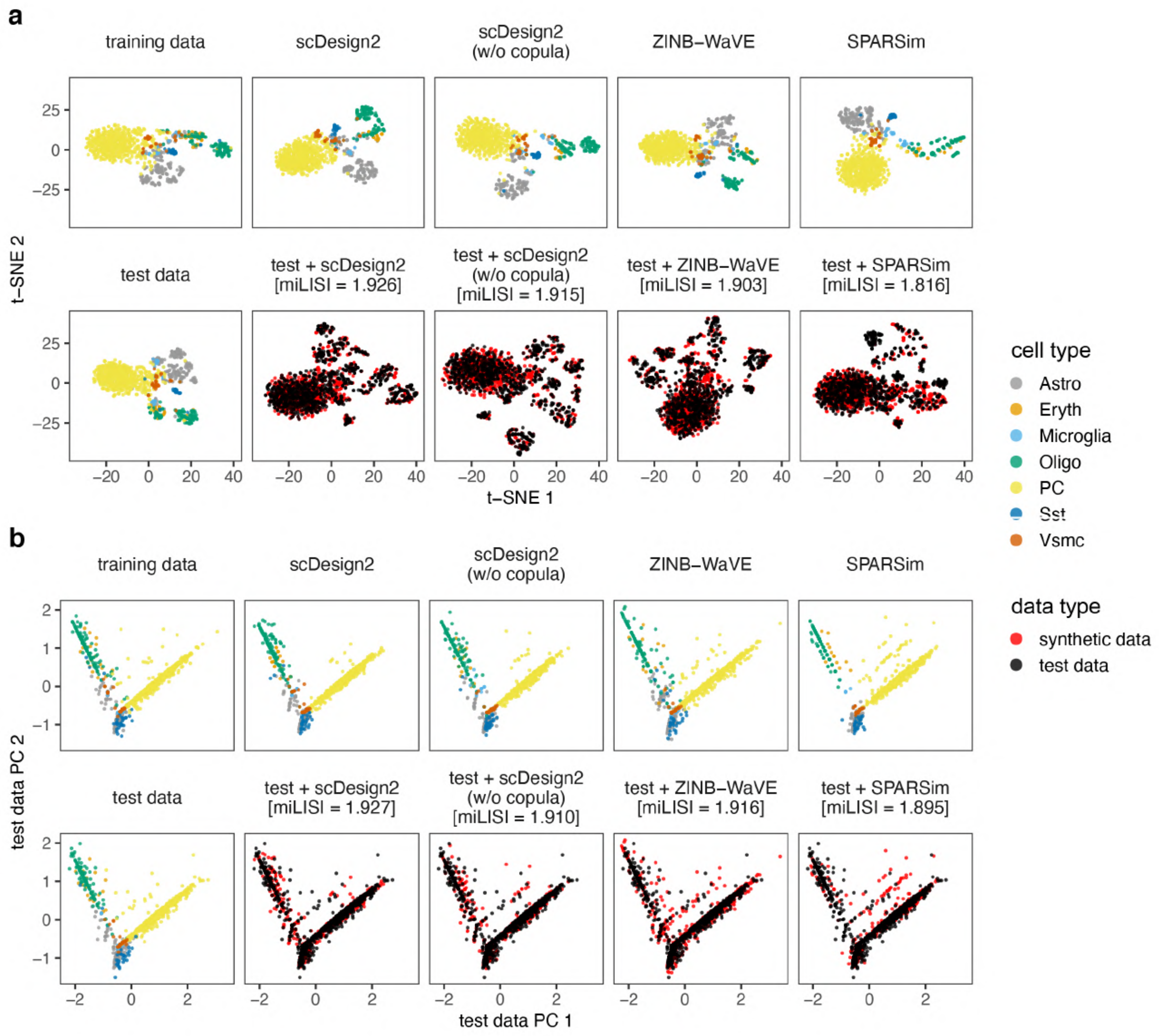


**Figure S32: Comparison of pciSeq data and synthetic data generated by scDesign2, its variant without copula, ZINB-WaVE, and SPARSim in 2D visualization.** (a) t-SNE plots and (b) principle component (PC) plots of training data, test data, synthetic data generated by each simulator, and combinations of test data and each synthetic dataset. Gene expression counts are transformed as $\log(1 + \text{count})$ before dimensionality reduction. miLISI is short for median integration local inverse Simpson's Index, a higher value of which indicates that the simulated data mix better with the test data in the 2D visualization plot. By visually inspecting the patterns in these plots as well as comparing the miLISI values, we find that the synthetic data generated by scDesign2 most resemble the test data.

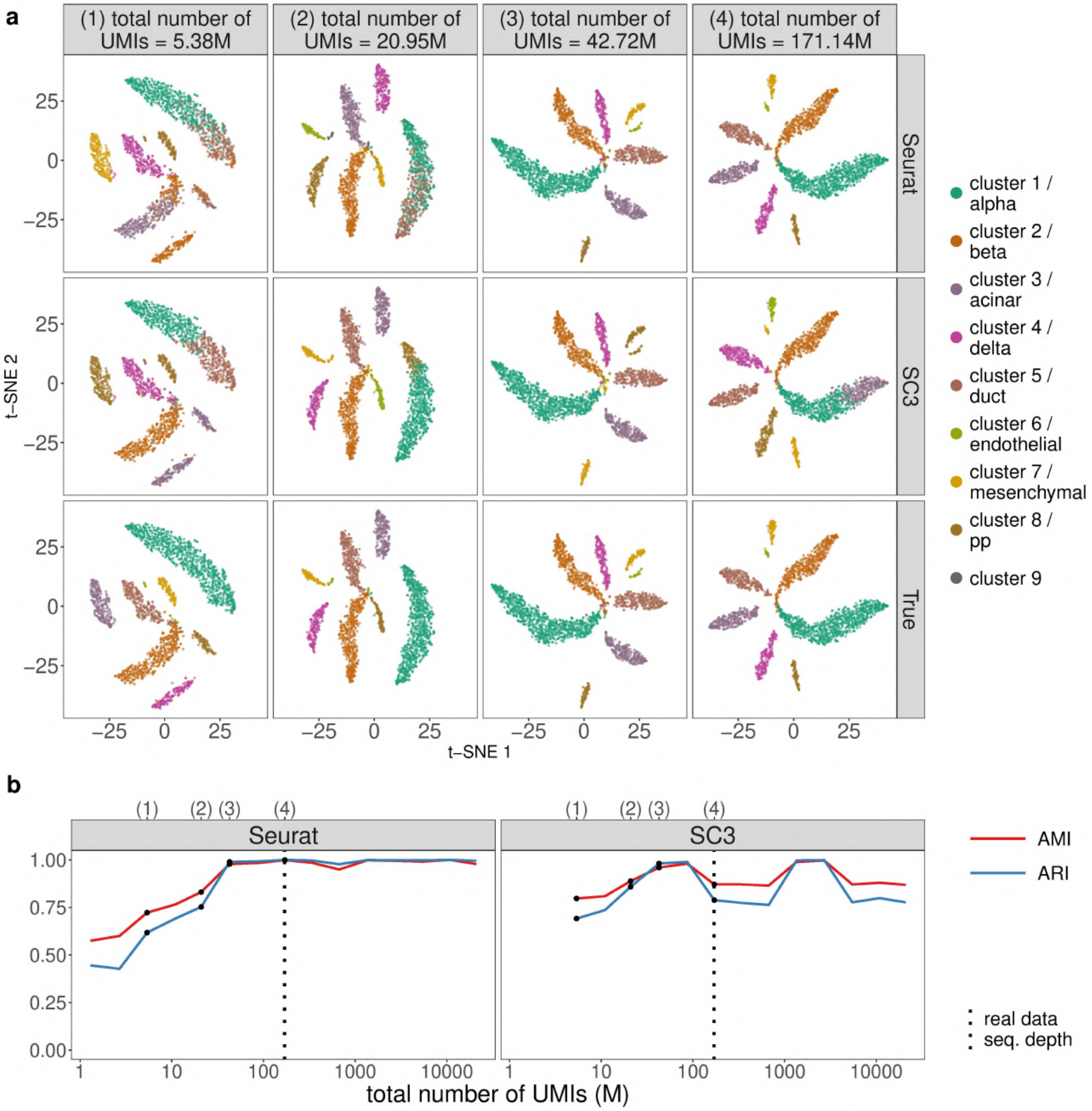


**Figure S33: scDesign2 guides the choice of sequencing depth in cell clustering.** scDesign2 generates synthetic CEL-Seq2 data with fifteen sequencing depths. Two cell clustering methods—Seurat and SC3—are applied to each synthetic dataset to partition cells into cell clusters. (a) t-SNE visualization of four synthetic datasets, where cells are labelled by Seurat clusters (top), SC3 clusters (middle), and annotated cell types (bottom). (b) Two clustering accuracy measures (AMI and ARI) vs. sequencing depth; left: Seurat; right: SC3. In (b), the results of the four sequencing depths in (a) are marked as dots and in the top, and the sequencing depth of the real dataset [77] is marked as vertical dashed lines.

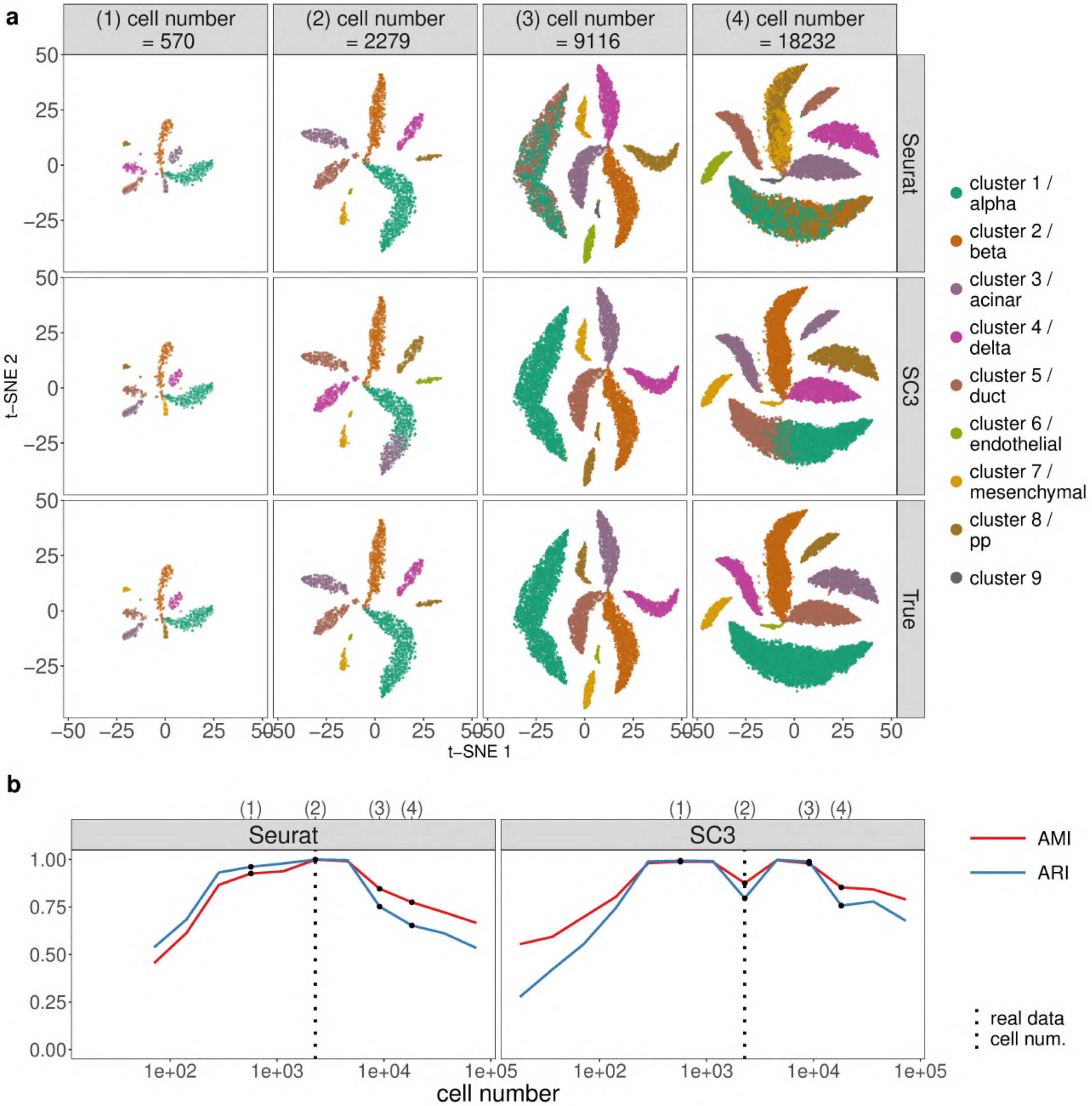


**Figure S34: scDesign2 guides the choice of cell number in cell clustering, in the case where the total sequencing depth is kept as fixed.** scDesign2 generates synthetic CEL-Seq2 data with thirteen cell numbers. Two cell clustering methods—Seurat and SC3—are applied to each synthetic dataset to partition cells into cell clusters. (a) t-SNE visualization of four synthetic datasets, where cells are labelled by Seurat clusters (top), SC3 clusters (middle), and annotated cell types (bottom). (b) Two clustering accuracy measures (AMI and ARI) vs. sequencing depth; left: Seurat; right: SC3. In (b), the results of the four cell numbers in (a) are marked as dots and in the top, and the cell number of the real dataset [77] is marked as vertical dashed lines.

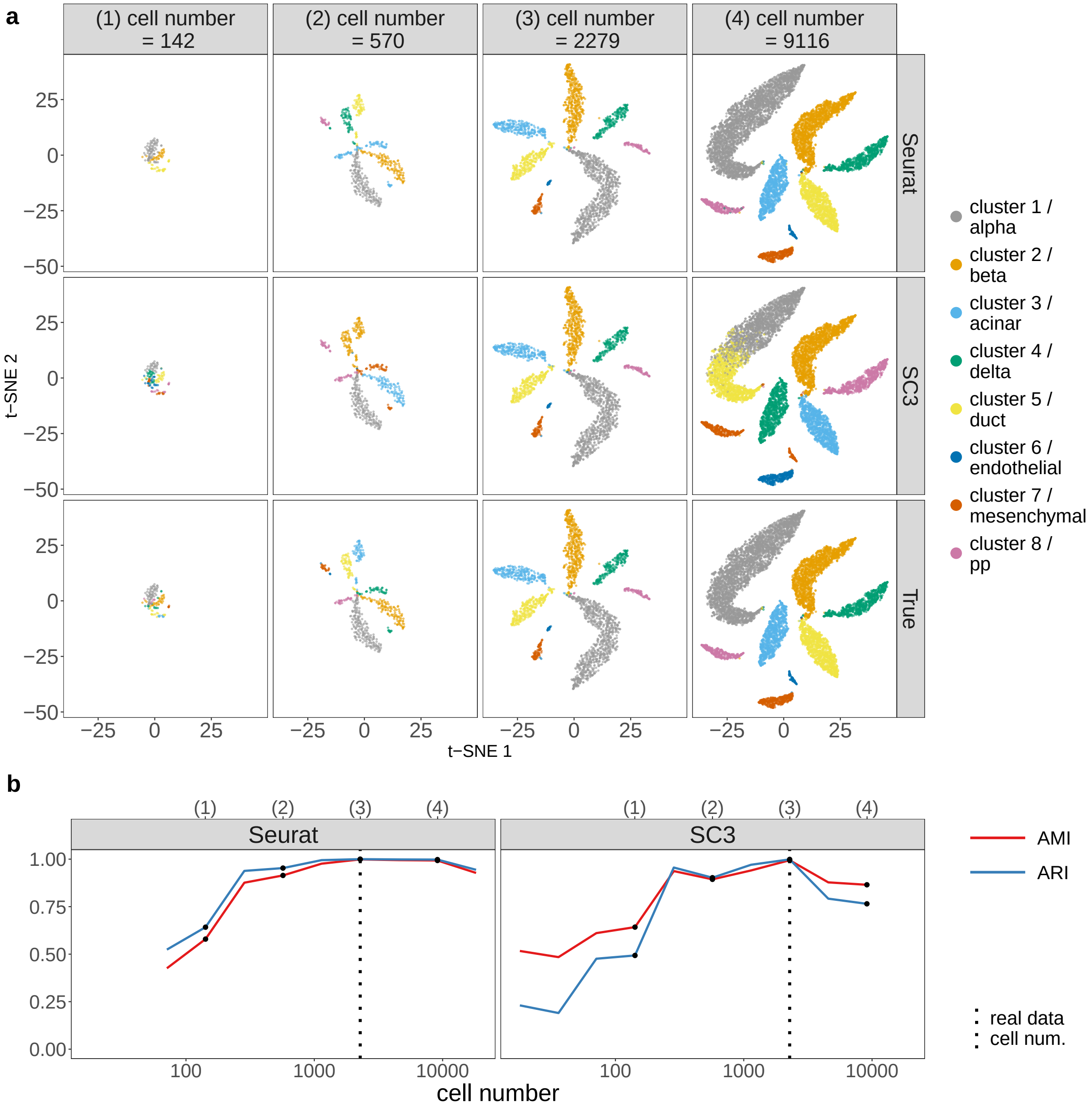


**Figure S35: scDesign2 guides the choice of cell number in cell clustering, in the case where the average sequencing depth is kept as fixed.** scDesign2 generates synthetic CEL-Seq2 data with ten cell numbers. Two cell clustering methods—Seurat and SC3—are applied to each synthetic dataset to partition cells into cell clusters. (a) t-SNE visualization of four synthetic datasets, where cells are labelled by Seurat clusters (top), SC3 clusters (middle), and annotated cell types (bottom). (b) Two clustering accuracy measures (AMI and ARI) vs. sequencing depth; left: Seurat; right: SC3. In (b), the results of the four cell numbers in (a) are marked as dots and in the top, and the cell number of the real dataset [77] is marked as vertical dashed lines.

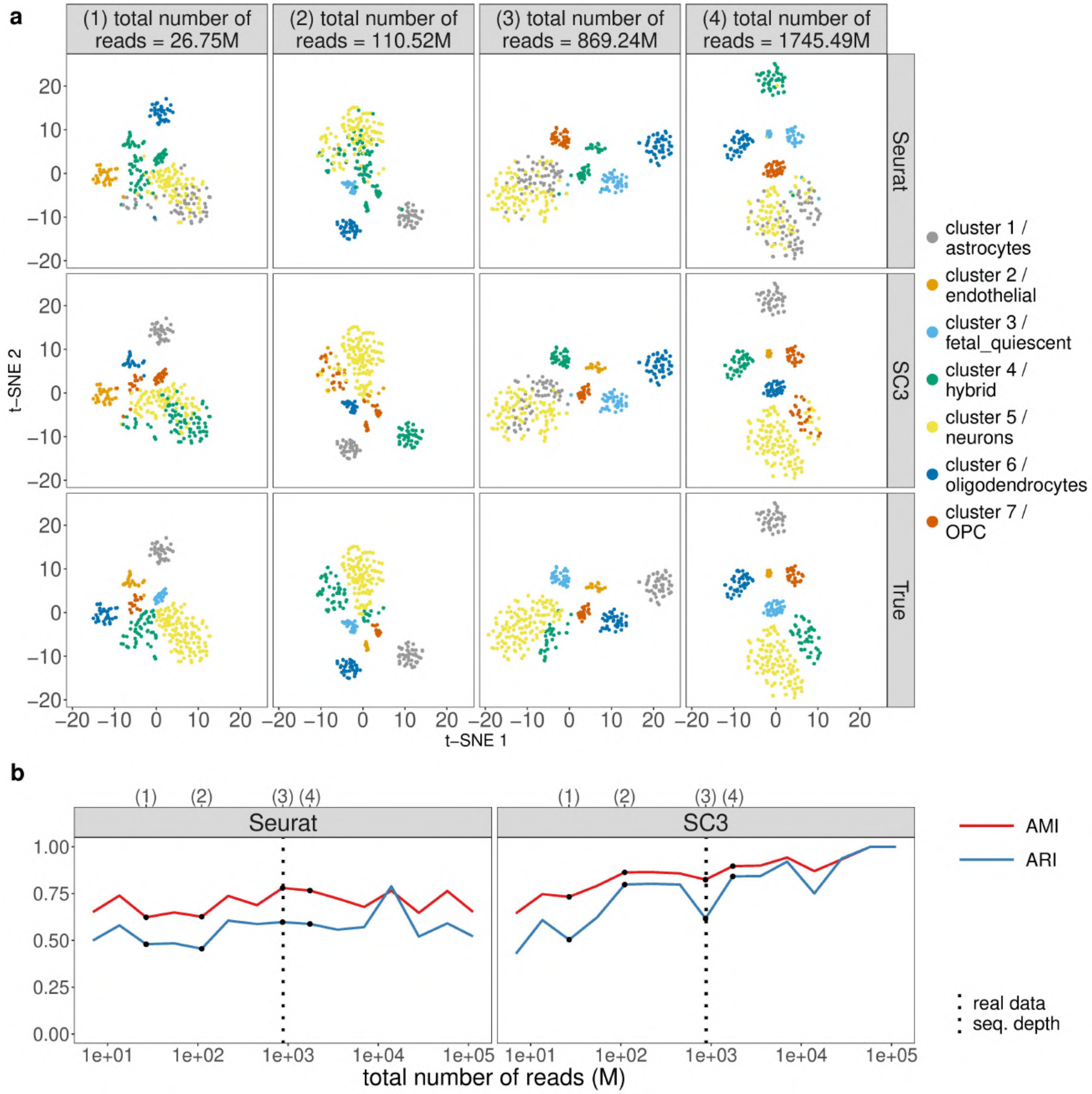


**Figure S36: scDesign2 guides the choice of sequencing depth in cell clustering.** scDesign2 generates synthetic Fluidigm C1 (SMARTer) data with fifteen sequencing depths. Two cell clustering methods—Seurat and SC3—are applied to each synthetic dataset to partition cells into cell clusters. (a) t-SNE visualization of four synthetic datasets, where cells are labelled by Seurat clusters (top), SC3 clusters (middle), and annotated cell types (bottom). (b) Two clustering accuracy measures (AMI and ARI) vs. sequencing depth; left: Seurat; right: SC3. In (b), the results of the four sequencing depths in (a) are marked as dots and in the top, and the sequencing depth of the real dataset [78] is marked as vertical dashed lines.

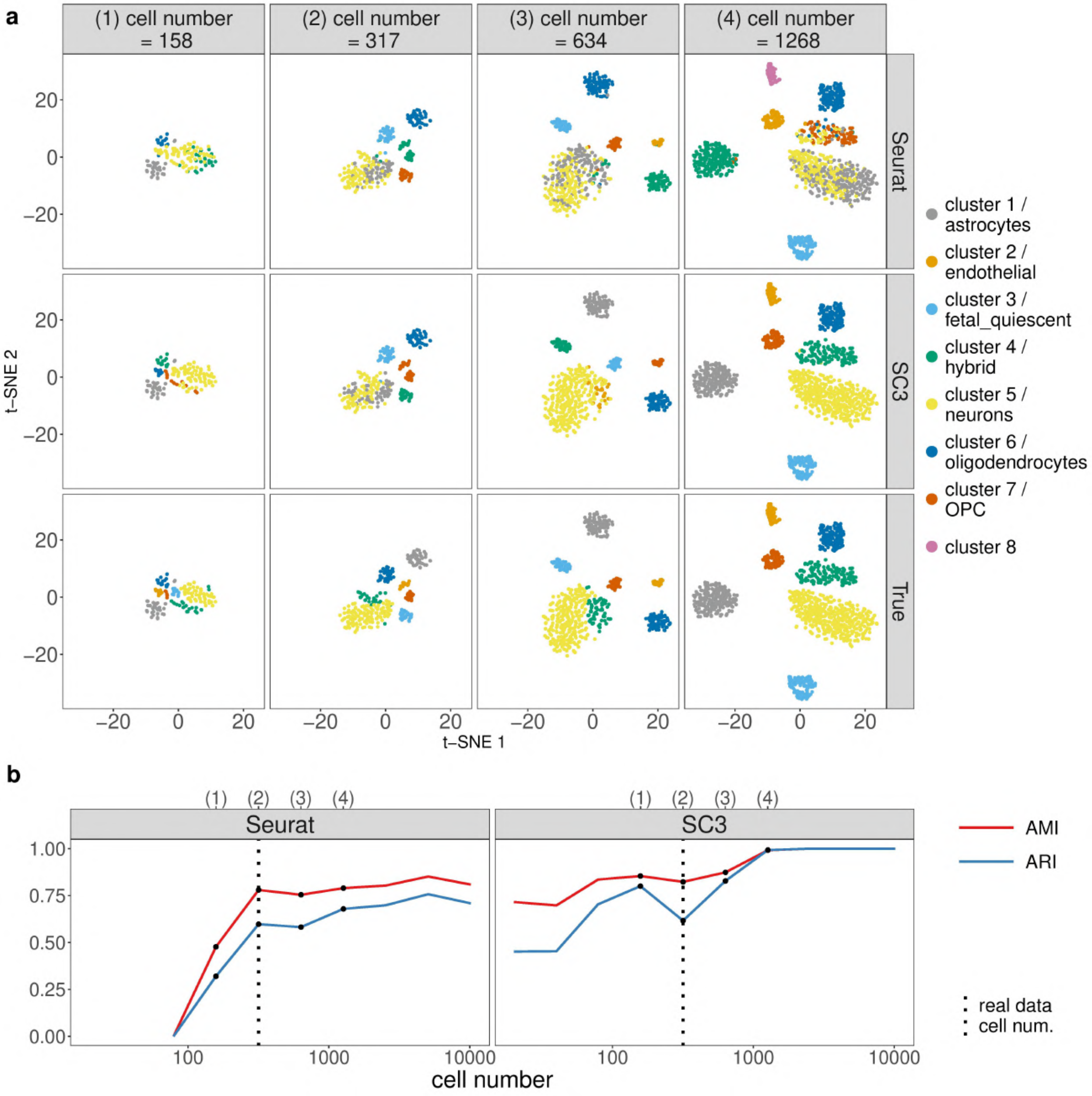


**Figure S37: scDesign2 guides the choice of cell number in cell clustering, in the case where the total sequencing depth is kept as fixed.** scDesign2 generates synthetic Fluidigm C1 (SMARTer) data with ten cell numbers. Two cell clustering methods—Seurat and SC3—are applied to each synthetic dataset to partition cells into cell clusters. (a) t-SNE visualization of four synthetic datasets, where cells are labelled by Seurat clusters (top), SC3 clusters (middle), and annotated cell types (bottom). (b) Two clustering accuracy measures (AMI and ARI) vs. sequencing depth; left: Seurat; right: SC3. In (b), the results of the four cell numbers in (a) are marked as dots and in the top, and the cell number of the real dataset [78] is marked as vertical dashed lines.

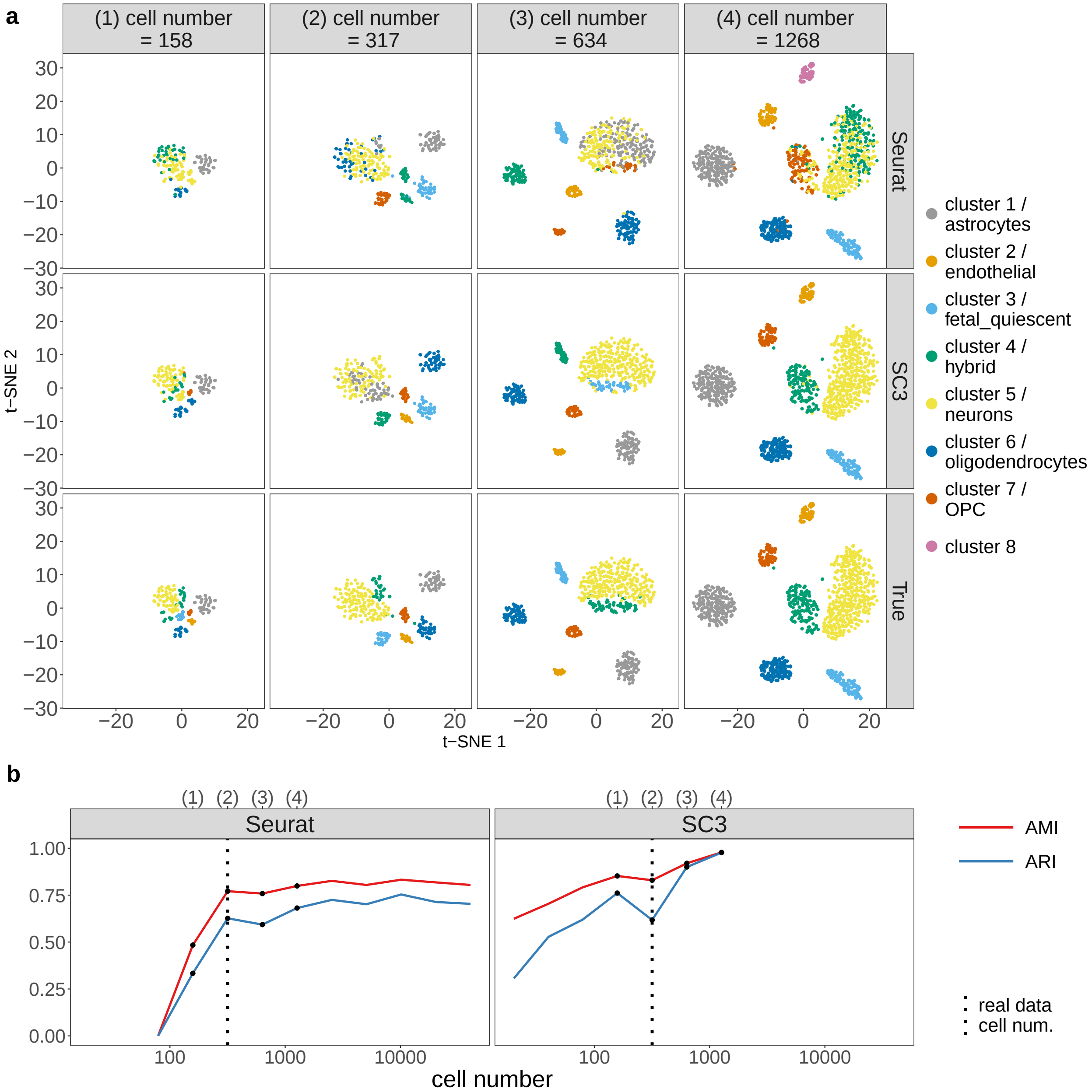


**Figure S38: scDesign2 guides the choice of cell number in cell clustering, in the case where the average sequencing depth is kept as fixed.** scDesign2 generates synthetic Fluidigm C1 (SMARTer) data with twelve cell numbers. Two cell clustering methods—Seurat and SC3—are applied to each synthetic dataset to partition cells into cell clusters. (a) t-SNE visualization of four synthetic datasets, where cells are labelled by Seurat clusters (top), SC3 clusters (middle), and annotated cell types (bottom). (b) Two clustering accuracy measures (AMI and ARI) vs. sequencing depth; left: Seurat; right: SC3. In (b), the results of the four cell numbers in (a) are marked as dots and in the top, and the cell number of the real dataset [78] is marked as vertical dashed lines.

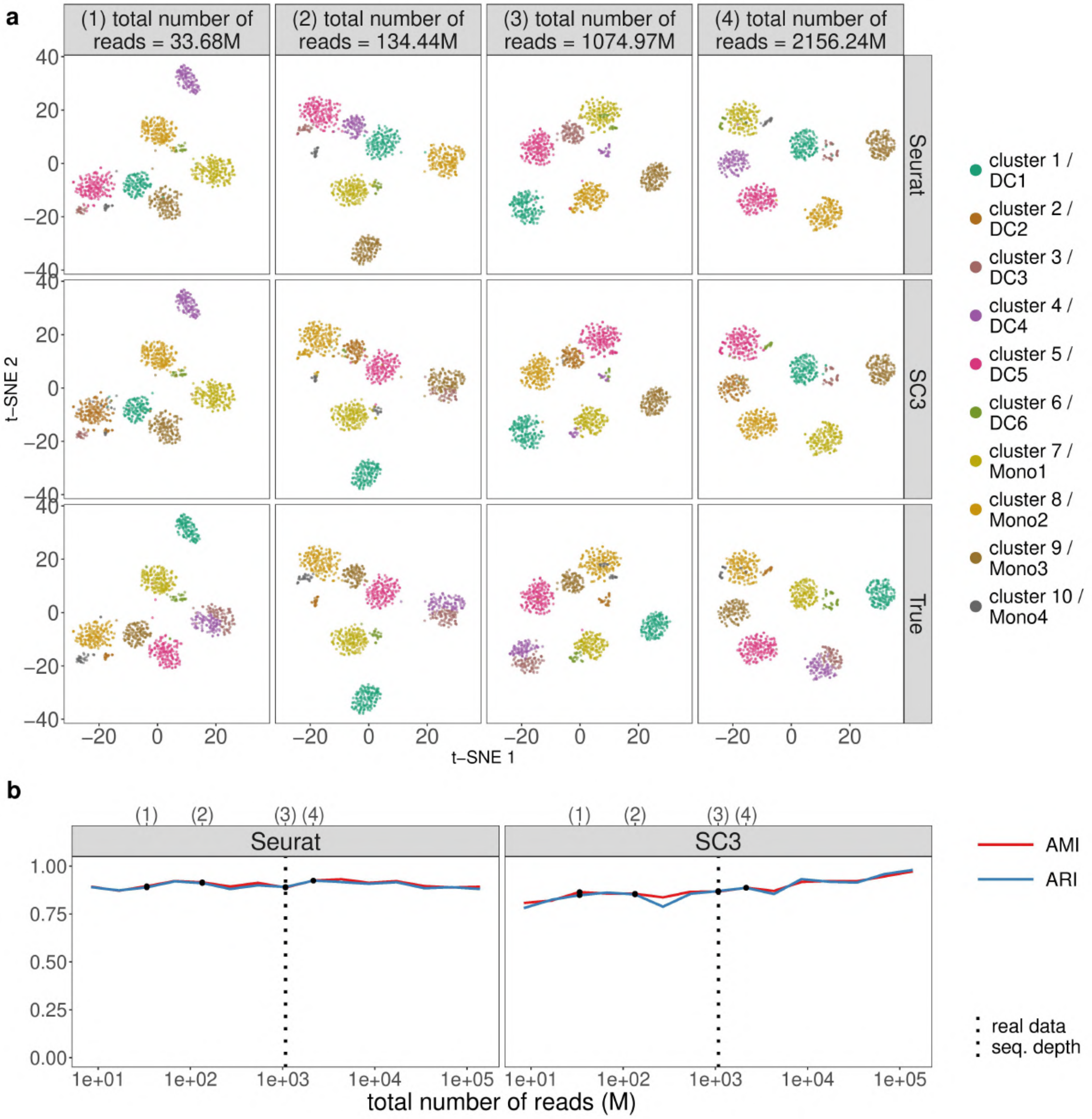


**Figure S39: scDesign2 guides the choice of sequencing depth in cell clustering.** scDesign2 generates synthetic Smart-Seq2 data with fifteen sequencing depths. Two cell clustering methods—Seurat and SC3—are applied to each synthetic dataset to partition cells into cell clusters. (a) t-SNE visualization of four synthetic datasets, where cells are labelled by Seurat clusters (top), SC3 clusters (middle), and annotated cell types (bottom). (b) Two clustering accuracy measures (AMI and ARI) vs. sequencing depth; left: Seurat; right: SC3. In (b), the results of the four sequencing depths in (a) are marked as dots and in the top, and the sequencing depth of the real dataset [79] is marked as vertical dashed lines.

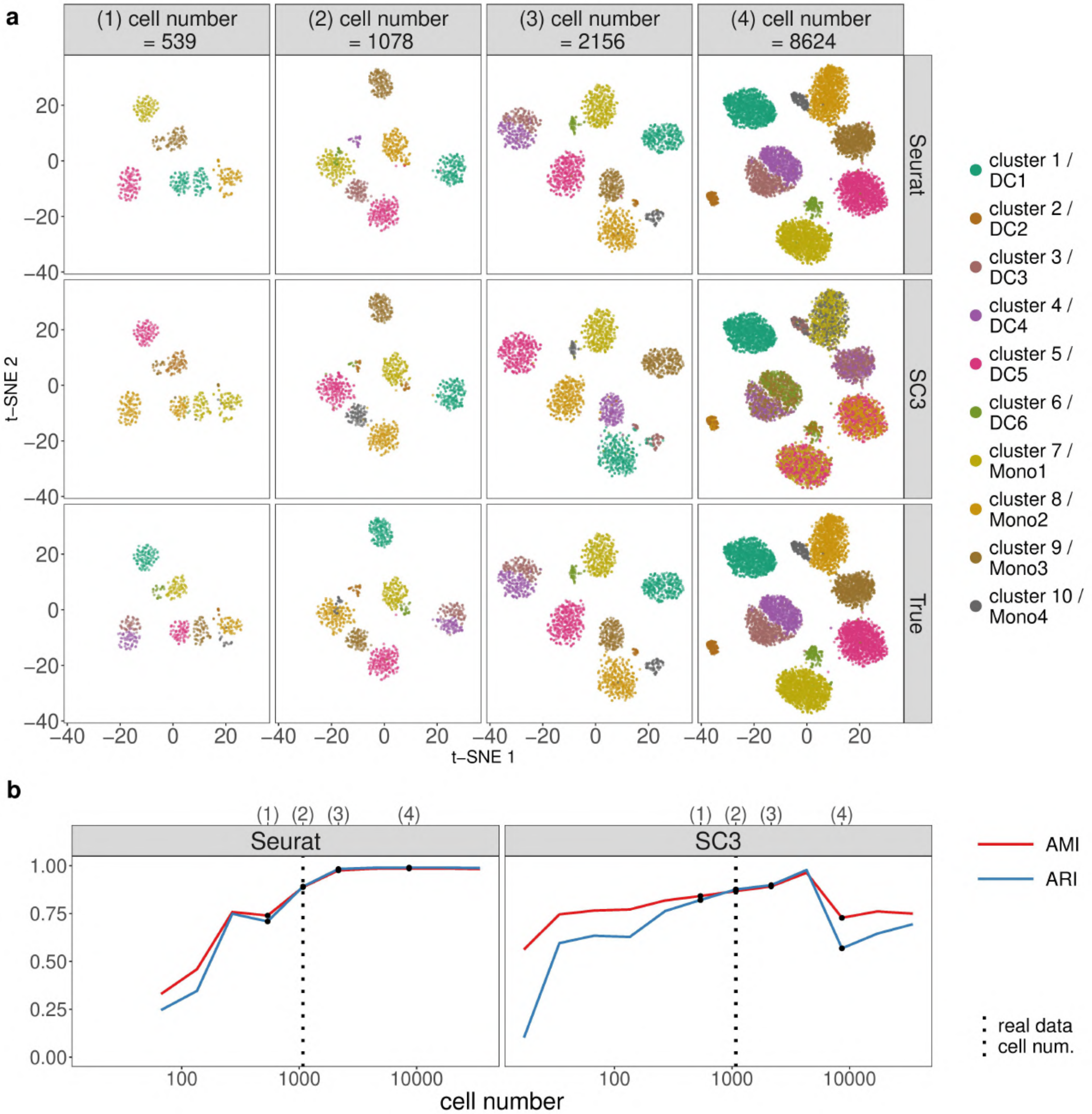


**Figure S40: scDesign2 guides the choice of cell number in cell clustering, in the case where the total sequencing depth is kept as fixed.** scDesign2 generates synthetic Smart-Seq2 data with twelve cell numbers. Two cell clustering methods—Seurat and SC3—are applied to each synthetic dataset to partition cells into cell clusters. (a) t-SNE visualization of four synthetic datasets, where cells are labelled by Seurat clusters (top), SC3 clusters (middle), and annotated cell types (bottom). (b) Two clustering accuracy measures (AMI and ARI) vs. sequencing depth; left: Seurat; right: SC3. In (b), the results of the four cell numbers in (a) are marked as dots and in the top, and the cell number of the real dataset [79] is marked as vertical dashed lines.

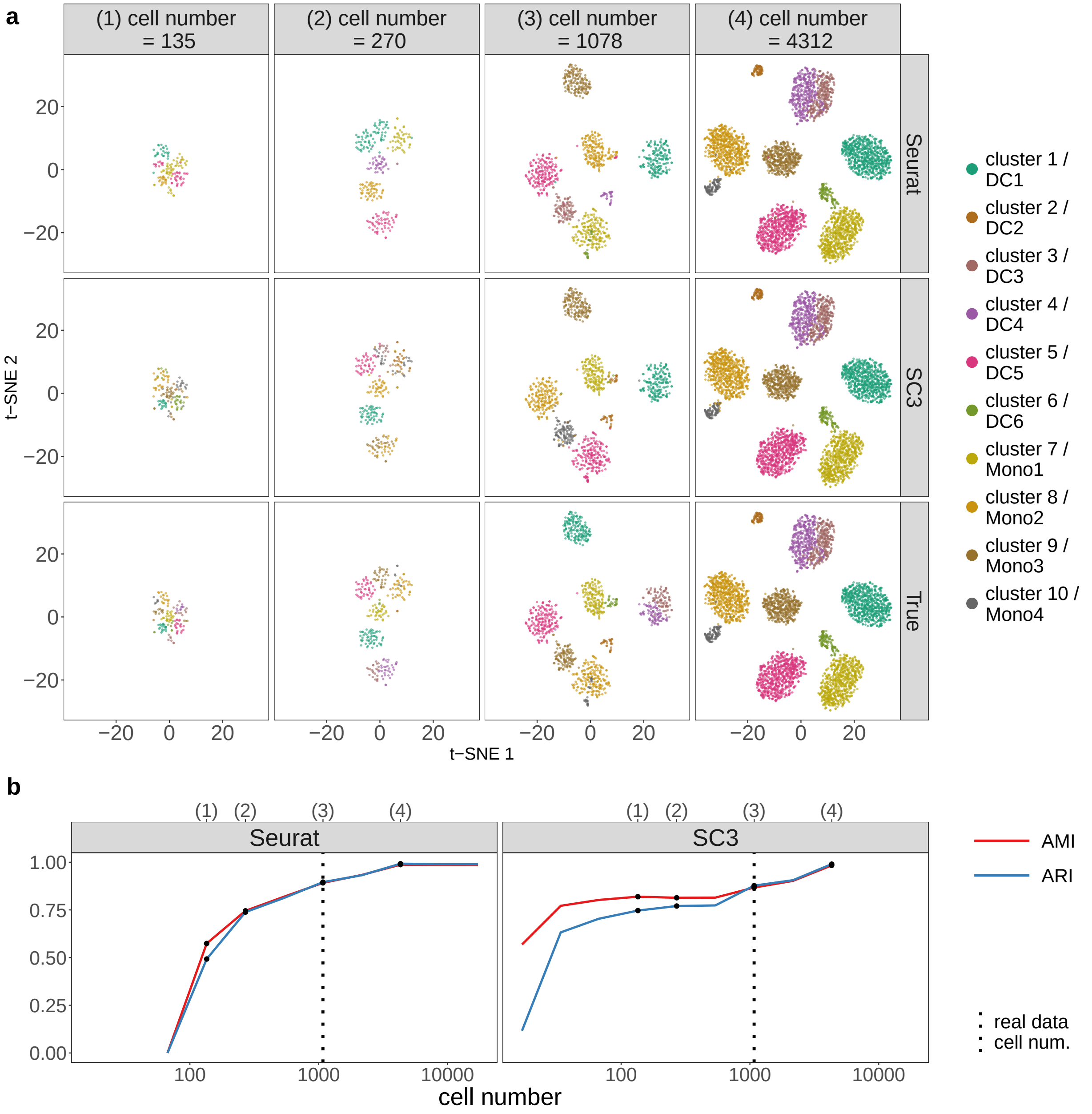


**Figure S41: scDesign2 guides the choice of cell number in cell clustering, in the case where the average sequencing depth is kept as fixed.** scDesign2 generates synthetic Smart-Seq2 data with eleven cell numbers. Two cell clustering methods—Seurat and SC3—are applied to each synthetic dataset to partition cells into cell clusters. (a) t-SNE visualization of four synthetic datasets, where cells are labelled by Seurat clusters (top), SC3 clusters (middle), and annotated cell types (bottom). (b) Two clustering accuracy measures (AMI and ARI) vs. sequencing depth; left: Seurat; right: SC3. In (b), the results of the four cell numbers in (a) are marked as dots and in the top, and the cell number of the real dataset [79] is marked as vertical dashed lines.

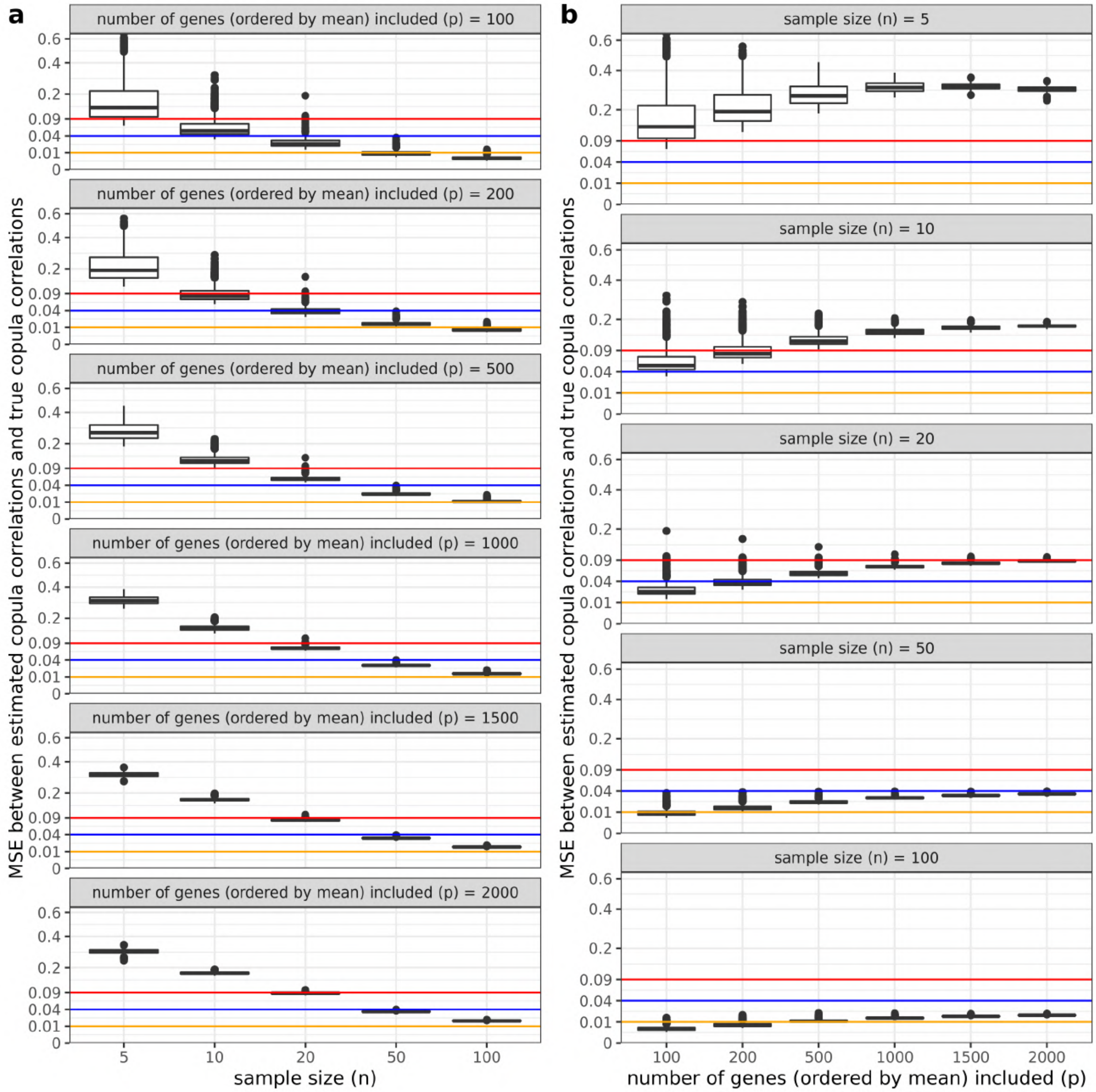


**Figure S42: The effects of $n$ (the sample size, i.e., the number of cells) and $p$ (the number of top highly expressed genes) on the estimation of the copula correlation matrix in the context of 10x Genomics stem cell data.** The mean squared error (MSE) between the estimated copula correlations and the true copula correlations are calculated. For each sample size $n$, 1000 random samples are simulated from a known Gaussian copula model, which is fitted to the 10x Genomics stem cell data, and each sample is then used to estimate the copula correlation matrix. For computational efficiency, we estimate copula correlations using the formula $\rho_z = \sin\left(\frac{\pi}{2}\tau\right)$ by plugging in the sample tau values. (a) The relationship between MSE and $n$, for each $p$ varying from 100 to 2000. (b) The relationship between MSE and $p$, for each $n$ varying from 5 to 100. The vertical axes are on the square-root scale.

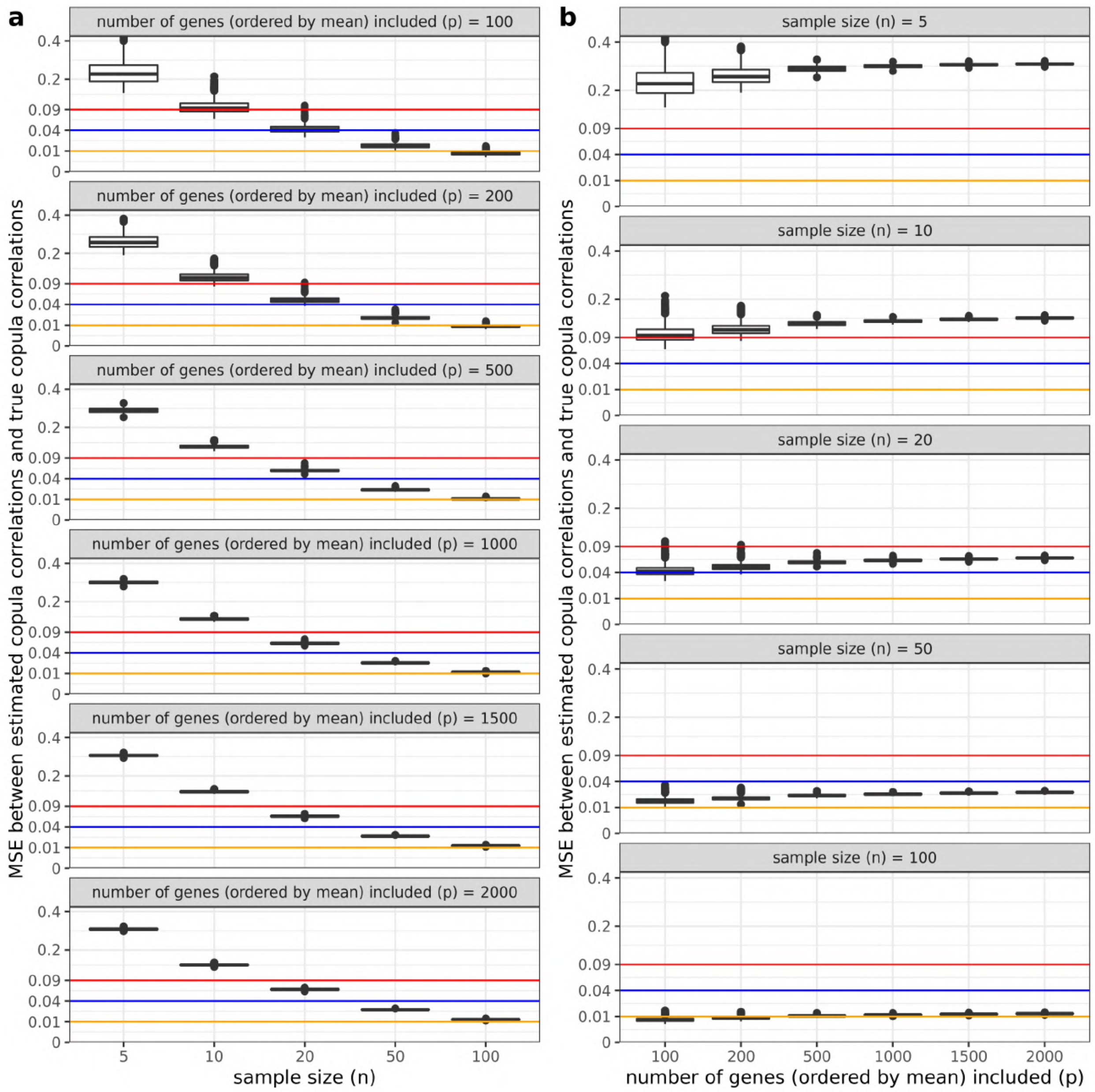


**Figure S43: The effects of $n$ (the sample size, i.e., the number of cells) and $p$ (the number of top highly expressed genes) on the estimation of the copula correlation matrix in the context of Smart-Seq2 dendrocytes (subtype 1) data.** The mean squared error (MSE) between the estimated copula correlations and the true copula correlations are calculated. For each sample size $n$, 1000 random samples are simulated from a known Gaussian copula model, which is fitted to the Smart-Seq2 dendrocytes (subtype 1) data, and each sample is then used to estimate the copula correlation matrix. For computational efficiency, we estimate copula correlations using the formula $\rho_z = \sin\left(\frac{\pi}{2}\tau\right)$ by plugging in the sample tau values. (a) The relationship between MSE and $n$, for each $p$ varying from 100 to 2000. (b) The relationship between MSE and $p$, for each $n$ varying from 5 to 100. The vertical axes are on the square-root scale.

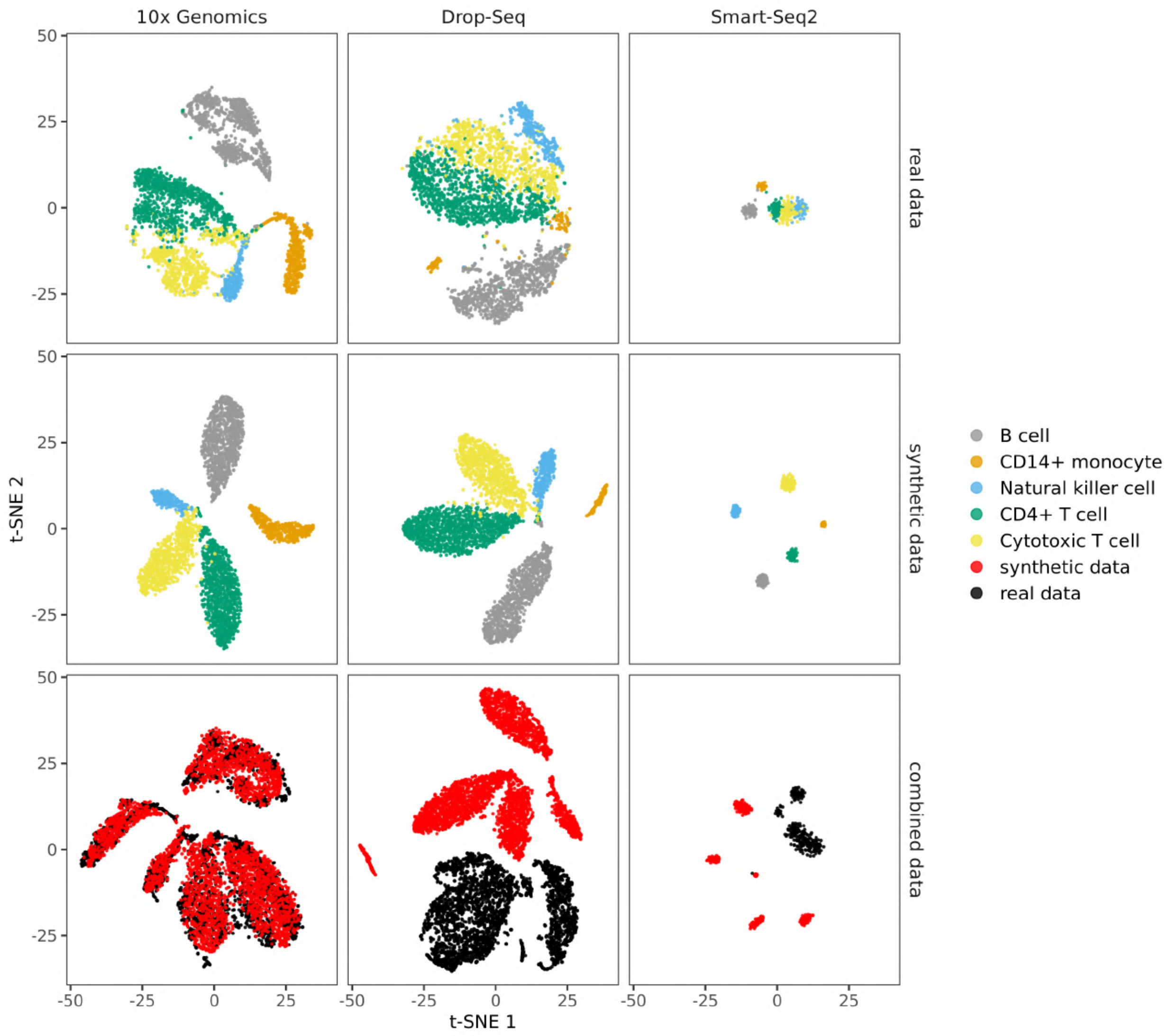


**Figure S44: 2D t-SNE visualization of the results of a cross-platform simulation experiment.** We use a multi-protocol dataset of peripheral blood mononuclear cells (PBMCs) generated for benchmarking purposes [19]. We select data of five cell types measured by three protocols, 10x Genomics, Drop-Seq, and Smart-Seq2, and we train scDesign2 on the 10x Genomics data. Then we adjust the fitted scDesign2 model for the Drop-Seq and Smart-Seq2 protocols by rescaling the mean parameters in the fitted model (see Methods for details). After the adjustment, we use the model for each protocol to generate synthetic data. The 2D t-SNE visualization plot is generated for the real data, the synthetic data, and the combination of the real data and the synthetic data, for each of the three protocols. From the combination plot, we can see that the synthetic cells do not mix well with the real cells for the two cross-protocol scenarios; only for 10x Genomics, the same-protocol scenario, the synthetic cells mix well with the real cells. As a quantitative comparison, the median integration local inverse Simpson's Index (miLISI) for the three combination plots are $1.756$, $1.000$, and $1.000$, from left to right, with the largest value indicating the best mixing for the same-protocol scenario, confirming our conclusion.

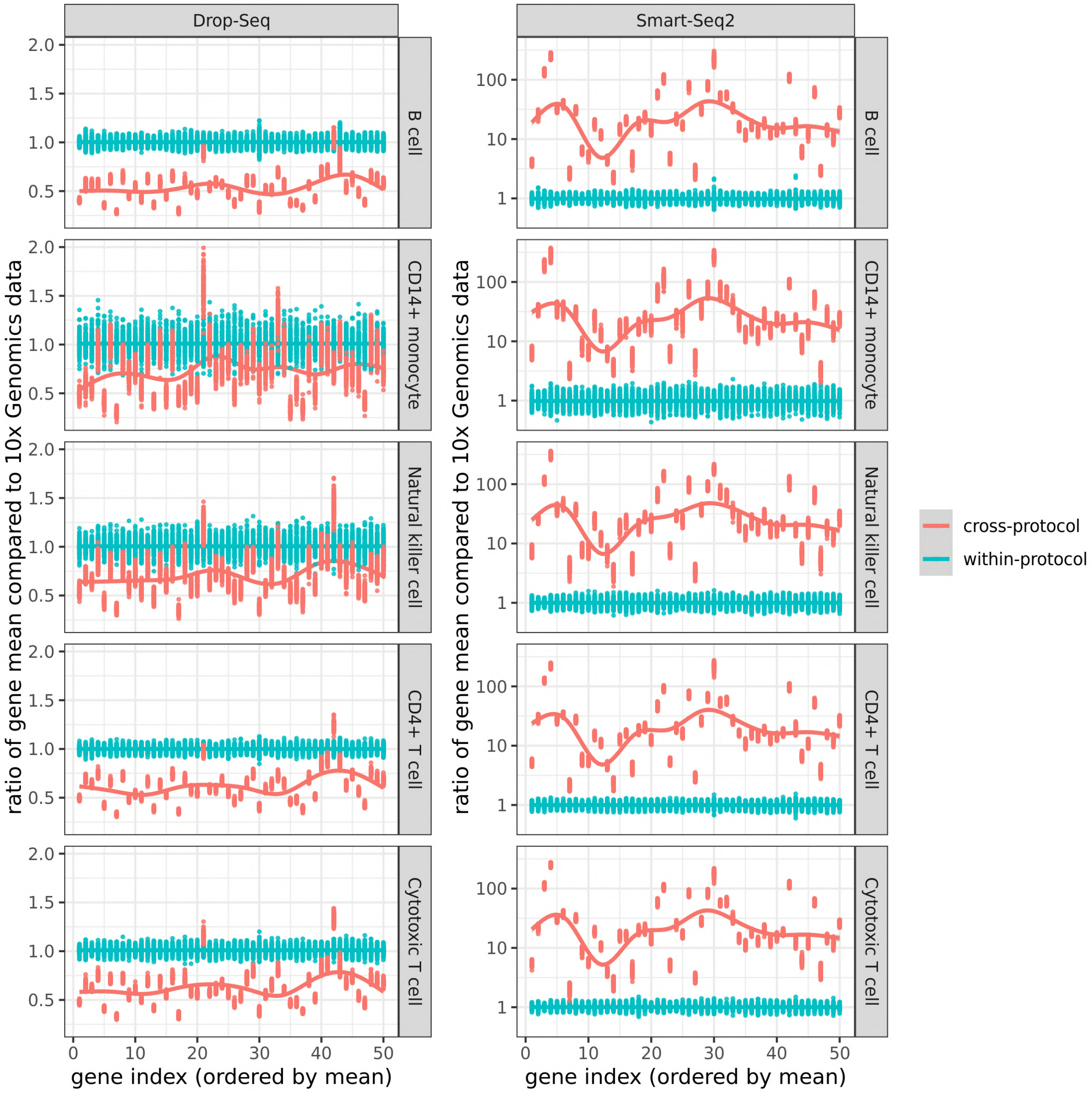


**Figure S45: The cross-protocol and within-protocol ratios of genes' mean expression levels between a target protocol (Drop-Seq or Smart-Seq2) and the reference protocol (10x Genomics) in five cell types.** The results are shown for the top 50 highly expressed genes across the five cell types in the reference protocol, ordered by mean expression from high to low. In each cell type (row) and each target protocol (column), a gene has 100 cross-protocol ratios and 100 within-protocol ratios as a result of random partitioning. To illustrate the trends of the ratios, smoothed curves are added by the R function `geom_smooth()`. The calculation detail is as follows. For each cell type, suppose the target protocol has $m$ cells, and the reference protocol has $n$ cells. In each random partition, we first randomly select $\frac{\min(m,n)}{2}$ cells from the target protocol and the reference protocol each, and compute a gene's two means respectively. Second, for the remaining cells in the reference protocol, we randomly select $\frac{n}{2}$ cells and compute the same gene's mean as a reference. Third, we calculate the cross-protocol ratio and the within-protocol ratio by dividing the first two means by the reference mean. We repeat the above three steps for 100 times for every gene in each cell type and each target protocol. From the plot, we can see that, unlike the within-protocol ratios, which center around the constant value of 1, the cross-protocol ratios fluctuate between genes, and do not center around a constant value. This shows that there does not exist a single scaling factor to convert all genes' expression levels from one protocol to another.

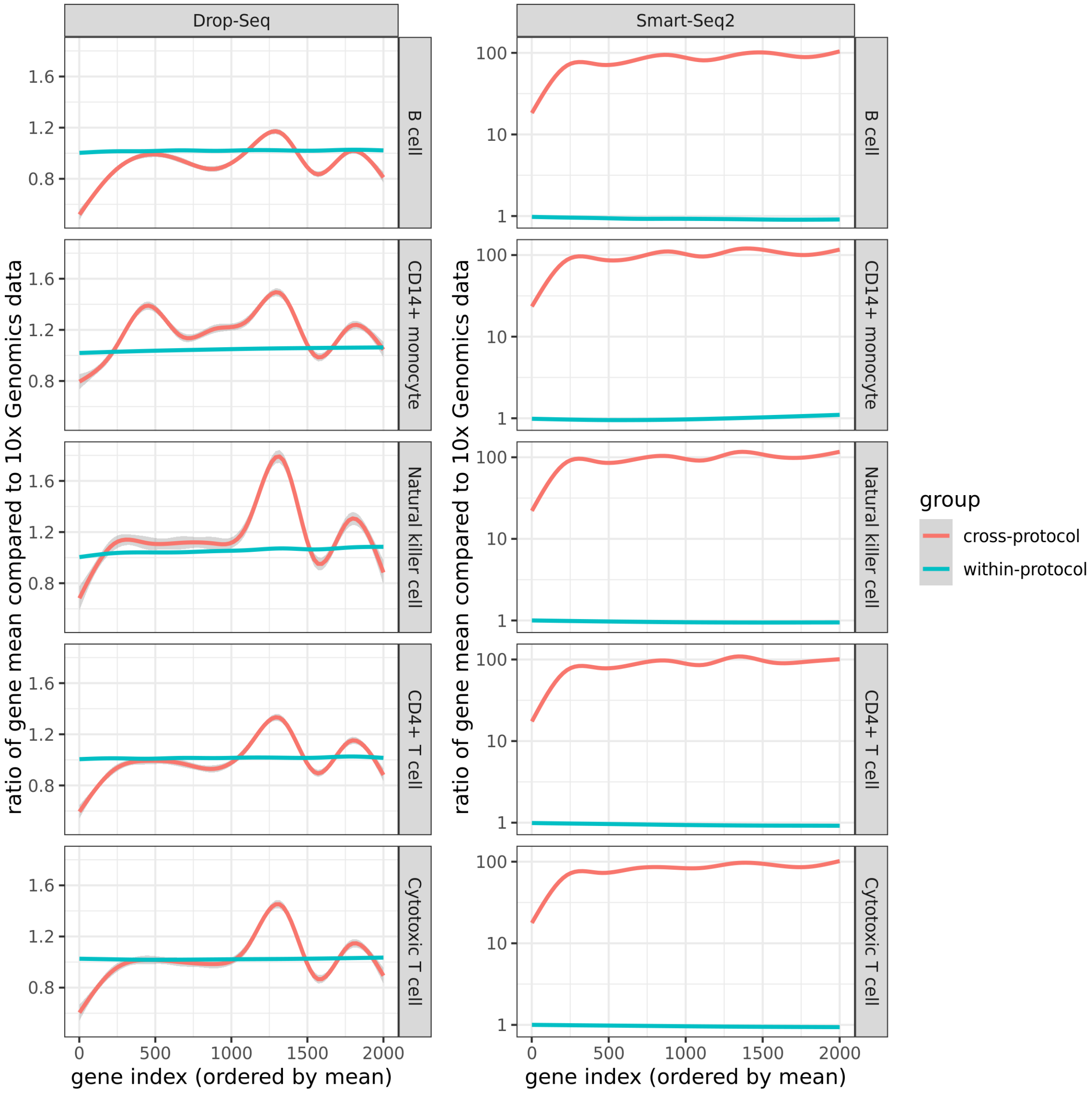


**Figure S46: The cross-protocol and within-protocol ratios of genes' mean expression levels between a target protocol (Drop-Seq or Smart-Seq2) and the reference protocol (10x Genomics) in five cell types.** The results are computed for the top 2000 highly expressed genes in each cell type in the reference protocol, ordered by mean expression from high to low. In each cell type (row) and each target protocol (column), a gene has 100 cross-protocol ratios and 100 within-protocol ratios as a result of random partitioning. For clarity of illustration, only the trends of the ratios are shown, which are smoothed curves, added by the R function `geom_smooth()`. The calculation detail is as follows. For each cell type, suppose the target protocol has $m$ cells, and the reference protocol has $n$ cells. In each random partition, we first randomly select $\frac{\min(m,n)}{2}$ cells from the target protocol and the reference protocol each, and compute a gene's two means respectively. Second, for the remaining cells in the reference protocol, we randomly select $\frac{n}{2}$ cells and compute the same gene's mean as a reference. Third, we calculate the cross-protocol ratio and the within-protocol ratio by dividing the first two means by the reference mean. We repeat the above three steps for 100 times for every gene in each cell type and each target protocol. From the plot, we can see that, unlike the within-protocol ratios, which center around the constant value of 1, the cross-protocol ratios fluctuate between genes, and do not center around a constant value. This shows that there does not exist a single scaling factor to convert all genes' expression levels from one protocol to another.

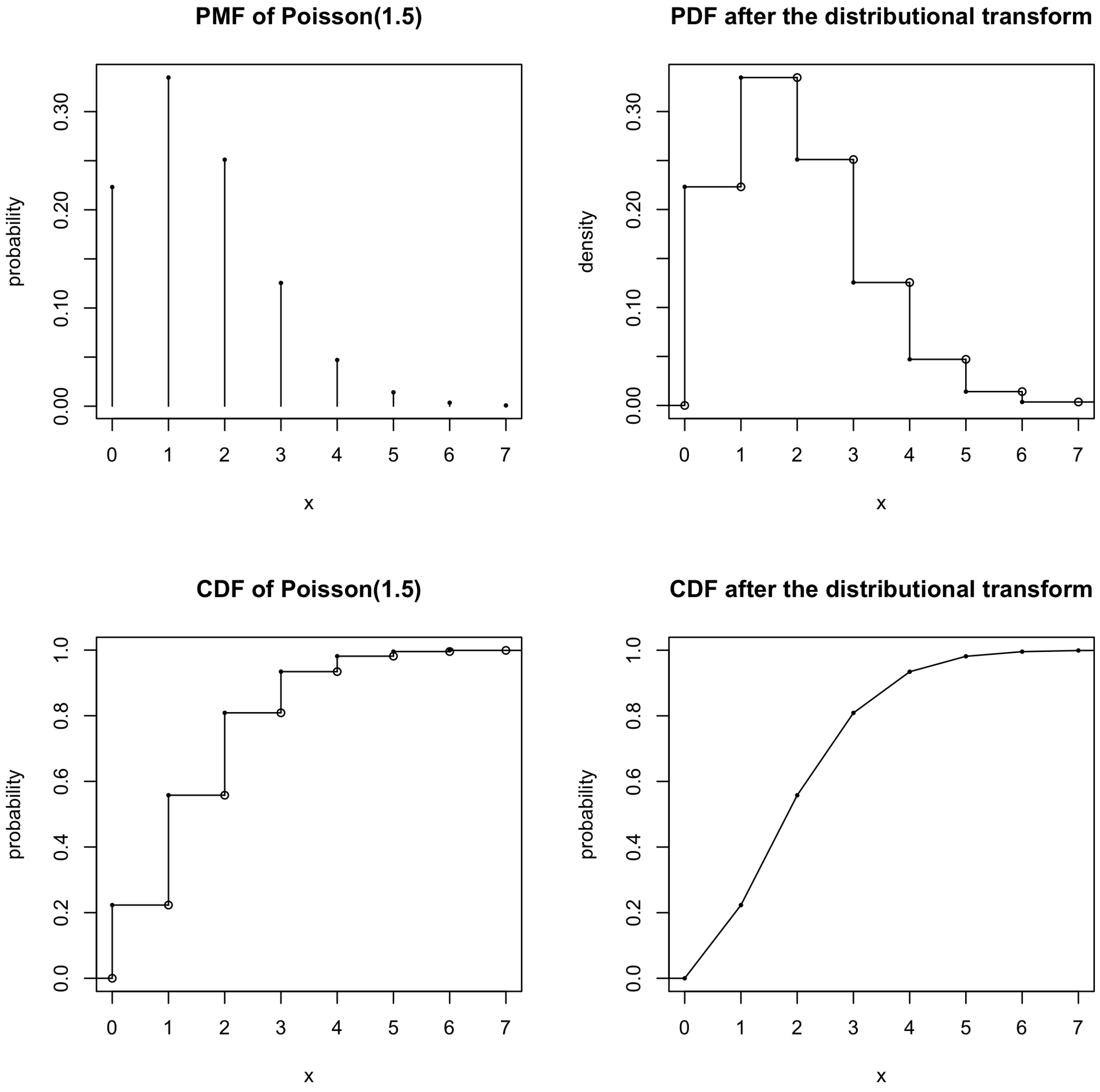


**Figure S47: A toy example showing the effect of the distributional transform.** For a discrete random variable $X$, the distributional transform maps its probability mass at each value $x$, which has a non-zero probability mass, uniformly to the interval $[x, x+1)$, thus converting $X$ to a continuous random variable. The top left and right panels show the probability mass function (PMF) before the transform and the probability density function (PDF) after the transform; the bottom left and right panels show the cumulative distribution functions (CDFs) before and after the transform.

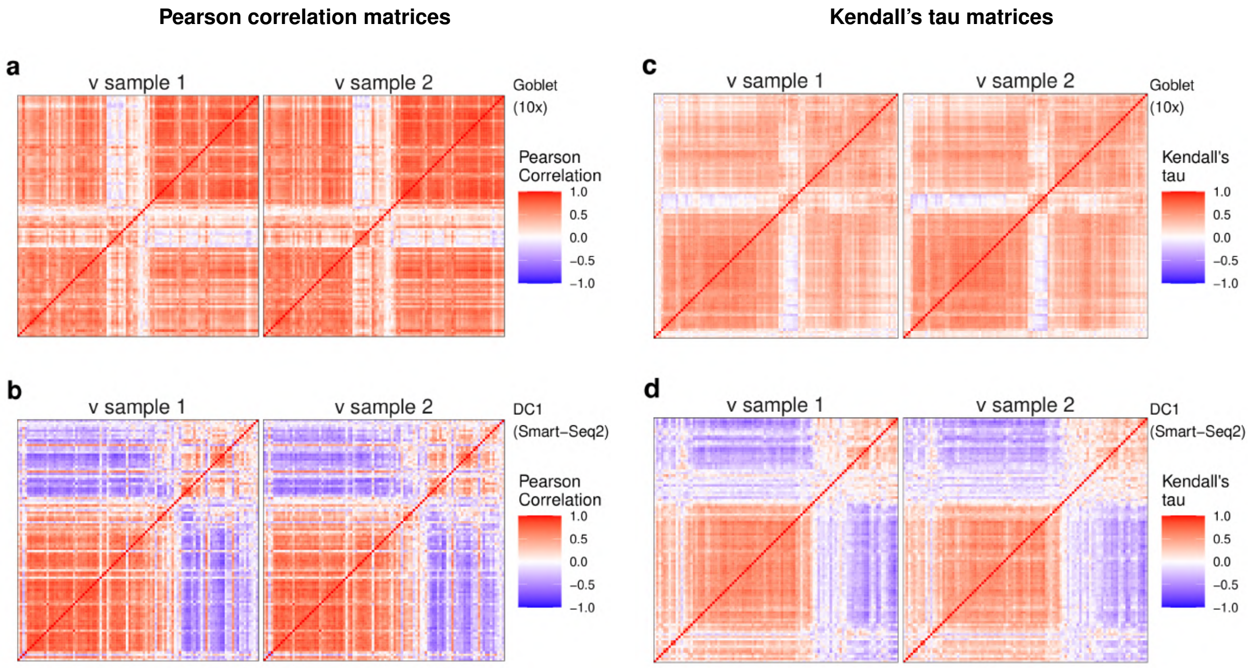


**Figure S48: Heatmaps of gene correlation matrices estimated from synthetic data generated by scDesign2, with $\widehat{\mathrm{R}}$ estimated under two different random samples of $v_{ij}^*$.** (a)-(b) Pearson correlation matrices; (c)-(d) Kendall's tau matrices. In (a) and (c), the scDesign2 model is fitted with goblet cells measured by 10x Genomics [76]; In (b) and (d), the scDesign2 model is fitted with cells of dendrocytes subtype 1 (DC1) measured by Smart-Seq2 [79]. For each cell type, the Pearson correlation matrices and Kendall's tau matrices are shown for the 100 genes with the highest mean expression values in the test data; the rows and columns (i.e., genes) of all the matrices are ordered by the complete-linkage hierarchical clustering of genes (using Pearson correlation as the similarity in (a)-(b) and Kendall's tau in (c)-(d)) in the test data. We find that the effect of the sampling of $v_{ij}^*$ on the estimated gene correlation matrices of the synthetic data is negligible, since the two matrices in each panel are very similar.
